# Thioether-functionalized cellulose for the fabrication of oxidation-responsive biomaterial coatings and films

**DOI:** 10.1101/2024.07.24.604973

**Authors:** Eric M. DuBois, Kate E. Herrema, Matthew G. Simkulet, Laboni F. Hassan, Payton R. O’Connor, Riya Sen, Timothy M. O’Shea

## Abstract

Biomaterial coatings and films can prevent premature failure and enhance performance of chronically implanted medical devices. However, current hydrophilic polymer coatings and films have significant drawbacks, including swelling and delamination. To address these issues, we modified hydroxyethyl cellulose with thioether groups to generate an oxidation-responsive polymer, HEC_MTP_. HEC_MTP_ readily dissolves in green solvents and can be fabricated as coatings or films with tunable thicknesses. HEC_MTP_ coatings effectively scavenge hydrogen peroxide, resulting in conversion of thioether groups to sulfoxide groups on the polymer chain. Oxidation-driven, hydrophobic-to-hydrophilic transitions that are isolated to the surface of HEC_MTP_ coating under physiologically relevant conditions increase wettability, decrease stiffness, and reduce protein adsorption to generate a non-fouling interface with minimal coating delamination or swelling. HEC_MTP_ can be used in diverse optical applications and permits oxidation-responsive, controlled drug release. HEC_MTP_ films are non-resorbable *in vivo* and evoke minimal foreign body responses. These results highlight the versatility of HEC_MTP_ and support its incorporation into chronically implanted medical devices.

## 1. Introduction

Chronically implanted medical devices, such as pacemakers^1^, catheters^2^, electrodes^3^, pumps^2,4^ and prostheses,^5,6^ are used widely for monitoring and treating disease. Despite successful clinical implementation, these devices can experience various problems that cause premature device failure.^7–11^ Biomaterials formulated as coatings, which adhere or bond to the device surface, or films, which can be press-fit as a separate material layer onto devices, can prevent premature failure of chronically implanted medical devices as well as enhance their functionality and performance.^12^ Employing hydrogel- and hydrophilic polymer-based coatings or films of polyethylene glycol (PEG),^13,14^ zwitterionic polymers (eg. polycarboxybetaine, polysulfobetaine),^13,15,16^ or other synthetic polymers^17,18^ improves long-term medical device performance by altering device surface chemistry, wettability, topography, and/or mechanical properties. Coating-or film-derived surface modifications mitigate molecular and cellular fouling and thrombosis, reduce foreign body response (FBR)-mediated inflammation, and minimize fibrotic encapsulation that would otherwise cause the formation of opaque, cell-dense, fibrotic scar tissue layers around uncoated devices.^12,19–22^ Biomaterial coatings and films can also reduce the mechanical stiffness mismatch at the implant-tissue interface to minimize the activation of mechanosensitive receptors,^19,20,22^ limit friction and wear,^23,24^ mitigate damage from device micromotion,^25^ and prevent corrosion and subsequent device degradation,^26^ thereby attenuating pro-inflammatory macrophage states and the fibroblast to myofibroblast transitions that promote capsular contracture. Naturally-derived biomolecules and extracellular matrix (ECM) protein-based coatings and films have also been incorporated as biomimetic substrates on devices to limit immune recognition and enhance device-tissue integration.^27–31^ Aside from regulating FBR and cell interactions, biomaterial coatings and films can be imparted with additional traits to improve long-term device performance including antimicrobial resistance to prevent biofilm formation, controlled local small molecule drug release, and localized gene or viral vector delivery.^12^

Despite conferring important functions to medical devices, currently available biomaterial coatings and films still have significant drawbacks. Hydrophilic polymer coatings implanted in a dehydrated state can swell greater than 3 times in size *in vivo*, leading to increased compressive tissue damage, increased impedance, coating delamination from the underlying device, and patient discomfort.^32–34^ Coatings and films applied as soft, hydrated polymers can also easily delaminate or shear off during implantation, leading to the exposure of the underlying surface of the implant, which can stimulate deleterious FBRs.^35^ Hydrophilic polymers delaminated from devices also pose an increased embolism risk and can disseminate and deposit at distant tissue sites away from the implant, leading to persistent, non-healing abscesses.^36^ Direct chemical bonding of hydrophilic polymers to the device surface can overcome delamination issues but requires specialized surface preparation that can not be applied to all devices.^13,37^ Natural ECM-based coatings and films suffer from sourcing and scaling issues, as well as increased immunogenic risk.^38–41^ To address these issues, materials engineering approaches are needed to generate new hydrophilic, non-fouling biomaterials that can be stably incorporated onto medical devices, so that they are resistant to disruption during implantation, persist chronically, and do not significantly swell *in vivo*. Additionally, beyond satisfying these general specifications, applications of biomaterial coatings and films could require other context-dependent properties, such as limited light scattering for optical devices, minimal impedance for electrical recording and stimulation, or the ability to elute drugs to direct local biological actions. Cost and manufacturability should also be considered for the effective scale up and clinical translation of coating and film technologies.

Cellulose-based materials have been used in industrial coating applications dating back to the late 1800’s, starting with water repellent, scratch resistant, nitrocellulose-based lacquers.^42^ Due to their natural abundance and renewable sourcing, coatings and films derived from cellulose esters have become ubiquitous in many diverse industrial applications including: dialysis and filtration membranes (regenerated cellulose),^43^ photography film and coatings on drug formulations to modify release kinetics (cellulose esters, including cellulose acetate and cellulose acetate butyrate),^44,45^ lacquer coatings (cellulose acetate butyrate),^42,46–49^ and tape (cellulose acetate).^50^ Across all these applications, chemical modification of the hydroxyls on the repeating anhydroglucose unit of cellulose confers emergent and modular properties, including the ability to be dissolved in organic solvents and increased hydrophobicity necessary to repel water.^51–55^ Cellulose-based materials have received only limited use as coatings or films on chronically implanted medical devices because hydrophilic derivatives swell, readily delaminate, and have challenging solubility profiles for fabrication.^53^ Although cellulose esters have improved solubility profiles in organic solvents, which allows them to form effective coatings or films, their poorer surface wettability compared to hydrophilic polymers causes increased protein and cell fouling that impacts overall biocompatibility.^56–58^ New cellulose derivatives that afford the processability of cellulose esters to generate effective coatings and films, while also facilitating the hydrophilic interface necessary for enhanced biocompatibility, would be broadly appealing for use with chronically implanted medical devices.

Here, we derived a new, smart, cellulose-based material that can be readily formulated as a coating or film, persists chronically *in vitro* and *in vivo*, and enables the generation of a soft, non-fouling, hydrophilic interface with minimal material swelling. To achieve this, we leveraged chemistry bioinspired by methionine-based materials^59^ to endow a specific cellulose polymer, hydroxyethyl cellulose (HEC), with thioether groups and confer oxidation-responsive properties. Upon oxidation under physiologically relevant conditions, this new thioether-functionalized cellulose readily transitions from a hydrophobic to a hydrophilic polymer via the conversion of the pendant thioethers to sulfoxide groups. We hypothesized that a thioether-functionalized, cellulose material would be soluble in organic solvents and thus processable as both coatings or films and that the exposed surface of these preparations would be readily oxidizable *in vivo* by physiological concentrations of reactive oxygen species, leading to the formation of a substrate that is more hydrophilic, non-fouling, and soft, which would confer enhanced biocompatibility. To test this hypothesis, we synthesized the thioether-functionalized cellulose polymer, HEC_MTP_, at large scales using a robust, single step isothiocyanate-based reaction, and investigated its ability to serve as an effective biomaterial coating and film. HEC_MTP_ readily dissolves in organic solvent systems, and coating outcomes, such as thickness and surface roughness, could be tuned by adjusting composition and fabrication parameters. Treatment of HEC_MTP_-coated substrates with physiological concentrations of hydrogen peroxide *in vitro* resulted in selective oxidation of thioether groups that served to increase surface hydrophilicity, decrease the effective surface mechanical stiffness, and reduce the extent of protein adsorption and neural cell fouling while causing negligible material swelling. We demonstrated the utility of HEC_MTP_ coatings for optical applications and HEC_MTP_ films for oxidation-responsive controlled release of molecules. HEC_MTP_ films persist *in vivo* for at least 4 weeks when implanted subcutaneously and show minimal immune cell recruitment and fibrotic scarring compared to equivalent non-oxidation responsive cellulose biomaterials. Our findings demonstrate the feasibility of using this oxidation-responsive, thioether-functionalized cellulose platform to improve the function and longevity of chronically implanted medical devices.

## 2. Results

### 2.1. Thioether-functionalized cellulose (HECMTP) is an oxidation-responsive polymer

We selected hydroxyethyl cellulose (HEC) as a template cellulose polymer for post-polymerization modification with thioether groups as it is highly abundant, inexpensive, has been previously used in medical applications, and is commercially available in numerous molecular weights. HEC contains accessible primary hydroxyls on the repeating anhydroglucose unit that can be used as nucleophilic species in isothiocyanate- and isocyanate-based reactions. HEC (Mw 430kDa (GPC), **Figure S1A**) was reacted with excess 3-(methylthio)propyl isothiocyanate in dimethyl sulfoxide (DMSO) with n,n-diisopropyl ethylamine (DIEA) as a catalyst to generate a new polymer, HEC_MTP_, which has 3-(methylthio)propyl (MTP) groups attached to the HEC polymer backbone by a stable thiocarbamoyl linker (**Figure 1A, Table S1**). HEC_MTP_ could be prepared with 5-23 mol% MTP functionalization relative to the available free primary hydroxyls, with moderate control of functionalization achieved through tuning reaction stoichiometry (**Figure 1B-C, Figure S1**). While HEC readily dissolved in water to form a viscous solution, HEC_MTP_ at functionalities as low as 9 mol% formed hydrated gels, and at functionalities greater than 15 mol% was insoluble in water (**Figure 1D, S1**). Based on an initial screen of different functionalizations, HEC_MTP_ with MTP functionalization at an average of 21 mol%, or ∼0.63 MTP groups per estimated anhydroglucose unit repeat, was selected for detailed evaluation. All studies reported here were completed using a single batch of HEC_MTP_ that was derived by blending the yields from 4 individual, 1 gram-scale reactions of near equivalent functionality (**Table S2, Figure S2B**).

**Figure 1:**
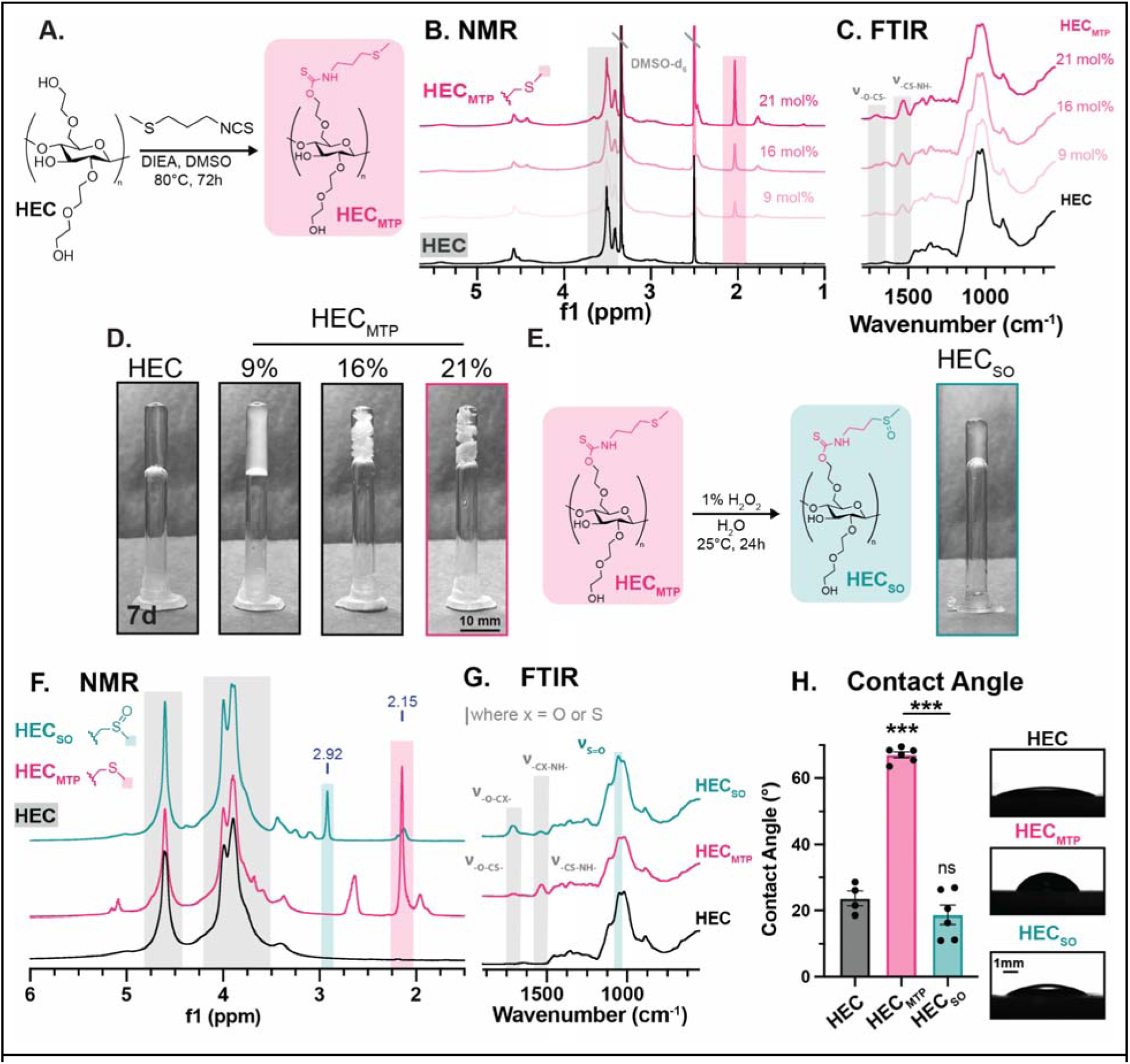
Characterization of HEC_MTP_ polymers demonstrates control of physicochemical properties. **(A)** Synthesis scheme for base-catalyzed, nucleophilic addition of MTPI to HEC to form HEC_MTP_. **(B)** ^1^H NMR of HEC and HEC_MTP_ at 9, 16, and 21mol% functionality in dDMSO. **(C)** FTIR of HEC and HEC_MTP_ at 9, 16, and 21mol% functionality highlighting MTP-associated peaks. **(D)** HEC and HEC_MTP_ at 9, 16, and 21mol% solubilized at 5wt% in PBS. **(E)** HEC_MTP_ can be oxidized to form HEC_SO_ via the application of 1% H_2_O_2_ in water for 24hr. Resulting HEC_SO_ solubilized at 5wt% in PBS. **(F)** ^1^H NMR in dTFA highlighting characteristic peaks associated with HEC, HEC_MTP_ and HEC_SO_. Labels in blue demarcate peak location. **(G)** FTIR with characteristic absorption bands of chemical bonds in HEC, HEC_MTP_ and HEC_SO_. **(H)** Contact angle data demonstrating a significant increase in hydrophobicity on the addition of the thioether moiety to HEC, and a subsequent, significant decrease in hydrophobicity upon oxidation of the thioether moiety (HEC_SO_). Inset: brightfield images from the goniometer demonstrating the deposition of deionized (DI) water on HEC, HEC_MTP_ and HEC_SO_ films. Not significant (ns) and ***P < 0.0001 compared to HEC or as designated, one-way ANOVA with Tukey’s multiple comparison test. Graph shows mean ± s.e.m. with individual data points showing n = 4 for HEC, n = 6 for HEC_MTP_ and HEC_SO_.

To determine whether thioether groups on HEC_MTP_ could be oxidized to generate the sulfoxide- functionalized polymer, designated as HEC_SO_, we exposed HEC_MTP_ to oxidizing conditions for one day using 1% (327mM) hydrogen peroxide (H_2_O_2_) (**Figure 1E**). Conversion of thioether groups to sulfoxide groups was detected by ^1^H NMR as a readily identifiable shift in the thioether methyl protons from 2.15 ppm to 2.92 ppm (**Figure 1F**).^60^ Using 1% H_2_O_2_, all thioether groups were converted to sulfoxide groups, verified by complete loss of the thioether methyl protons at 2.15 ppm (**Figure 1F, S2**). By FTIR, HEC_MTP_ showed emergent -CS-NH- and -O-CS- (thiocarbamoyl) stretches at 1535 cm^-1^ and 1710 cm^-1^ respectively following MTP addition,^61,62^ while oxidation of HEC_MTP_ to HEC_SO_ showed characteristic emergence of a S=O stretch at 1055 cm^-1^ (**Figure 1G**).^63^ Under the 1% H_2_O_2_ oxidation conditions, the FTIR spectra within the thiocarbamoyl region showed a notable increase in the intensity of the 1710 cm^-1^ stretch and a concurrent reduction in the 1535 cm^-1^ stretch which is consistent with the sulfur atom in the thiocarbamoyl being substituted by an oxygen atom to form -O-CO-NH- (**Figure 1G**).^61^ Despite these changes, there was preserved total integration of the two stretches within the thiocarbamoyl region (**Figure S3A-B**).

Following oxidation, there was a reduction in the total integrated area of methyl protons by NMR in HEC_SO_ samples compared to unmodified HEC_MTP_. We suspected that the attenuated total detected sulfoxide-associated methyl protons on HEC_SO_ by NMR was a result of significantly reduced solubility in dTFA (deuterated trifluoroacetic acid) as well as other common NMR solvents, whereas HEC and HEC_MTP_ dissolved completely in dTFA and returned clean NMR spectra. Oxidizing HEC_MTP_ with lower concentrations of H_2_O_2_ within the physiological range (80-320µM) for one day resulted in partial oxidation of the polymer, such that it was still completely soluble in dTFA for NMR evaluation. Under these dilute oxidizing conditions, we detected partial conversion of thioether groups to sulfoxides and preservation of methyl protons by NMR, implying no cleavage of functionalized groups from the HEC backbone (**Figure S3C**). Evaluation of HEC_MTP_ samples before and after oxidation by X-ray photoelectron spectroscopy (XPS) also corroborated the preservation of sulfur content on the polymer after oxidation (**Fig S4**). We did not detect any notable changes to stretches in the thiocarbamoyl region by FTIR in these samples implying preferential thioether oxidation over thiocarbamoyl oxidation upon exposure to physiological concentrations of H_2_O_2_ (**Fig S5**). The template polymer, HEC, was not altered following one day of 1% H_2_O_2_ exposure, which indicated that there was no detectable oxidation of ethylene oxide groups on the polymer backbone to which MTP groups were linked (**Figure S6**). Collectively, these data demonstrate that there was negligible cleavage of the MTP group from the HEC polymer backbone after oxidation and preferential oxidation of the thioether moiety, consistent with previous reports using similar chemistries.^64,65^ Characterization of HEC_MTP_ by differential scanning calorimetry (DSC) and thermal gravimetric analysis (TGA) showed that HEC_MTP_ was thermally stable up to 215°C, making it amenable to a wide range of processing conditions and methods (**Figure S7**).

To assess differences in wettability between HEC variants, we performed sessile-drop goniometry on casted films of HEC, HEC_MTP_, and HEC_SO_ to evaluate relative hydrophilic-hydrophobic character. As anticipated, MTP functionalization resulted in a significant decrease in wettability, with HEC_MTP_ displaying a contact angle of 67° which was more than double that of the unmodified HEC (24°) and within the range of reported values for other cellulose ester benchmarks (**Figure 1H**).^66^ Oxidation of HEC_MTP_ to HEC_SO_ yielded a significant 48° decrease in contact angle, resulting in a surface that had equivalent wettability to HEC, demonstrating that oxidation with 1% H_2_O_2_ is an effective chemical stimulus to substantially increase the hydrophilicity of this thioether-functionalized cellulose by converting thioether functional groups to sulfoxides (**Figure 1H**).

Collectively, these data show that HEC can be readily modified with thioether functionalities at high scale and with stoichiometric control to generate a new, oxidation responsive polymer, HEC_MTP_. Oxidation of HEC_MTP_ prompted sulfoxide conversion but no pendant group cleavage and was sufficient to transition HEC_MTP_ from a water insoluble polymer to a hydrophilic polymer. Since HEC_MTP_ possessed the unique capacity to be processed using organic solvents like a cellulose ester but also to selectively transition into a hydrophilic interface via a ubiquitous biological stimulus, we were prompted to explore in earnest its potential as a coating and film for biomedical applications.

### 2.2. HEC_MTP_ can be fabricated as a coating or film

Dip coating is a standard industrial technique for processing coatings of cellulose esters with coating properties, including thickness and surface finish, readily altered by key formulation parameters such as solvent system and temperature.^67–69^ To determine an appropriate solvent system in which HEC_MTP_ could be solubilized for coating applications, we prepared HEC_MTP_ at 10mg/mL in common green organic solvents including DMSO, methanol, ethanol, isopropyl alcohol (IPA), acetone, and water. At this relatively dilute concentration, HEC_MTP_ was soluble in DMSO but was insoluble, and did not disperse, in any of the other solvents (**Figure 2A**). To identify a wider range of useful solvent systems for preparing HEC_MTP_, we generated and tested blends of DMSO with the same array of green organic solvents at 50%/50% v/v composition. HEC_MTP_ was soluble in all DMSO blended solvent systems except for DMSO/water in which an insoluble, dispersed polymer phase was noted (**Figure 2B**).

**Figure 2:**
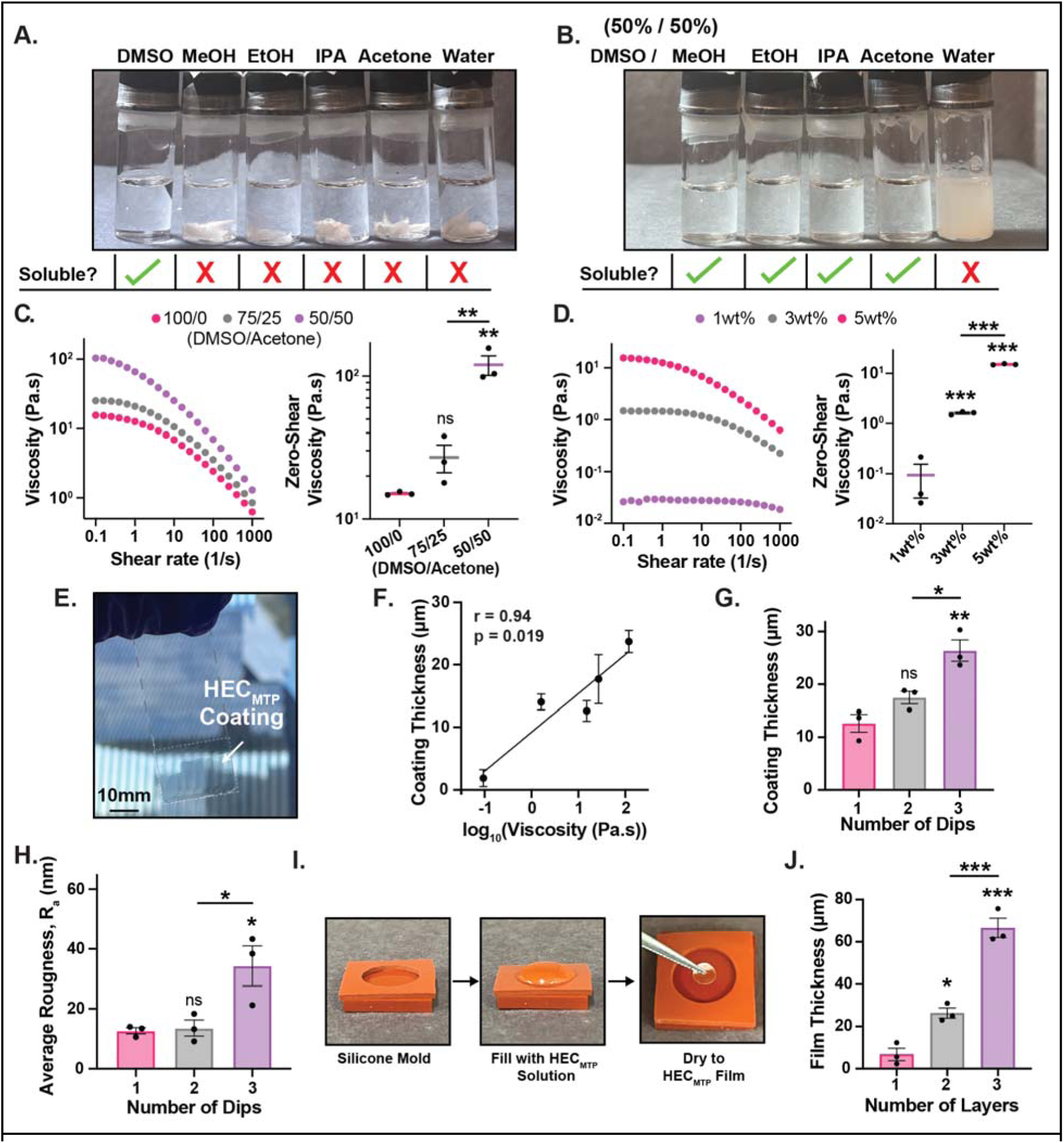
HEC_MTP_ polymer can be formulated as coatings or films with tunable thickness. **(A)** Solubility table showing images of HEC_MTP_ polymer solubilized at 10mg/mL in DMSO and insoluble in MeOH, EtOH, IPA, acetone and water. **(B)** Solubility table showing images of HEC_MTP_ polymer solubilized at 10mg/mL in a 50%/50% solvent system of DMSO and MeOH, EtOH, IPA, or acetone, but precipitated in DMSO/water (50%/50%). **(C)** *Left:* Dynamic rheological shear rate sweep measurements for HEC_MTP_ solubilized at 5wt% in a 100/0, 75/25, 50/50 DMSO/acetone solvent system showing viscosity (Pa.s) against increasing shear rate (s^-1^). *Right:* Box-and-whisker plot showing the increase in zero-shear viscosity (Pa.s), measured at a shear rate of 0.1 (s^-1^), with increasing concentration of acetone. Not significant (ns) and **P < 0.003 compared to 100/0 DMSO/Acetone or as designated. **(D)** *Left:* Dynamic rheological shear rate sweep measurements for HEC_MTP_ at 1, 3, and 5wt% in DMSO showing viscosity (Pa.s) against increasing shear rate (s^-1^). *Right:* Box-and-whisker plot showing the increase in zero-shear viscosity (Pa.s), measured at a shear rate of 0.1 (s^-1^), with increasing concentration. ***P ≤ 0.001 compared to 1wt% or as designated. **(E)** Brightfield image showing a HEC_MTP_ coating on a glass substrate. Black dashed lines outline the glass slide. White dashed lines outline the HEC_MTP_-coated portion of the glass slide. **(F)** Semi-log scatter plot showing the relationship between viscosity and coating thickness. Graph shows mean viscosity vs. mean coating thickness ± s.e.m. with individual data points showing n = 3 per group. The semi-log data line is fit with a linear line of best fit and the correlation is reported as Pearson’s Correlation Coefficient (r). **(G)** Bar graph showing effect of multiple dips of HEC_MTP_ solubilized at 5wt% in 100/0 DMSO/Acetone on coating thickness. *P < 0.03, **P <0.003, and not significant (ns) compared to 1 dip or as designated. **(H)** Bar graph showing the effect of multiple dips on the average roughness of the coating. *P < 0.04, and not significant (ns) compared to 1 dip or as designated. **(I)** Brightfield images showing the film-making process, including: the silicone mold, deposition of soluble HEC_MTP_, and the resulting dried and punched HEC_MTP_ film. **(J)** Bar graph showing effect of multiple layers on film thickness using a 3wt% HEC_MTP_ solution in DMSO. *P < 0.02, ***P <0.0005 compared to 1 layer or as designated. All statistical comparisons were made using one-way ANOVAs with Tukey’s multiple comparison tests. Graphs show mean ± s.e.m. with individual data points showing n = 3 per group.

Viscosity of cellulose solutions influences the thickness and regularity of finished coatings, as well as its workability during processing.^67–69^ Therefore, we next used rheological testing to assess the effect of solvent system, polymer concentration, and temperature on the viscosity of HEC_MTP_ solutions. The composition of DMSO/organic solvent blends impacted HEC_MTP_ solution viscosity, with increasing proportions of acetone in a DMSO/acetone blend resulting in a dramatic increase in the viscosity of 5wt% HEC_MTP_ solutions, ranging from 15.1 Pa.s at 0% acetone to 119.8 Pa.s with 50% acetone incorporation (**Figure 2C**). Increasing the concentration of HEC_MTP_ in DMSO-only formulations from 1wt% to 3wt% also caused a significant increase in viscosity from 0.09 to 1.6 Pa.s, while the viscosity of 5wt% preparations increased by an order of magnitude to 15.1 Pa.s (**Figure 2D**). There was a notable effect of temperature on HEC_MTP_ solution viscosity and we observed a significant, linear decrease in viscosity from 10 Pa.s at 30°C to 2.0 Pa.s at 70°C (**Figure S8A-B**). Taken together, these data suggest that HEC_MTP_ solution viscosity can be readily tuned across 3 orders of magnitude by adjusting polymer concentration, solvent system, and temperature at processing.

We next tested how HEC_MTP_ solution viscosity influences the thickness and roughness of prepared coatings. We used acid-cleaned glass slides and dipped HEC_MTP_ solutions at a fixed rate of 1 mm/sec (the fastest rate that still approximately maintains zero-shear viscosity or a shear rate of 1 sec^-1^) and allowed slides to dry in a vacuum oven at 40°C for at least 72 hours to remove residual DMSO solvent and form the coating (**Figure 2E**). We identified differences in the thickness of HEC_MTP_ coatings prepared from different DMSO/acetone blends at 5wt% HEC_MTP_ with all formulations resulting in coatings with a thickness ranging from 10-25μm (**Figure 2F, S8C-D**). There was a notable change in surface roughness due to solvent blending, with 50% DMSO/acetone solutions demonstrating large, undulating surface roughness on the micron scale and a significantly higher average roughness (∼29.6nm) that was more than double that of the 100% DMSO solvent system (12.7nm) (**Figure S9A, S9D**). To test the effect of polymer solution concentration on coating thickness and roughness, we prepared dip coated slides at 1, 3, and 5wt% in DMSO only. Increasing the solution’s polymer concentration significantly increased the coating thickness from 1.9µm at 1wt% to 14.1 µm at 3wt%, but there was minimal increase in coating thickness between the 3wt% and 5wt% preparations (**Figure S8D**). Surface profilometry of these coatings revealed that there was a large increase in roughness from 1wt% to 3wt%, but an apparent smoothing at 5wt% (**Figure S9B, S9E**), suggesting that the higher wt% coating may deposit a significantly greater total amount of HEC_MTP_, effectively creating a smoother and more uniform coating. Scanning electron microscopy (SEM) micrographs of HEC_MTP_ coatings applied to aluminum stubs mirrored this trend. While 3wt% HEC_MTP_ preparations had essentially featureless surfaces (**Figure S9G**), 1wt% HEC_MTP_ coatings showed regular, dimple-like features approximately 2µm in diameter, suggesting that this lower HEC_MTP_ concentration was insufficient to cleanly coat the substrate (**Figure S9H**). These data demonstrate that HEC_MTP_ coatings, when prepared at appropriate concentrations, could effectively smooth the surface of a roughened underlying substrate. When comparing all the different formulations together, HEC_MTP_ coating thickness was directly proportional to the measured magnitude of viscosity of the coating solution irrespective of solvent system or concentration (**Figure 2F**). We also evaluated the effects of fabricating multilayer coatings by dipping a substrate in a solution of 5wt% HEC_MTP_ in DMSO and incorporating a 24-hour drying step between subsequent dips. Increasing from 1 to 3 dips resulted in a significant increase in coating thickness from 12.6 to 26.4µm (**Figure 2G**). Evaluating the average roughness, we observed a small, but significant increase from 1 dip at 12.7nm to 3 dips at 34.3nm (**Figure 2H, S9C, S9F**).

To form free-standing films from HEC_MTP_, we first fabricated a silicone mold composed of two, press-fit silicone sheets (**Figure 2I)**. HEC_MTP_ in DMSO was pipetted to fill the silicone mold cavity and dried in a vacuum oven at 40°C to form a large free-standing film. Individual wafers of defined geometries could be punched out from larger film preparations using a biopsy punch, enabling a rapid and scalable approach to film fabrication (**Figure 2I**). By depositing a layer of HEC_MTP_ solution every 24 to 48 hours we could tune film thickness from 6.8µm (1 layer) to 66.6 µm (3 layers) with no evidence of lamination defects within multilayered constructs (**Figure 2J**). These data show that HEC_MTP_ can be dissolved in green organic solvents and processed to form coatings and films whose thickness and roughness can be readily tuned through careful selection of solvent and processing parameters.

### 2.3. HEC_MTP_ coatings are oxidation-responsive and stable under simulated physiological conditions

Next, we evaluated the physical and oxidation-responsive properties of HEC_MTP_ when formulated as a coating. To elucidate structure-function correlates for HEC_MTP_ coatings, we synthesized two other HEC derivatives to serve as benchmarking controls: (i) HEC_HEX_, which contains a thiocarbamoyl linked aliphatic hexyl group of similar length to that on HEC_MTP_ but without the thioether functionality, and (ii) HEC_OCT_, which contains an aliphatic octyl group linked to the polymer backbone by a carbamate linker (**Figure S10-11, Table S3-4**). HEC_HEX_ and HEC_OCT_ were appropriate functional controls for evaluating the physical and oxidation- responsive properties of HEC_MTP_ as they were synthesized via equivalent chemistries (isothiocyanate and isocyanate reactions respectively), had similar hydrophobicity to the MTP group, and both lacked thioether functionality. However, important limitations with the fabrication of HEC_HEX_ and HEC_OCT_ benchmarking controls warrant brief review here. Due to the limited reactivity of hexyl isothiocyanate, HEC_HEX_ could only be functionalized to approximately 5 mol% hexyl groups on the polymer chain, which was considerably less than the 21 mol% functionalization that could be readily achieved for HEC_MTP_. HEC_OCT_, by virtue of employing more reactive isocyanate rather than isothiocyanate chemistries in its synthesis, could be readily tuned up to 100 mol% functionality. However, at functionalities of 12 mol% and higher, we observed issues with HEC_OCT_ solubility whereby the polymer would either be insoluble in DMSO or would solubilize with mild heating but rapidly crystallize as it cooled (**Figure S12**). In an attempt to address this issue, we tested 18mol% HEC_OCT_ in a variety of different solvent formulations, including: water, acetone, toluene, and 50/50 DMSO/toluene (**Figure S13**). HEC_OCT_ at 18 mol% octyl functionalization formed swollen hydrogels in water, was insoluble in acetone and toluene, and was only partially soluble in 50/50 DMSO/toluene, preventing its effective formulation as a free-standing film or coatings. Therefore, with consideration of the above limitations, HEC_HEX_ at 5 mol% hexyl and HEC_OCT_ at 8 mol% octyl groups were used to fabricate coatings for comparisons with HEC_MTP_ coatings.

As a first test of the oxidation-responsive properties of HEC_MTP_ coatings, we drop-casted HEC_MTP_ and benchmarking polymers into wells of a 96 well plate to form thin coatings that were incubated overnight in 100µL of 4mM hydrogen peroxide at 37, which equated to an approximate 2:1 molar ratio of MTP moieties to H_2_O_2_. At 24 hours, HEC_MTP_ coatings had scavenged 82.8% (or 1046.5 µmol/cm^2^) of the available H_2_O_2_ whereas HEC and HEC_OCT_ coatings did not scavenge any detectable amount of H_2_O_2_ (**Figure 3A**). HEC_HEX_ scavenged a small but significant amount of incubated H_2_O_2_ (15.6%), likely through oxidation of the thiocarbamoyl linker as noted earlier (**Figure 3A**). FTIR-based comparisons of HEC, HEC_HEX_, HEC_OCT_, and HEC_MTP_ before and after oxidation with 1% H_2_O_2_ confirmed that both HEC_HEX_ and HEC_MTP_ had H_2_O_2_-induced changes in the two stretches in the thiocarbamoyl region consistent with the sulfur atom on the thiocarbamoyl being substituted for by an oxygen atom^61^ (**Figure S14**). HEC and HEC_OCT_ showed no apparent change in their respective FTIR spectra after oxidation which further strengthened our confidence in the assessment that the thiocarbamoyl was altered in the HEC_HEX_ and HEC_MTP_ polymers under these millimolar-range H_2_O_2_ conditions.

**Figure 3:**
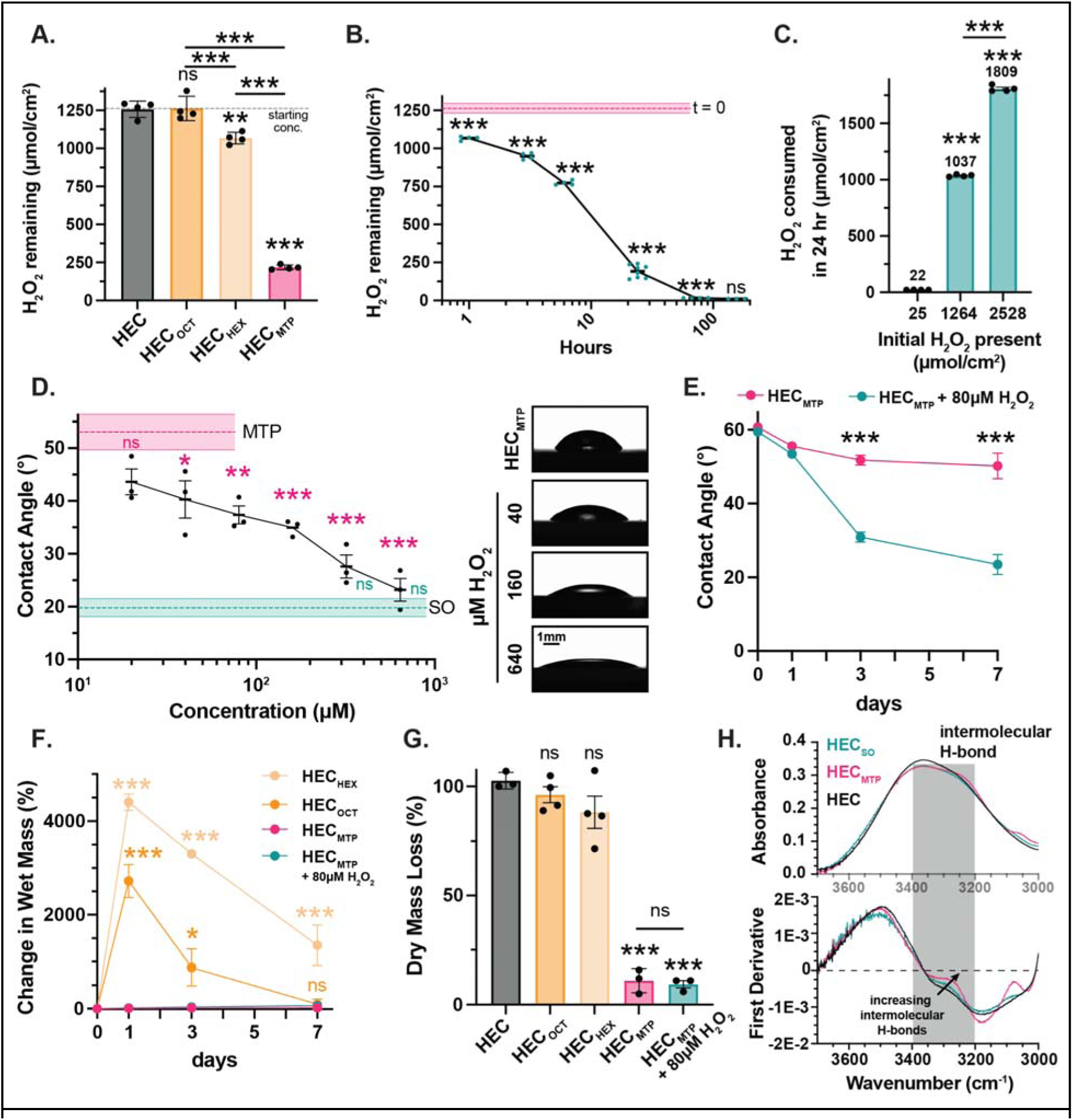
HEC_MTP_ coatings oxidize in response to H_2_O_2_ exposure, but are impervious to dissolution in aqueous buffers. **(A)** H_2_O_2_ remaining (µmol/cm^2^) following exposure of HEC, HEC_OCT_, HEC_HEX_, and HEC_MTP_ coatings to 4mM H_2_O_2_ for 1d. **P <0.0015, ***P ≤ 0.001, and not significant (ns) compared to HEC or as designated, one-way ANOVA with Tukey’s multiple comparison test. Graph shows mean ± s.e.m. with individual data points showing n = 4 per group. **(B)** H_2_O_2_ remaining (µmol/cm^2^) after exposure of HEC_MTP_ coating to 4mM H_2_O_2_ for 1-168 hours (7d). ***P≤ 0.0007, each timepoint is compared with the previous timepoint, one-way ANOVA with Tukey’s multiple comparison test. Graph shows mean ± s.e.m. with individual data points showing n = 8 for 24hrs, and n = 4 for all other groups. The line for t = 0 shows mean ± s.e.m., with s.e.m. represented as light shaded areas. **(C)** µmol/cm^2^ H_2_O_2_ consumed after exposure of HEC_MTP_ coatings to 100µL of H_2_O_2_ at variable concentrations for 1d. ***P< 0.0001 compared to 25 µmol/cm^2^ H_2_O_2_ or as designated, one-way ANOVA with Tukey’s multiple comparison test. Graph shows mean ± s.e.m. with individual data points showing n = 4 per group. **(D)** Exposure to increasing concentrations of H_2_O_2_ for 24 hrs leads to significant decreases in contact angle. *P < 0.03, **P ≤0.004, ***P <0.001, and not significant (ns) with comparisons in pink being made to HEC_MTP_ (MTP) and comparisons in turquoise being made to HEC_SO_ (SO), one- way ANOVA with Tukey’s multiple comparison test. Graph shows mean ± s.e.m. with individual data points showing n = 6 for SO and n = 3 per group for all other conditions. Inset: brightfield images from the goniometer demonstrating the deposition of deionized water on HEC_MTP_ and HEC_MTP_ coatings treated with 40, 160, and 640µM of H_2_O_2_. **(E)** Contact angle measurements on HEC_MTP_ coatings over 7d exposed to either PBS or PBS + 80µM H_2_O_2_. ***P< 0.0001 comparisons between HEC_MTP_ and HEC_MTP_ + 80µM H_2_O_2_ at equivalent timepoints, two-way ANOVA with Tukey’s multiple comparison test. Graph shows mean ± s.e.m., n = 3 per group. **(F)** Change in wet mass of HEC_HEX_, HEC_OCT_ or HEC_MTP_ coatings over 7d exposed to either PBS or PBS + 80µM H_2_O_2_. *P< 0.04, ***P <0.0005, and not significant (ns) comparisons are made between HEC_HEX_ or HEC_OCT_ with HEC_MTP_ at equivalent timepoints, two-way ANOVA with Tukey’s multiple comparison test. Graph shows mean ± s.e.m., n = 4 for HEC_HEX_ and HEC_OCT_ and n = 3 for all other groups. **(G)** %Change in dry mass of HEC_HEX_, HEC_OCT_ or HEC_MTP_ coatings incubated in PBS or PBS + 80µM H_2_O_2_ over 7d. ***P< 0.0001, and not significant (ns) compared to HEC or as designated, one-way ANOVA with Tukey’s multiple comparison test. Graph shows mean ± s.e.m. with individual data points showing n = 4 for HEC_HEX_ and HEC_OCT_ and n = 3 for all other groups. **(H)** FTIR and first derivative of FTIR of HEC, HEC_MTP_ and HEC_SO_ films in -OH region showing increase in intermolecular H-bonds. First derivative was smoothed (0th order, 200 neighbors) and inverted.

To profile the kinetics of HEC_MTP_ oxidation we assessed the amount of H_2_O_2_ scavenged at defined time points from 1 hour to 7 days (**Figure 3B**). H_2_O_2_ scavenging by HEC_MTP_ demonstrated second order rate kinetics resulting in a sigmoidal time dependency, which was consistent with an autocatalytic process that has previously been observed in polysulfide-based materials^70^ (**Figure S15A**), with nearly all H_2_O_2_ (>99%) consumed following one week of incubation. HEC_MTP_ coatings scavenged H_2_O_2_ in a concentration dependent manner which was directly proportional to the initial concentration of applied H_2_O_2_ within the concentration range of 80µM to 8mM (**Figure 3C**). To further dissect the concentration and load dependent effects, we exposed HEC_MTP_ coatings to either 100µL of H_2_O_2_ at 4mM or 200µL of H_2_O_2_ at 2mM to evaluate different concentrations but conserved total molar equivalents of H_2_O_2_. After 24 hours, the HEC_MTP_ coatings had scavenged 82% of the 4mM H_2_O_2_ but only 55% of the 2mM H_2_O_2_, demonstrating the strong dependency of H_2_O_2_ concentration on the scavenging rate of the film (**Figure S15B**). We also noted a finite H_2_O_2_ scavenging capacity for HEC_MTP_ coatings that affected scavenging kinetics, such that high initial H_2_O_2_ loads (8mM) reduced the effective scavenging rate of the coating upon subsequent exposure to a second 4mM H_2_O_2_ solution (**Figure S15C**). By contrast, HEC_MTP_ coatings exposed to a 100-fold lower initial H_2_O_2_ load (80µM) could more effectively scavenge the 4mM H_2_O_2_ solution upon subsequent exposure (**Figure S15C**).

At sites of tissue injury and cell death *in vivo*, reactive oxygen species (ROS) are generated at concentrations ranging from 1-100µM, with exposure to 80-100µM ROS capable of causing oxidative stress-mediated cell necrosis or apoptosis.^71,72^ We assessed how H_2_O_2_ scavenging by HEC_MTP_ coatings within such physiological ROS concentrations altered the surface wettability of the coating. Exposing HEC_MTP_ coatings to H_2_O_2_ concentrations at or greater than 40µM for one day, caused a significant decrease in contact angle with 320µM H_2_O_2_ incubation resulting in contact angles indistinguishable from HEC_SO_, which defined the maximum wettability of the surface (**Figure 3D**). To assess whether the maximum wettability of the surface could be achieved by prolonged exposure to lower, physiologically relevant concentrations of H_2_O_2_, we incubated HEC_MTP_ coatings in 80µM H_2_O_2_ for 7 days. Under these milder oxidizing conditions, HEC_MTP_ coatings gradually transitioned to be maximally wettable over the course of 7 days with the greatest rate of change occurring from 1 to 3 days of incubation (**Figure 3E**). HEC_MTP_ incubated in PBS only showed a small change in contact angle over the 7 day incubation period indicative of a low amount of hydration of the polymer coating, but such effects on surface wettability paled in comparison to those induced by the oxidizing conditions (**Figure 3E**). Contact angle measurements were paired with bulk physical evaluations including coating swelling and dry mass loss (**Figure 3F-G, S16**). The wet mass of HEC_MTP_ coatings changed minimally over the course of 7 days when incubated in PBS only, while the gradual hydrophobic to hydrophilic surface transition provoked by oxidation with 80µM H_2_O_2_ caused a concurrent increase in coating wet mass that reached 67.7% by 7 days (**Figure 3F, S16**). This change in material wet mass was small and insignificant when compared to the HEC_HEX_ and HEC_OCT_ coatings, which swelled considerably with an increase in wet mass of over 4400% and 2700% respectively (**Figure 3F**). Over the course of 7 days, HEC_HEX_ coatings slowly dissolved, while HEC_OCT_ coatings delaminated and fragmented, resulting in both coatings subsequently decreasing in both wet and dry mass (**Figure 3F-G**). By contrast, HEC_MTP_ coatings persisted with minimal polymer resorption over the 7 days, with incubation in either PBS or 80µM H_2_O_2_ resulting in less than 12% total dry mass loss (**Figure 3G**). We hypothesized that the persistence and minimal swelling of HEC_MTP_ coatings could be due to strong physical bonding between polymer chains, which is characteristic of certain cellulose ester formulations.^73^ Reexamining the FTIR for HEC_MTP_, we noted a prominent shoulder in the spectra within the 3200-3300 cm^-1^ region which can be attributed to enhanced intermolecular hydrogen bonding (**Figure 3H**).^73,74^ This hydrogen bonding phenomena was present only in HEC_MTP_ and was not identified in HEC, HEC_SO_, or any of the other benchmarking materials. To confirm the presence of intermolecular hydrogen bonding, we assessed whether we could re-dissolve dried HEC_MTP_ films in 1,1,1,3,3,3- hexafluoro-2-propanol (HFIP), a known hydrogen bond disrupting solvent.^75,76^ While HEC_MTP_ films swell to nearly 4000% of their original dry mass but do not dissolve after 48 hour incubation in DMSO, HFIP effectively dissolved films in their entirety over the same time course, confirming the presence of strong hydrogen bonding within the dry HEC_MTP_ material (**Figure S17**).

These data show that HEC_MTP_, when formulated as a coating, readily scavenges physiologically relevant concentrations of H_2_O_2_ with second order reaction kinetics that are dependent on both the H_2_O_2_ and thioether concentrations. Oxidation of HEC_MTP_ coatings under physiologically relevant conditions achieves the maximal wettability of the surface over the course of several days and is associated with a small increase in wet mass as the coating hydrates. Despite hydration at the coating surface, strong intermolecular hydrogen bonding between polymer chains comprising the bulk of the HEC_MTP_ coating results in minimal total swelling and negligible mass loss to ensure long-term preservation of coating integrity.

### 2.4. HEC_MTP_ coatings and films show oxidation-driven changes in mechanical properties

Polymer coatings or films that effectively reduce the cell-perceived mechanical stiffness of the underlying medical device may promote improved biocompatibility. As such, there has been recent interest in engineering mechanically adaptable materials which are capable of *in situ* softening once implanted *in vivo* to further improve biocompatibility outcomes.^77^ We hypothesized that the oxidation-induced increase in surface hydrophilicity and hydration observed for HEC_MTP_ coatings under physiologically relevant conditions may also confer a concurrent reduction in mechanical stiffness, making it an effective mechanically adaptable material. To test this premise, we assessed the mechanical properties of HEC_MTP_ coatings in a number of different ways and benchmarked results against HEC and a crosslinked ethoxylated polyol (CEP)-based control (**Table S5**). We first conducted lap shear testing to evaluate polymer coating adhesion and interfacial shear strength in the dry state on a glass substrate. HEC_MTP_ coatings displayed a higher ultimate interfacial shear strength (2782 kPa) at failure relative to HEC (1158 kPa) and CEP (376 kPa) coating benchmarks (**Figure 4A-B, S18**). The observed adhesive strength of HEC_MTP_ on glass in the dry state makes it comparable to the performance of commonly used n-butyl-2-cyanoacrylate -based surgical glues under equivalent testing parameters.^78^ Both HEC_MTP_ and HEC coatings failed in lap shear tests by a combination of cohesive and adhesive failure whereas the CEP-based interface failed by a purely adhesive failure mechanism (**Figure S18**). The strong glass substrate adhesive properties of HEC_MTP_ coatings were maintained after 7 days of incubation in PBS and we observed minimal change in coating adhesion under mild oxidizing conditions (80µM H_2_O_2_). However, oxidation at supraphysiological concentrations of H_2_O_2_ (>640µM) for just one day caused HEC_MTP_ coatings to readily delaminate due to widespread conversion of polymer thioether groups to sulfoxides, which ultimately results in bulk material changes that are not seen under physiological oxidizing conditions. Next, we evaluated HEC_MTP_ films by tensile testing to assess how different oxidizing conditions alter bulk mechanical properties. HEC_MTP_ films in the dry state behaved as tough plastics with an elastic modulus of 455 MPa but softened significantly when equilibrated for 7 days in PBS such that the elastic modulus decreased by almost 3 orders of magnitude to 597 kPa and the hydrated material displayed behavior characteristic of a thermoplastic elastomer (**Figure 4C-D, S19**). Hydration alone reduced the ultimate tensile strength of HEC_MTP_ by over an order of magnitude, from 30.1 MPa to 894 kPa. However, engineering strain at failure increased nearly three-fold, from 0.53 to 1.38 (**Figure S19**). Oxidation of HEC_MTP_ films with 80µM H_2_O_2_ for 7 days caused no significant change in the elastic modulus, ultimate tensile strength (UTS), or strain to failure compared to the PBS incubated sample, indicating that conditions sufficient to oxidize the surface of the film provoked minimal bulk material changes. Exposing HEC_MTP_ films for 4 days to supraphysiological concentrations of H_2_O_2_ (800µM) that provoked detectable bulk physical changes to films as noted above resulted in a material that was softer (Elastic modulus of 233 kPa), weaker (71% reduction in UTS), and more brittle (13% decrease in strain at failure) (**Figure 4D, S19**). To assess whether physiologically relevant oxidation conditions that do not induce bulk material changes could otherwise promote changes in the surface mechanical properties of fabricated HEC_MTP_ coatings, we performed contact-mode atomic force microscopy (AFM) to determine the micromechanical properties of the material interface after one day of hydration in PBS, then after 7 days of exposure to 80µM H_2_O_2_. This physiologically relevant oxidation of HEC_MTP_ coatings promoted a significant and nearly 3-fold decrease in surface elastic modulus, reducing it from 1718kPa to 674kPa (**Figure 4E**).

**Figure 4:**
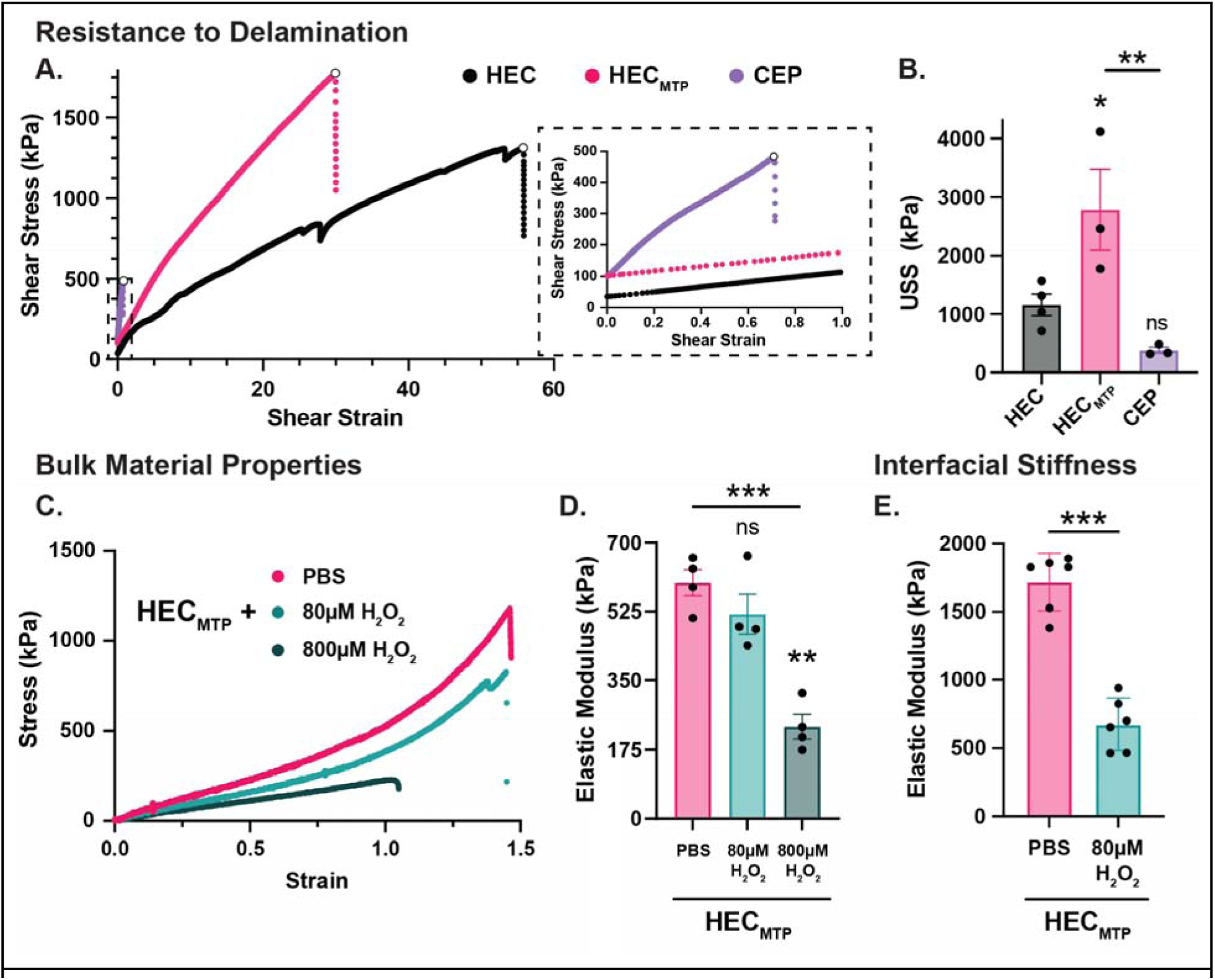
HEC_MTP_ coatings are resistant to delamination and demonstrate softening on oxidation. **(A)** Graph showing representative trace for lap shear test of HEC, HEC_MTP_, and CEP coatings on glass. Inset: Detailed trace from 0.0 to 1.0 shear strain showing the CEP coating failure. **(B)** Bar graph showing the ultimate shear stress (USS) for HEC, HEC_MTP_, and CEP coatings on glass. *P<0.05, **P< 0.009, and not significant (ns), one-way ANOVA with Tukey’s multiple comparison test. Graph shows mean ± s.e.m. with individual data points showing n = 4 for HEC, and n = 3 per group. **(C)** Graph showing representative trace for tensile test of HEC_MTP_ films incubated for 7d in PBS, 7d in PBS + 80µM H_2_O_2_, or 4d in PBS + 800µM H_2_O_2_. **(D)** Bar graph showing the elastic modulus for HEC_MTP_ films, incubated for 7d in PBS, 7d in PBS + 80µM H_2_O_2_, or 4d in PBS + 800µM H_2_O_2_. **P<0.002, ***P< 0.0004, and not significant (ns), one-way ANOVA with Tukey’s multiple comparison test. Graph shows mean ± s.e.m. with individual data points showing n = 4 per group. **(E)** Bar graph showing elastic modulus of HEC_MTP_ coatings incubated in PBS for 1d, then measured by atomic force microscopy (AFM) before and after incubation in PBS + 80µM H_2_O_2_ for 7d. ***P< 0.0001, Student’s t-test (paired). Graph shows mean ± s.e.m. with individual data points showing n = 6.

These data show that HEC_MTP_ forms strongly adhered coatings on glass substrates and that exposure to physiologically relevant oxidizing conditions promotes an effective softening of the surface without imparting bulk changes to HEC_MTP_ coatings and films. Given that surface softening can readily occur under physiologically relevant oxidizing conditions, HEC_MTP_ can be considered a mechanically adaptable material that should soften when implanted *in vivo*.

### 2.5. HEC_MTP_ coatings can be used for optical applications

Optical devices, such as probes, microprisms, optical windows, and GRIN (Gradient-Index) lenses, incorporate transparent glass windows or fibers to image and/or optogenetically control cell functions *in vivo*. However, these optical devices often experience signal instability issues or chronic noise artifacts due to deleterious FBRs stimulated by device implantation.^79^ In anticipation of a potential application for HEC_MTP_ as a surface coating on implanted optical devices to improve *in vivo* performance, we tested the optical properties of HEC_MTP_ coatings to establish their potential compatibility with such devices. Applying HEC_MTP_ coatings to transparent glass caused no observable distortion of underlying features under brightfield microscopy (**Figure 5A**), which we validated using an established, no-reference image quality assessment^80^ to calculate image sharpness (**Figure 5B**). HEC_MTP_ coatings did not attenuate transmitted light within the full spectrum tested from ultraviolet (UV) (300nm) to infrared (IR) (900nm) as assessed by UV-Vis spectroscopy (**Figure 5C**). Brightfield images and UV-Vis spectra measured through glass alone as well as HEC_MTP_ and oxidized HEC_MTP_ coatings demonstrated no detectable change in sharpness (**Figure 5B**) or transmittance (**Figure 5C**), whereas semi-opaque tape and black ink benchmarking controls effectively decreased both transmittance and sharpness in the visible wavelengths (400-700nm) by over 57% (**Figure 5B- C**).

**Figure 5:**
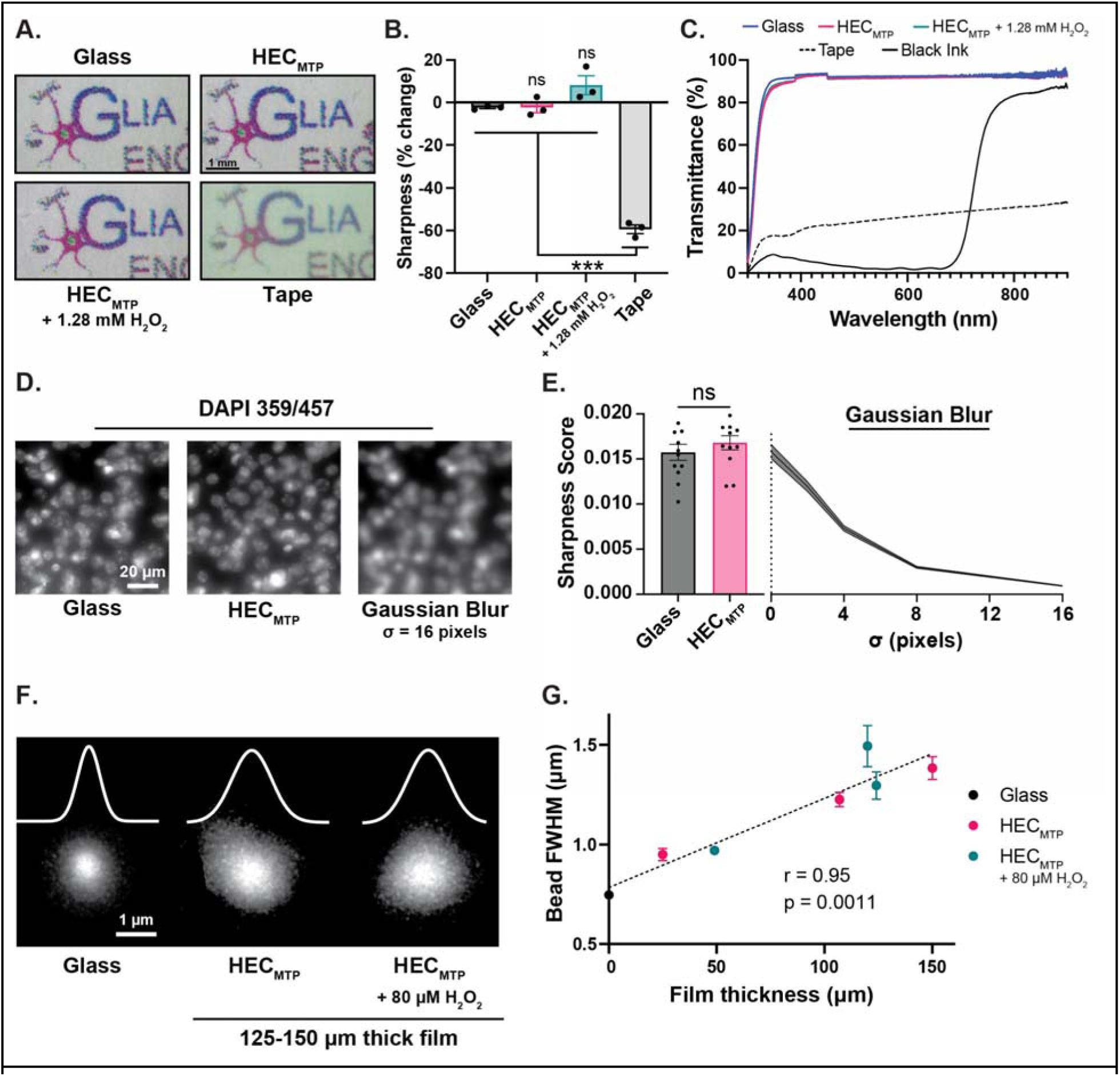
HEC_MTP_ minimally scatters visible and fluorescent light, enabling optical applications. **(A)** Representative brightfield images of printed Glia Engineering logo covered by a glass coverslip with no coating, HEC_MTP_, HEC_MTP_ + 1.28mM H_2_O_2_ for 1d, or opaque tape. **(B)** Graph showing the sharpness %change relative to ground truth (the logo without the coverslip). Not significant (ns) and ***P <0.0001 compared to glass or as designated, one-way ANOVA with Tukey’s multiple comparison test. Graph shows mean ± s.e.m. with individual data points showing n = 3 for all groups. **(C)** Graph showing UV-Vis trace from 300-900nm for glass coverslips that are: uncoated, or have black ink, tape, HEC_MTP_ or HEC_MTP_ + 1.28mM H_2_O_2_ for 1d coatings applied. Each trace shows mean ± s.e.m., with s.e.m. represented as light shaded areas. **(D)** Widefield fluorescence images showing DAPI-stained, coronal, murine brain sections with and without a HEC_MTP_ coating on the glass coverslip, or with a Gaussian Blur filter. **(E)** Bar graph showing the sharpness scores for images of murine brain sections with uncoated or HEC_MTP_ coated glass coverslips. Not significant (ns), Student’s t-test. Graph shows mean ± s.e.m. with individual data points showing n = 12 images per group. Line plot showing the sharpness scores of images of brain sections with uncoated glass coverslips with Gaussian filters of increasing size applied. Trace shows mean ± s.e.m., with s.e.m. represented as light shaded areas, and n = 12 images per group. **(F)** 2-photon fluorescent images of sub-resolution fluorescent beads, covered with a glass coverslip or a glass coverslip with a press fit film of either HEC_MTP_ or HEC_MTP_ + 80µM H_2_O_2_ for 7d. Overlaid traces are one-dimensional Gaussian fits characterizing the bead point spread function (PSF). **(G)** Graph showing the average apparent size of sub-resolution fluorescent beads as a function of film thickness for an uncoated glass coverslip or a glass coverslip with a press fit film of either HEC_MTP_ or HEC_MTP_ + 80µM H_2_O_2_ for 7d. Apparent size was measured by the full width at half maximum (FWHM) of a 1D Gaussian fit. Graph shows mean ± s.e.m. with n = 3 beads per group. Line shows linear regression of all data points, and the correlation is reported as Pearson’s Correlation Coefficient (r).

To assess HEC_MTP_ coating compatibility with fluorescence microscopy, we first stained coronal mouse brain slices (40µm thick) using a DAPI nuclear stain (**Figure 5D**) or primary antibodies for NeuN (neuronal nuclear stain) followed by secondary antibodies conjugated with different common fluorophores that have excitation and emission wavelengths spanning the visible light spectrum (**Figure S20**) and imaged them using a widefield fluorescence microscope. We then calculated the relative change in image sharpness when imaging through HEC_MTP_-coated coverslips (35µm coating thickness) compared to uncoated, glass-only coverslips (**Figure 5D-E, S20**). Qualitative (**Figure 5D, S20**) and quantitative (**Figure 5E**) assessments both demonstrated that the HEC_MTP_ coatings caused no appreciable change in image sharpness relative to the images taken using the uncoated glass. To confirm that this result was not simply caused by limited dynamic range of the scoring algorithm, we also applied standard Gaussian smoothing to the DAPI images, which imparts an incremental and defined blurring of the labeled nuclei (**Figure 5D**). As expected, the application of Gaussian filters of increasing size results in significant reductions in image sharpness, contextualizing the lack of any appreciable shift in image sharpness imparted by HEC_MTP_ coatings (**Figure 5E**).

Next, to further assess any conferred optical effects of our biomaterial in fluorescence microscopy applications, we tested the effect of HEC_MTP_ and oxidized films of different thicknesses on the apparent distortion of sub-diffraction-sized fluorescent beads, when imaged with two-photon microscopy (2PM). Imaging objects such as these beads that are smaller in size than the diffraction-limited resolution of the microscope creates a bead image of larger apparent size, with its fluorescent intensity profile parametrized by a Gaussian distribution in three dimensions. While the ideal apparent bead diameter is equivalent to the diffraction-limited resolution of the imaging system, optical scattering can lead to larger than ideal diameters, ultimately limiting resolution of the image. HEC_MTP_ films were created at thicknesses varying between 25 and 150µm, and then hydrated in PBS for 7 days, while oxidized films at the maximally wettable state were created by incubating dry HEC_MTP_ films in 10mL of 80 µM H_2_O_2_ in PBS for 7 days. After fluorescent beads were dried onto glass slides, the samples were covered with mounting media and then biomaterial films, press-fit onto glass coverslips, were layered over the top. Compared to a coverglass-only control, imaging through HEC_MTP_ and oxidized films visibly altered the lateral point spread function (PSF) of the beads, as shown by one-dimensional Gaussian fits of lateral fluorescent intensity profiles taken through the center of the bead (**Figure 5F**). Changes in apparent resolution were derived from differences in the full width at half maximum (FWHM) for each Gaussian curve, which describes the apparent diameter of the bead. The bead FWHM detected through HEC_MTP_ films showed a small but significant increase compared to the glass only controls, and this deviation was significantly correlated with film thickness (r=0.95) (**Figure 5G**). However, there was no apparent change in conferred resolution between HEC_MTP_ and oxidized films at equivalent film thicknesses (**Figure S20**).

These data show that HEC_MTP_ coatings and films are suitable for diverse optical applications requiring resolution down to the micron scale. While HEC_MTP_ films ultimately impacted fluorescent image resolution on the order of nanometers, the broad array of tested optical applications suggests that sufficiently thin HEC_MTP_ coatings and films are compatible with optical devices, and that this compatibility is not affected by the oxidation-driven, hydrophobic-to-hydrophilic transition.

### 2.6. HEC_MTP_ films promote oxidation-responsive, controlled release of molecules

Poly(propylene sulfide)-based copolymers have been used to formulate oxidation- sensitive drug delivery systems which can selectively release drug cargo via the hydrophobic-to- hydrophilic transition stimulated by sulfoxide formation.^81^ Inspired by these studies, we next tested the capacity for HEC_MTP_ films to act as oxidation-responsive controlled release systems using fluorescently detectable model small molecules and polysaccharide-based macromolecules (**Figure 6A-B**). To characterize small molecule release from HEC_MTP_ films, we used three coumarin derivatives that have different relative hydrophobicities including 4- methylumbelliferone (4MU) (logP = 1.78), 7-Diethylamino- 4-methylcoumarin (C1) (logP=2.90), and coumarin 6 (C6) (logP=4.79). We observed first order release profiles for both 4MU and C1 with rate constants scaling with log P values such that there was complete recovery of loaded 4MU and C1 after 3 and 8 days respectively (**Figure 6A**). Due to its limited aqueous solubility, C6 showed limited release from HEC_MTP_ films under PBS-only incubation over 14 days. The addition of 0.1% triton-X100 surfactant into the incubation media subsequently promoted a sigmoidal pattern of release for C6, resulting in near total recovery over the subsequent 14 days (**Figure 6A**).

**Figure 6:**
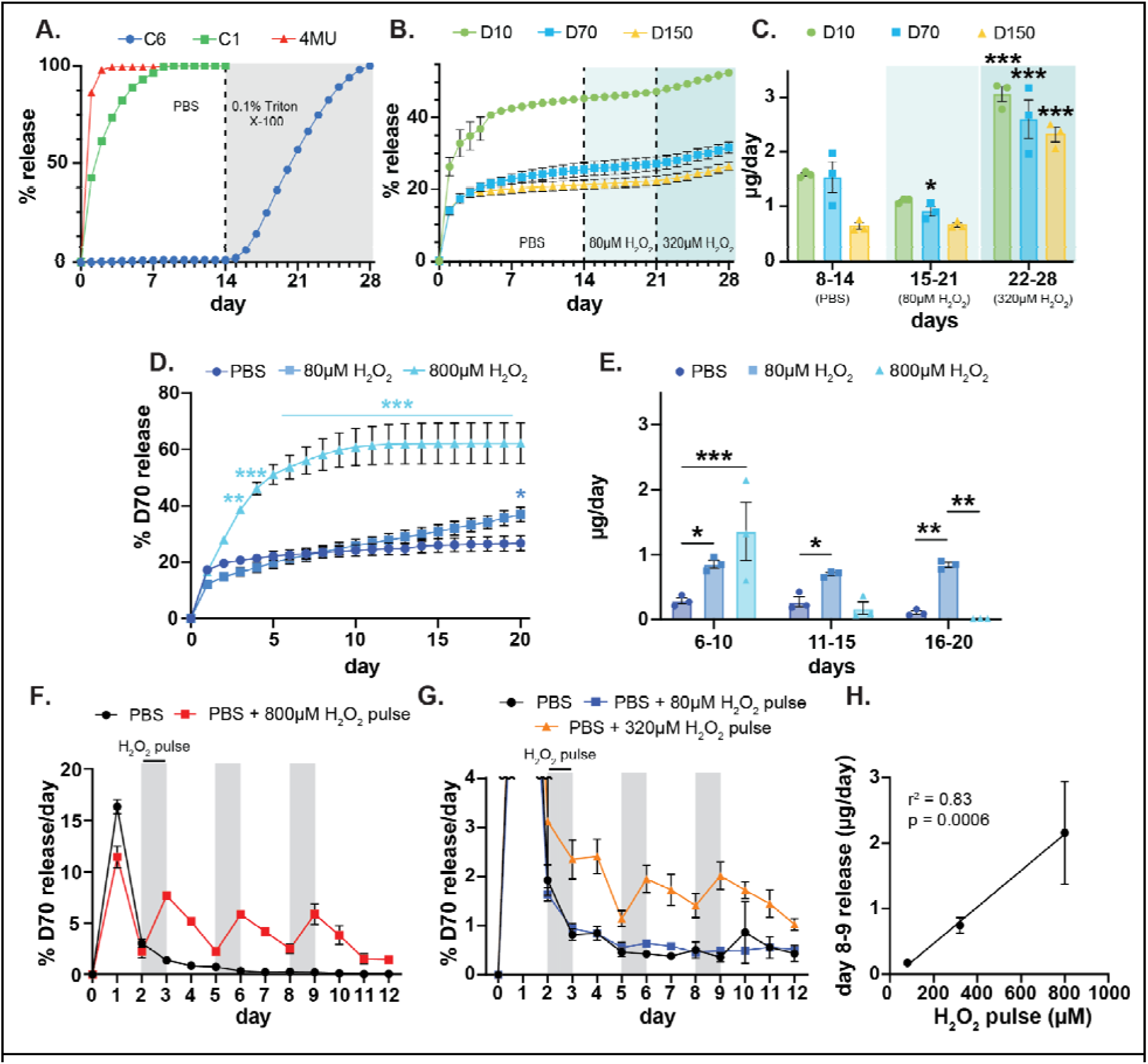
HEC_MTP_ coatings demonstrate controlled drug release. **(A)** Graph showing 28d release profile for Coumarin 6 (C6), 7-Diethylamino-4-methylcoumarin (C1), and 4- methylumbelliferone (4MU). Molecules were released into PBS (d0-d14) or PBS + 0.1% Triton X-100 (d15-d28). **(B)** Graph showing 28d release profile for FITC-dextran polymers (10kDa, 70kDa, and 150kDa). Polymers were released into PBS (d0-d14), PBS + 80µM H_2_O_2_ (d15-d21), or PBS + 320µM H_2_O_2_ (d22-d28). **(C)** Graph showing the average daily dextran for release from d8-d14, d15-d21, and d22-d28. All statistical comparisons are made with the previous duration (ie. D10 d22-d28 is compared with d15-d21) *P<0.05, ***P< 0.0007, and not significant (ns), two-way ANOVA with Tukey’s multiple comparison test. Graph shows mean ± s.e.m. with individual data points showing n = 3 per group. **(D)** Graph showing 20d release profile for 70kDa FITC-Dextran (D70) from HEC_MTP_ films into PBS, PBS + 80µM H_2_O_2_, or PBS + 800µM H_2_O_2_. *P<0.02, **P<0.003, ***P<0.0001 compared to PBS. Two-way ANOVA with Tukey’s multiple comparison test for all 3 groups (800µM H_2_O_2_ comparisons). Two-way ANOVA with Šídák multiple comparison test between 80µM H_2_O_2_ and PBS groups (80µM H_2_O_2_ comparisons). Graph shows mean ± s.e.m. with individual data points showing n = 3 per group. **(E)** Graph showing the average daily D70 release from d6-d10, d11-d15, and d16-d20. *P<0.03, **P<0.008, ***P<0.0002 as designated, two-way ANOVA with Tukey’s multiple comparison test. Graph shows mean ± s.e.m. with individual data points showing n = 3 per group. **(F)** Graph showing 12d daily D70 release profile into PBS with pulses of PBS + 800µM H_2_O_2_ at 2d intervals. **(G)** Graph showing 12d daily D70 release profile into PBS with pulses of PBS + 80µM/320µM H_2_O_2_ at 2d intervals. **(G)** Graph showing relationship between D70 release and H_2_O_2_ pulse concentration. Data was corrected for baseline release using PBS daily release values and line of best fit was fit with a simple linear regression.

To model macromolecule loading and release kinetics, FITC-dextrans with molecular weights of 10, 70, and 150kDa were incorporated into HEC_MTP_ films. These model macromolecules have matched hydrophilicities but hydrodynamic radii that are proportional to molecular weight.^82^ All model dextrans exhibited a modest initial burst release from HEC_MTP_ films when incubated in PBS which was greatest for the 10kDa FITC-dextran at 26%. By 5 days, approximately 41% of total 10kDa dextran cargo was released after which there was only minimal additional release up to 14 days. (**Figure 6B**). Over the same 14-day period, only ∼20% of the total loaded 70 and 150kDa FITC-dextrans were released from HEC_MTP_ films showing minimal but steady release after 7 days. To test for oxidation-stimulated release of the residual dextran cargo in these HEC_MTP_ films, we exposed films to 80µM H_2_O_2_ for 7 days followed by 320µM H_2_O_2_ for an additional 7 days. The 80µM H_2_O_2_ media promoted no significant increase in the rate of dextran release relative to the preceding 7 days in PBS only (**Figure 6C**). However, at 320µM H_2_O_2_ there was a significant near 3-fold increase in the release rates for all dextrans (**Figure 6C**).

To further dissect the oxidation-sensitive release properties of HEC_MTP_ films, we studied the initial release profiles of freshly prepared films loaded with 70kDa FITC-dextran under 80µM and 800µM H_2_O_2_ oxidizing conditions, which we established induces surface and bulk hydrating effects respectively. FITC-dextran released from films exposed to physiologically relevant 80µM H_2_O_2_ showed a near constant, zeroth order release profile following the first day burst which is consistent with molecular release being mediated by material surface hydrating effects (**Figure 6D-E, S21)**. Under these oxidizing conditions, approximately 37% of loaded cargo had been released after 20 days, which was nearly 1.4x the total amount that was released from PBS-only incubated samples during the same timeframe. Incubating films in 800µM H_2_O_2_ resulted in an additional and significant increase in both the total amount and rate of FITC-dextran release, with over 60% of FITC-dextran recovered after just 10 days. Under these supraphysiological oxidizing conditions, the molecular release profile showed first order release kinetics, consistent with bulk-hydration of the material (**Figure 6D-E, S21)**. Next, to test the capacity for HEC_MTP_ films to promote on-demand molecular release upon H_2_O_2_ exposure, we introduced intermittent 24-hour pulses of 800µM H_2_O_2_ (**Figure 6F**), 320µM H_2_O_2_ (**Figure 6G**), or 80µM H_2_O_2_ (**Figure 6G**), to dextran-loaded films after 2, 5, and 8 days of incubation in PBS-only solutions. There was a significant spike in dextran release in response to the three intermittent exposures to H_2_O_2_ and release was attenuated upon removal of the oxidizing stimulus (**Figure 6F-H**). These bursts in dextran release scaled linearly with the concentration of applied H_2_O_2_, further demonstrating that HEC_MTP_ films provide oxidation-responsive release. Since tissue environments at chronically implanted medical devices or wounds can experience local fluctuations in ROS that can be provoked by new injuries or chronic infections, the on-demand release properties of HEC_MTP_ films may be useful for dynamically delivering drugs to regulate such responses.

### 2.7. HEC_MTP_ films are nonfouling substrates that persist *in vivo* with minimal foreign body response for up to 4 weeks

To test the cytocompatibility of HEC_MTP_ coatings, we first cultured quiescent mouse astrocytes on surfaces coated with HEC, HEC_MTP_, or HEC_MTP_ oxidized to its maximal wettability state with 320µM H_2_O_2_ incubation. Astrocytes are a principal cell type involved in the central nervous system foreign body response (FBR) to biomaterials *in vivo,* and therefore represent an appropriate cell type for initial *in vitro* evaluations.^82^ Astrocytes plated on either of the three experimental coatings showed no significant decrease in cell viability relative to tissue culture plastic controls (**Figure 7A**). Next, we characterized astrocyte adhesion to coatings made from HEC_MTP_ and HEC_MTP_ pre-treated with incrementally increasing concentrations of H_2_O_2_ spanning physiological and supraphysiological values. While cultured astrocytes completely adhered and readily spread onto tissue culture plastic, astrocyte adhesion was reduced by more than 57% on HEC_MTP_ coatings, with cells coalescing into discrete multi-cellular aggregates that were approximately 100 μm in diameter (**Figure 7B-C**). Oxidized HEC_MTP_ coatings exposed to at least 20µM H_2_O_2_ showed further attenuated astrocyte adhesion with less than 5% total cell fouling detected across all oxidized HEC_MTP_ conditions. Such minimal cell adhesion was equivalent to that noted on HEC-only coated substrates and is consistent with results observed for other sulfoxide based materials^60^ (**Figure 7B-C**). The attenuated cell fouling results were likely conferred by minimal protein adsorption to HEC_MTP_-based substrates, since HEC_MTP_ and oxidized HEC_MTP_ films showed a nearly 94% reduction in total albumin adsorption compared to a gelatin benchmarking surface by *in vitro* testing (**Figure S22**).

**Figure 7:**
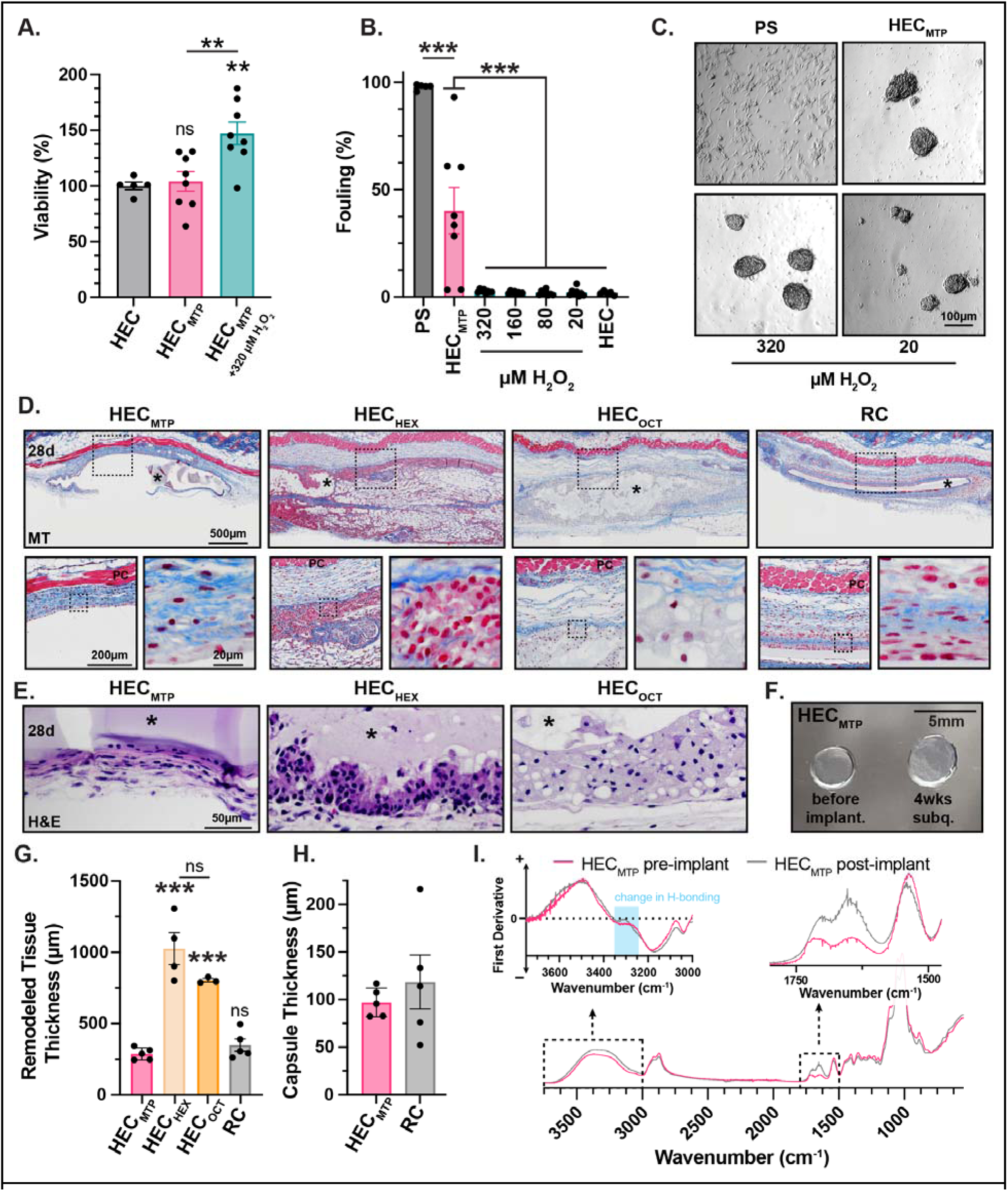
HEC_MTP_ materials are non-fouling *in vitro* and *in vivo*. **(A)** Cytotoxicity assay using Calcein AM demonstrates that coatings of HEC, HEC_MTP_, and HEC_MTP_ treated with 320µM H_2_O_2_ are cytocompatible. Not significant (ns) and **P <0.008 compared to HEC, one-way ANOVA with Tukey’s multiple comparison test. **(B)** Oxidation of HEC_MTP_ with all concentrations of H_2_O_2_ leads to a significant decrease in astrocyte fouling relative to an untreated polystyrene (PS) surface. ***P <0.0001, one-way ANOVA with Tukey’s multiple comparison test. **(C)** Phase contrast images showing morphology of cells seeded on an untreated polystyrene (PS) surface, HEC_MTP_, and HEC_MTP_ treated with either 320 or 20µM H_2_O_2_. **(D)** Survey and detail images of histology sections stained with Masson’s Trichrome (MT) showing implant site and foreign body response (FBR) to subcutaneously implanted HEC_MTP_, HEC_HEX_, HEC_OCT_, and regenerated cellulose (RC) films after 28d. Materials are marked with * and the panniculus carnosus with PC. **(E)** Detail images of histology sections stained with hematoxylin and eosin (H&E) showing FBR to subcutaneously implanted HEC_MTP_, HEC_HEX_, and HEC_OCT_ films after 28d. Materials are marked with *. **(F)** Brightfield images of hydrated HEC_MTP_ films before and after 4wk subcutaneous (subq) implantation. **(G)** Quantification of remodeled tissue thickness for HEC_MTP_, HEC_HEX_, HEC_OCT_, and RC films. Not significant (ns) and ***P ≤0.0003, one-way ANOVA with Tukey’s multiple comparison test. Graph shows mean ± s.e.m. with individual data points showing n = 3 for HEC_OCT_, n = 4 for HEC_HEX_, and n = 5 for HEC_MTP_ and RC. **(H)** Quantification of deep capsule thickness for HEC_MTP_ and RC films. Graph shows mean ± s.e.m. with individual data points showing n = 5 per group. **(I)** FTIR of HEC_MTP_ films pre- and post-implantation. Dashed boxes highlight regions of interest from 1500-1750 cm^-1^ and 3000-3750 cm^-1^. First derivative was smoothed (0th order, 100 neighbors) and inverted.

Having established that HEC_MTP_ coatings are cytocompatible and minimally cell and protein fouling *in vitro*, we next assessed the stability and FBRs of HEC_MTP_ films *in vivo*. Before HEC_MTP_ films could be implanted *in vivo*, we first tested their resilience to standard sterilization methods including autoclaving and soaking in alcohol solutions. We identified minimal changes in HEC_MTP_ dry mass across all conditions (<1%), but soaking HEC_MTP_ coatings in 70% IPA overnight led to a 5-10% decrease in optical transmittance across the 300-900nm spectrum. Based on these findings, autoclaving was deemed to be the most appropriate sterilization method for HEC_MTP_ coatings and films (**Figure S23**). To evaluate the biocompatibility and FBR to the HEC_MTP_ material, we prepared free-standing 3mm diameter HEC_MTP_ films at ∼115µm thickness using an optimized fabrication method (**Figure 2I**). HEC_MTP_ films were sterilized by autoclave and hydrated in sterile PBS for one day before use. Films of HEC_HEX_, HEC_OCT_, and regenerated cellulose (RC) served as benchmarking controls for this *in vivo* FBR analysis. RC films demonstrated limited swelling (144%) after one day and negligible degradation after 7 days, making the physical properties of RC suitable for comparison with HEC_MTP_ (**Figure S24**). Despite the noted swelling and degradation limitations of the HEC_HEX_ and HEC_OCT_ materials (**Figure S24**), we proceeded with evaluating the FBRs of all four materials (HEC_MTP_, HEC_HEX_, HEC_OCT_, and RC) *in vivo* in mice.

For FBR evaluations, we implanted films subcutaneously in pro-fibrotic, C57BL/6 mice by placing hydrated films into pockets made bilaterally in the fascia tissue plane immediately below the panniculus carnosus (PC) muscle layer of the skin in regions on the back just above the hindlimbs. After 4 weeks, we perfused mice and excised skin encompassing the implant site for gross and histological analysis (**Fig 7D-E, S25**). On gross inspection we noted persistence of all implanted HEC_MTP_ films at 4 weeks with minimal change in size, morphology, or transparency. For some, but not all, HEC_MTP_ treated mice, films readily separated from the subcutaneous implant site on dissection due to sparse peri-implant tissue around the film and we were able retrieve these samples as intact films (**Figure 7F, S25**). Neither HEC_HEX_ or HEC_OCT_ films could be identified on gross inspection, however obvious localized regions of differentially-colored scar tissue remained at the implant sites for mice receiving these materials. Histology of explanted films by hematoxylin and eosin (H&E) and Masson’s Trichrome (MT) staining revealed thin and loose peri-implant tissue around HEC_MTP_ films with minimal numbers of persisting immune cells at the film surface and no cell infiltration into the bulk of the material (**Fig 7D-E, S26**). By contrast, both HEC_HEX_ and HEC_OCT_ films showed significant immune cell infiltrates and collagen-rich fibrosis that penetrated throughout pockets of residual polymer material. HEC_HEX_ promoted a more severe immune response compared to all other tested materials with immune cells having larger nuclei as well as cytoplasms that were more basophilic (**Fig 7D-E, S26**). There was a greater extent of immune cell infiltration and fibrosis with organized collagen fibers permeating throughout the implant site in animals receiving HEC_HEX_ compared to all other groups. The immune response to HEC_OCT_ showed notably paler immune cells with more condensed nuclei and large, clear cytoplasmic vacuoles consistent with foamy macrophages, which was unique to this material. Meanwhile, examination of peri-implant tissue at RC films revealed a fibrotic capsule that had a similar abundance of recruited immune cells as HEC_MTP_ both within the fibrotic tissue layers above and below the RC film as well as deposited along the material-tissue interface (**Fig 7D-E, S26, S27A-B**). Quantification of peri-implant fibrotic capsule thickness inclusive of the superficial and deep layers measured from the end of the PC muscle layer verified that HEC_MTP_ films were not significantly different from RC, a widely used, biocompatible biomaterial, and promoted a significantly less severe FBR compared to both HEC_HEX_ and HEC_OCT_ films (**Fig 7G**). Further analysis of the deep fibrotic capsule layer and dense collagen layer around HEC_MTP_ and RC films revealed no significant differences between the two biomaterials (**Fig 7H, S27**). HEC_MTP_ films retrieved from mice after 4 weeks had a small, but detectable, increase in hydrated mass compared to pre-implant films and chemical characterization of retrieved films by FTIR showed changes in the hydroxyl and thiocarbamoyl stretch regions consistent with oxidation-induced sulfoxide conversion of some of the thioether groups along the HEC_MTP_ polymer chain (**Fig 7I**).

These data show that HEC_MTP_ films are cytocompatible and form non-fouling interfaces upon exposure to physiologically relevant oxidizing conditions. HEC_MTP_ is non-resorbable and persists for up to 4 weeks *in vivo* with minimal induced FBR or physical property changes, which was uniquely conferred by its functionalization with MTP groups. Sulfoxide conversion of the pendant thioether groups on the polymer chains in explanted films demonstrated that HEC_MTP_ can undergo the hydrophobic-to-hydrophilic transition *in vivo* under physiological conditions and serves as proof-of-principle evidence that justifies the future exploration of HEC_MTP_ for applications as an oxidation-responsive biomaterial.

## 3. Discussion

Performance longevity of chronically implanted medical devices remains a significant challenge. Hydrophilic polymer coatings or films are a potential way to prevent premature device failure by improving overall biocompatibility. However, issues such as swelling, delamination, and manufacturing scale-up have hampered the effective incorporation of existing coating and film technologies into medical devices.^32–41^ Here, we addressed these issues by functionalizing a readily-sourced and inexpensive cellulose derivative, HEC, with thioether groups via a one-pot, isothiocyanate-based reaction to generate a new, oxidation-responsive biomaterial, HEC_MTP_. HEC_MTP_ can be fabricated into stable, minimally swelling coatings or films that undergo oxidation-driven hydrophobic-to-hydrophilic transitions under physiological conditions to generate non-fouling and mechanically adaptable substrates. Our findings demonstrate proof-of- principle that HEC_MTP_ is a biocompatible and non-resorbable biomaterial *in vivo* and has implications for: 1) extending the functionality of cellulose-based material coatings and films; 2) biomaterial coatings on chronically implanted medical devices; 3) incorporating HEC_MTP_ coatings/films onto optical devices, and 4) using HEC_MTP_ for acute inflammation resolution to improve medical device biocompatibility.

Cellulose is an abundant, inexpensive, and sustainable polymer feedstock that has provided significant societal value over the past century.^42,53^ Cellulose-based materials continue to have broad impact across many industrial applications because of their versatile physical and chemical properties which can be tuned by modifications to polymer structure including the degree and type of chemical substitutions on the repeating anhydroglucose unit.^55^ Here, we add to the growing list of permitted chemical modifications to cellulose materials by developing HEC_MTP_, a new thioether functionalized cellulose. HEC_MTP_ represents an important advance in cellulose-based materials as it can be directed to behave like either a cellulose ester or a hydrophilic cellulose derivative under different conditions, affording unique properties and processing capabilities. Cellulose esters are the predominant class of cellulose materials produced commercially in part because they can be dissolved and easily processed in organic solvents.^53,54^ Here, we show that HEC_MTP_ can be readily dissolved in DMSO and blends of common green solvents enabling a similar level of processability that is critical to fabricating commercially viable coatings and films. However, unlike cellulose esters which are moderately hydrophobic, HEC_MTP_ can be facilely transformed into a hydrophilic polymer by a simple, oxidation-driven conversion of pendant thioether groups to sulfoxides. While coatings made from other hydrophilic cellulose derivatives suffer from swelling and delamination issues,^53^ the controlled hydrophobic-to-hydrophilic transition of HEC_MTP_ that occurs at the surface of the material under physiologically-relevant oxidizing conditions enables the generation of a highly wettable surface without causing bulk material changes that compromise coating integrity or its adhesion to the underlying substrate. Further work will be required to optimize HEC_MTP_ solution parameters, such as polymer concentration and solvent composition, to address the protracted time required for film formation, which is a current limitation of this system, due to the low volatility of the DMSO-only solvent system. In addition to increased manufacturing speed, HEC_MTP_ formulation innovations would allow us to incorporate other coating techniques, such as layer-by-layer assembly or spin coating, to generate thinner HEC_MTP_ coatings with a more uniform surface topography than what was achieved here. Given the modular nature of the thioether functionalization method, important next steps will be to extend our approach to other cellulose derivatives, or to different molecular weight HEC, to tune chemical functionality and/or solution viscosity in order to derive coatings and films with adjustable physical and mechanical properties. Since thioether groups are susceptible to modification by alkylation reactions, the HEC_MTP_ template could also be used to generate other functional materials in future work. For example, reacting HEC_MTP_ substrates with alkyl halides would likely be an effective way to convert thioethers to sulfonium groups and generate potent antimicrobial surfaces.^59^ Also, modification of HEC_MTP_ with haloacetic acids could be used to create new zwitterionic substrates that would likely lead to further improvements in the cell and protein non-fouling properties.^13,59^

Independent of device geometry factors, the main areas where biomaterials engineering can improve long-term performance of chronically implanted medical devices lies in the optimization of surface chemistry, surface topography, and mechanical properties of the device at the device-tissue interface. Biomaterial coatings afford a way to control these surface properties. Ideally, coating technologies must be scalable, readily processable, evoke a minimal foreign body response, be capable of limiting the extent of tissue damage caused by device implantation, exhibit controlled or minimal degradation, and be suitably compatible with primary device functions. Commonly employed biomaterial coatings have also afforded additive functions such as anti-fouling, controlled degradation and drug release, enhanced wear resistance, and electrical conduction or insulation. However, most common synthetic polymers can only permit a few of these functions, and ultimately require concessions to be made to certain other properties. For example, polyethylene glycol (PEG)^13,14^ and zwitterionic polymers (eg. polycarboxybetaine, polysulfobetaine)^13,15,16^ have demonstrated excellent protein and cell anti-fouling, but suffer from swelling, delamination, and fabrication challenges.^32–36^ Materials designed to exhibit controlled degradation, such as poly(lactic acid) (PLA), poly(glycolic acid) (PGA), and poly(caprolactone) (PCL) offer excellent biocompatibility and predictable degradation rates which permit controlled drug release.^83^ However, while these properties can be advantageous in the short term, the exposure of the underlying substrate following degradation of the entire coating may exacerbate the wear, corrosion, or deterioration of device performance in the long term. Wear-resistant coatings such as poly(ethylene)-based coatings (PE, HDPE, UHMWPE)^84^ and Teflon (PTFE)^85^ can provide stiff and low friction surfaces for implanted orthopedic devices and cardiac stents. However, these materials are poor coatings for devices applied to soft tissue, as the mechanical mismatch with host tissue can stimulate persistent inflammation causing chronic tissue damage.^19,20,22^ Some polymers are chosen for certain electrically active medical devices because of their inherent insulative or conductive properties. For example, parylene C and polyimide are materials commonly used as coatings for implanted neural electrodes as they are both excellent insulators and acceptably biocompatible.^86^ Meanwhile, conducting polymers such as polypyrrole and poly(3,4-ethylene doxythiophene) (PEDOT) are also incorporated into neural prosthetics as they maintain conductivity while allowing for drug release and enhanced biointegration.^87^ While these conducting polymers offer promising enhancements to neural implants, they currently suffer from issues with processability including: rigidity/brittleness of coatings, relative insolubility in common solvents, and inability to support post-polymerization modifications.^87^ HEC_MTP_ coatings have the potential to serve critical coating functions while overcoming some of the challenges inherent with other polymer technologies. In particular, we demonstrated here that HEC_MTP_ is anti-fouling, biocompatible, and can provide controlled, responsive drug release, but also that this material exhibits minimal degradation and swelling, effectively resists delamination, and has the capacity to create mechanically adaptable substrates. These unique materials properties distinguish HEC_MTP_ from currently available medical device coating technologies. Future work should focus on extending the properties of HEC_MTP_ coatings to further improve upon its wear resistance, such as by covalently bonding the polymer to the substrate using isocyanate silanation or similar conjugation strategies to chemically couple some of the remaining free hydroxyls on the HEC_MTP_ to the underlying substrate surface. Another opportunity for future improvements of HEC_MTP_ coatings relevant to implanted medical devices could lie in enhancing its conductive properties. While cellulose-based polymers such as HEC_MTP_ are effective insulating materials, by incorporating conductive fillers, such as carbon nanotubes (CNTs) or graphene,^87^ we could enhance the conductivity of HEC_MTP_ coatings and films, allowing the material to be incorporated into the design of new electrode-based devices.

Since HEC_MTP_ coatings minimally alter the optical properties of underlying glass substrates, there are exciting possibilities for their incorporation with chronically implanted optical devices such as transparent glass fibers,^88^ miniature integrated microscopes,^89^ or microprisms.^90^ Such optical devices are becoming increasingly popular tools in preclinical research, and also have emerging clinical therapy applications.^91^ However, without effective coatings to regulate implantation-induced FBRs, these optical devices will continue to be plagued by premature device failure, signal drift over time, and unresolved noise artifacts.^79^ Coatings to prevent leaching of toxic metals from GRIN lenses have been developed^92^, but require a two-step coating process, utilize naturally-sourced biopolymers, and do not account for mechanical compatibility with tissue. Additionally, non-fouling, optically compatible coatings for endoscopes have been investigated for acute clinical purposes^93,94^, but have not been extended to any chronically implanted optical devices. HEC_MTP_ coatings may afford a new way to regulate FBRs at optical devices without compromising optical integrity parameters due to coating swelling or delamination-induced artifacts. However, to realize the potential of HEC_MTP_ for optical devices, more work will be needed to optimize the fabrication of uniform coatings of HEC_MTP_ onto complex device geometries. Furthermore, *in vivo* studies will be required to determine whether *in situ* oxidized HEC_MTP_ surfaces are sufficiently nonfouling to maintain the pristine optical interfaces needed for high imaging fidelity. Incorporation of selective surface oxidation of HEC_MTP_ coatings by either H_2_O_2_ solution incubation or plasma treatment prior to implantation could be integrated into the manufacture of these coating systems if *in situ* oxidation proves to be insufficient.^95^

HEC_MTP_ coatings and films, in addition to modulating surface wettability and cell perceived mechanical properties of implanted medical devices, may also enable new strategies to directly regulate inflammation induced by devices. Neutrophils recruited locally to the surface of implanted medical devices during acute inflammation produce ROS and other mediators as part of antimicrobial defense mechanisms.^96^ Neutrophils are sensitive to the surface properties of medical devices and excessive or prolonged neutrophil accumulation can exacerbate device- adjacent tissue damage, cause biomaterial degradation, or prevent acute inflammation resolution necessary for progression of wound healing processes.^97^ As such, there is growing interest in developing approaches to better control neutrophil functions at implanted medical devices.^98^ The ROS scavenging and ROS-dependent changes in surface properties of HEC_MTP_ coatings and films demonstrated here could make them an effective tool to regulate the oxidative functions of the neutrophil response, mitigate ROS-associated tissue damage, and accelerate acute inflammation resolution.^99^ Furthermore, given the oxidation-responsive, controlled release properties of HEC_MTP_ films also demonstrated here, there is an opportunity to leverage this mechanism for ROS-induced, on-demand delivery of immunoregulatory drugs to better control acute inflammation resolution locally at the sites with the highest neutrophil or ROS burden. To practically realize these strategies, further development efforts will be required to improve upon the ROS-sensitivity of the current system by carefully selecting the cellulose template and degree of thioether functionalization to better tune the physical changes to the polymer induced by the oxidation-driven, hydrophobic-to-hydrophilic transition.

In conclusion, this work describes the derivation and characterization of a new, thioether- functionalized, oxidation-responsive cellulose material. The results from this study lay the foundations for using this material to regulate the FBR to implanted medical devices, including implanted optical systems, and as an oxidation-responsive drug depot.

## 4. Materials and Methods

### Materials

All solvents used in this work are certified ACS grade or higher.

From Sigma Aldrich, we purchased: 2-Hydroxyethyl cellulose (HEC) (purchased Mw: 380kDa) (308633), 3- (methylthio)propyl isothiocyanate (MTPI) (W331201), 4-Methylumbelliferone (M1381-25G), Coumarin 6 (442631-1G), 10kDa Fluorescein isothiocyanate-dextran (FD10S- 100MG), 70kDa Fluorescein isothiocyanate-dextran (FD70S-100MG), 150kDa Fluorescein isothiocyanate-dextran (46946-100MG-F), gelatin from porcine skin (G1890-100G), albumin-fluorescein isothiocyanate conjugate (A9771-250MG).

From Tokyo Chemical Industry Co (TCI), we purchased: 7-Diethylamino-4-methylcoumarin (M0631), N, N-Diisopropylethylamine (DIEA) (D1599).

From ThermoFisher, we purchased: deuterated trifluoroacetic acid (dTFA) (Cat # 325310250), Methyl sulfoxide-d6 (Cat # 364650500), hydrogen peroxide, 30% (Cat # H325-500), B27 (no Vitamin A) (50X) (Cat # 12587010), Octyl isocyanate (Cat # AC41460100), 1-Hexyl isothiocyanate (Cat # AAA1222314).

From Peprotech, we purchased: Epidermal Growth Factor (EGF) (Cat# AF-100-15-100UG), Fibroblast Growth Factor (FGF) (Cat# 100-18B-100UG).

From Gibco, we purchased: Advanced Dulbecco’s Modified Eagle Medium/ Nutrient Mixture F- 12 (DMEM/F12) (Ref# 12634-010), PBS pH 7.4 (1x) (Ref# 10010-023).

From Abcam, we purchased: rabbit anti-NeuN (1:1000, Abcam, ab177487).

From Jackson ImmunoResearch Laboratories, we purchased: AffiniPure™ Donkey Anti-Rabbit IgG secondary antibodies conjugated to Alexa Fluor 488 (AB_2313584), Alexa Fluor 555 (AB_3095471), Alexa Fluor 647 (AB_2492288), or Alexa Fluor 790 (AB_2340628).

From Invitrogen, we purchased: ProLong Gold anti-fade reagent (Cat# P36934). We also purchased: Sylgard 184 (Cat# 24236-10) from Electron Microscopy Sciences, Mettler 40µL Aluminum Crucible (DSC22001) and lid (DSC22002) from DSC Consumables Inc., Calcein AM from Corning (Cat# 354217), 4’,6’-diamidino-2-phenylindole dihydrochloride (DAPI) from Molecular Probe (D1306), SensoLyte® ADHP Hydrogen Peroxide Assay Kit (AS- 71112) from AnaSpec, untreated microscope slides were purchased from Opto-Edu (E35.3501).

### Size-Exclusion Chromatography (SEC)

Prior to use in synthesis, 2-hydroxyethyl cellulose (HEC) was dialyzed, lyophilized and characterized using SEC. Molecular weight and dispersity were determined via SEC using a phosphate buffered saline (pH 7.4) mobile phase on an Agilent 1260 Infinity II GPC/SEC system with a refractive index (RI) detector. Results were standardized using Agilent PEG/PEO standards (PL2080-0201). All calculations were performed using Agilent GPC/SEC software.

### Synthesis of HEC_MTP_, HEC_SO_, HEC_HEX_, and HEC_OCT_

Prior to synthesis, 2-Hydroxyethyl cellulose (HEC) (purchased Mw: 380kDa) was lyophilized, and dimethyl sulfoxide (DMSO), as well as N, n-diisopropylethylamine (DIEA), were dried over 4A molecular sieves for at least 24hrs. HEC was reacted with MTPI to form HEC_MTP_ via a DIEA-catalyzed nucleophilic addition (**Table S1**). Briefly, HEC and MTPI were solubilized in DMSO and purged with nitrogen for 20min before the addition of DIEA. The reaction was heated at 80°C and stirred at 300RPM for 72hrs. The reaction was purified by precipitation in ethyl acetate. The solid was then collected, redissolved in DMSO, and precipitated again in ethyl acetate. The HEC_MTP_ precipitate was then dialyzed in deionized (DI) water for 72hrs with three water changes. HEC_SO_ materials were synthesized by oxidizing HEC_MTP_ in a 1% hydrogen peroxide (H_2_O_2_) solution for 24 hours. Following the reaction, the polymer was dialyzed in DI water and lyophilized as previously described. HEC_HEX_ (**Table S3**) was prepared in an identical manner to HEC_MTP_, with the reactive molecule (MTPI) being substituted for hexyl isothiocyanate. HEC_OCT_ (**Table S4**) was prepared by first solubilizing lyophilized HEC in DMSO at 10mg/mL. Octyl isocyanate was added at the proper stoichiometry, and the reaction was started via the addition of DIEA (**Table S4**). HEC_OCT_ reaction proceeded at 25°C while stirring at 300RPM for 24hrs before being purified as previously described.

### Nuclear Magnetic Resonance (NMR) Spectroscopy

^1^H NMR was performed on an Agilent 500 MHz VNMRS spectrometer at concentrations of ∼10mg/mL and a total volume of 750 µL in DMSO-d6 or dTFA as specified. The spectra were recorded at 117.42kG (1H 500MHz, 13C 125MHz) at ambient temperature. Hydrogen chemical shifts are expressed in parts per million (ppm) relative to the residual proton solvent resonance: DMSO-d6 δ2.50 or δ3.33, or dTFA δ11.50. Following the initial NMR of HEC in dTFA, all subsequent HEC-derivative NMRs were referenced to the HEC-associated peak at δ4.60.

### Fourier Transform Infrared Spectroscopy (FTIR)

FTIR was performed on lyophilized polymer samples using a Nicolet 4700 FTIR-ATR between the wavenumber region 4000-400 cm^-1^ (64 scans, 0.241 cm^-1^ resolution). All FTIR spectra were baseline corrected and normalized by the total area under the curve (AUC) unless otherwise specified.

### Biomaterial Film Fabrication

HEC-derived films were prepared by solubilizing dried material stocks at 3wt% in DMSO and drop-casting HEC_MTP_, HEC_HEX_, or HEC_OCT_ solutions in a silicone mold (150µL of solution in a 10mm diameter, silicone, circular mold). Each layer was dried in a vacuum oven at 35°C and 270 mbar for 24hrs before adding an additional layer. By applying 1 to 3 layers, films could be formed with thickness ranging from 7-70µm.

### X-ray Photoelectron Spectroscopy (XPS)

HEC films were prepared by solubilizing HEC stocks, which had been previously dialyzed and lyophilized, at 6wt%. HEC solution was then pipetted into a silicone mold and dried in a vacuum oven. HEC_MTP_ films were prepared as previously described and treated with PBS or 80µM H_2_O_2_ in PBS for 7d or 800µM H_2_O_2_ in PBS for 1d and then dialyzed against water. Films were then attached to the sample platen and spectra were taken on a PHI Genesis XPS system with an Al K-alpha source after degassing overnight. Binding energy shifts were referenced to the C 1s peak at 284.8 eV.

### Goniometry

Samples were prepared for goniometry by drop-casting a 3wt% polymer solution HEC or HEC_MTP_ in DMSO onto cleaned glass slides. Samples were dried in a vacuum oven for 72hrs at 35°C on house vacuum (∼270 mbar). Samples were then dipped in phosphate buffered saline (PBS) for 20 seconds, blown dry, and tested (**Figure 1H**). For reactive oxygen species (ROS) exposure assay (**Figure 3D**), slides were prepared as previously described. After drying, the coated substrate was exposed to H_2_O_2_ at the specified concentration for 24hrs, at 37°C. Slides were then transferred to a DI water bath to quench the oxidation reaction, blown dry, and tested immediately. Goniometry was performed on a Kruss DSA100 Contact Angle Goniometer by dropping 11µL of DI water onto glass substrates coated with polymer solution. OCA20 Software was used to measure the contact angle relative to the substrate over the course of 2 minutes immediately after dropping. Reported values are the average contact angle over this time.

### Thermal Gravimetric Analysis (TGA)/Differential Scanning Calorimetry (DSC)

TGA/DSC was performed on a Mettler Toledo TGA/DSC. Polymer samples were loaded in a 40µL aluminum crucible, perforated, and tested under nitrogen (50mL/min) over a range of temperature from 30°C to 400°C at a rate of 10°C/min.

### Rheological Testing

Rheological testing was conducted on a TA Instruments DHR-2 Rheometer. ∼200µL of polymer solution was tested during each collection. Viscosity was measured as a function of shear rate on a logarithmic scale with a sampling of five points per decade with shear rates ranging from 0.1- 1000s^-1^. Zero-shear viscosity values are represented as the viscosity values at a shear rate of 0.1s^-1^. Temperature sweeps were performed at a shear rate of 1s^-1^ from 25-80°C. All testing was done using a 20mm 2° cone, except for the 1wt% polymer solution, which was done on a 40mm 1° cone.

### Dipping Mold Fabrication and Dip-Coating Protocol

Polydimethylsiloxane (PDMS) molds were fabricated by mixing a silicone elastomer base and curing agent in a 10:1 ratio in the mold casing. The base was vacuumed for 1hr both before and after adding the curing agent. After vacuuming, a glass insert covered with aluminum foil was placed in the PDMS and placed into an oven at 30°C for 72hrs to cure. The glass insert was removed after drying and the molds were cleaned using DMSO and acetone.

Untreated glass slides were used as a substrate for all coatings. Glass slides were bathed overnight in 1N hydrochloric acid (HCl) to ensure a clean surface. Prior to coating, slides were dipped in HCl, followed by water, then acetone, and blown dry. The pre-made PDMS molds were filled with the appropriate polymer precursor solution. An Instron 5944 Micro-tester tensile testing machine was used to control the speed at which the glass slide was dipped and thereby the shear rate at the interface of the slide. After performing initial rheological testing, it was determined that slides should be dipped at 1 mm/sec (the fastest rate that still approximately maintains zero-shear viscosity or a shear rate of 1 sec^-1^). After dipping, glass slides were dried in a vacuum oven for 72hrs at 35°C on house vacuum (∼270 mbar).

### Coating Thickness and Roughness Measurements

Coating thickness was measured using a Neoteck Digital Thickness Gauge Meter with LCD Display (Range: 0.001-12.7mm/0.00005-0.5”). A Dektak Profilometer was used to measure HEC_MTP_ coating profiles. A tip radius of 5µm with 1mg of applied force was used to scan a 30000µm length over 60 seconds with a resolution of 1.667µm per sampled point and a measurement range of 65.5µm. All roughness data was processed by selecting 1000µm of the HEC_MTP_ coating profile, fitting a linear trendline, and baseline correcting the profile data using this trendline. HEC_MTP_ coating profile data was then centered on 0nm by subtracting the mean of the data from each data point. Average roughness was calculated by taking the average of the absolute value of this processed profile data.

### Scanning Electron Microscopy (SEM)

Samples for SEM were prepared by solubilizing HEC_MTP_ at 1wt% and 3wt% in DMSO. The 1wt% solution was then sterile filtered through a 0.2µm PTFE filter. The 3wt% HEC_MTP_ solution was dropcast onto an aluminum SEM stub (Electron Microscopy Sciences, Cat. # 75225-10) and dried for 3d in a vacuum oven as previously described. The 1wt% HEC_MTP_ solution was dropcast onto an aluminum SEM stub once per day, across 3d, with drying in a vacuum oven overnight. This resulted in a 3-layer film. Both samples were dried on a lyophilizer for 3d before imaging on a Thermo Fisher Phenom ProX SEM, equipped with a 4-segmented backscattered electron (BSE) detector. 4-segmented BSE detector allowed for imaging in three detector modes: Full, Topo A (x-bias), and Topo B (y-bias). Topo A and Topo B imaging modes generate topographic images by subtracting signal detected in the left/right hemisphere (Topo A) or top/bottom hemisphere (Topo B) of the detector.

### H_2_O_2_ Scavenging Experiments

All H_2_O_2_ scavenging experiments were performed by plating 50µL of the HEC material (HEC, HEC_MTP_, HEC_HEX_, HEC_OCT_) solubilized at 1wt% in DMSO in a 96-well plate and drying in a vacuum oven for 7d at 35°C on house vacuum (∼270 mbar). 100µL of H_2_O_2_, freshly diluted from a 30% (9.8M) stock, was plated on top of the HEC_MTP_ coatings, at the appropriate concentration and read after either 24hrs or as noted using the SensoLyte® ADHP Hydrogen Peroxide Assay Kit (AnaSpec, AS-71112).

### Temporal Changes in Contact Angle, Wet Mass, and Dry Mass

Measurements for contact angle, wet mass, and dry mass were performed on HEC_MTP_, drop-cast on glass slides as previously described. The glass slides were pre-weighed to allow for accurate determination of coating mass. After drying, slides were either placed in 25mL of PBS or 80µM H_2_O_2_ in PBS, as designated, and incubated at 37°C. To measure wet mass, slides were removed at 1, 3, and 7d and the surface moisture was removed using compressed air, slides were then weighed on a microbalance. Contact angle was measured as previously described. Dry mass was measured by incubating coated slides for 7d, and then removing the coated slide, drying with compressed air, and lyophilizing the coated slide before weighing on a microbalance.

### Lap Shear Test

Samples were prepared by drop-casting 100 µL of HEC or HEC_MTP_ onto cleaned glass slides and drying in a vacuum oven at 35°C and 270 mbar for 72 hrs. Crosslinked ethoxylated polyol (CEP) controls were prepared by first flashing poly(ethylene glycol) diacrylate (PEGDA 700) and ethoxylated trimethylolpropane tri (3-mercaptopropionate) (ETTMP 1300) over basic alumina. Next, flashed PEGDA 700 and ETTMP 1300 were mixed with 10µL of 2,2-Dimethoxy-2- phenylacetophenone (DMPA) solubilized at 20mg/mL in dimethyl sulfoxide (DMSO) (**Table S5**). This precursor solution was then drop-cast onto the glass substrate and cured under UV ligh (365 nm) for 1 hour. A second glass slide was then superglued (Krazy Glue, All Purpose Super Glue) to the applied sample (**Figure S18**), and the free ends of each slide were coated in epoxy (Gorilla, Clear Epoxy) and allowed to cure for 24 hours. This was employed to prevent slipping at the interface between the glass and the Instron grips, which was initially observed at high applied forces. Samples were positioned vertically with 2kN Instron 2710-114 screw side-action grip attachments. Standard tensile testing was performed at a rate of 5% of slide overlap length per minute until failure. Ultimate shear stress (USS) was defined as the maximum stress before failure.

### Bulk Mechanical Testing

HEC_MTP_ films were fabricated as previously described to create films of 75-150 µm thickness. To constrain mechanical failure to the material midsection, films were then punched into a dogbone shape with gauge length 10mm and gauge width 1.5mm using a Print-A-Punch^TM^ device (Northwest Tissue Mechanics Laboratory, Boise State University, Boise, Idaho).^100^ In addition to dry films, HEC_MTP_ films were then hydrated in PBS for 7 days or incubated in 10 mL of 80 µM H_2_O_2_ in PBS for 7 days. Regenerated cellulose (RC) films were punched from 1kDa RC dialysis bags (Spectrum, P/N: 132104) and hydrated in PBS for 7 days. Using an Instron 5944 Micro-tester tensile testing machine, uniaxial tensile testing was performed on all films at a rate of 1% nominal strain/sec until failure.

### Atomic Force Microscopy (AFM)

Elastic modulus measurements were performed using an Asylum MFP-3D-BIOTM atomic force microscope with silicon nitride tips (Nano World, PNP-TR-Au, Force Constant: 0.32 N/m or 0.08 N/m). HEC_MTP_ was drop-cast on glass slides as previously described, with the addition of a second layer of HEC_MTP_ 24 hours after the first. Measurements were taken first after 24 hours of sample hydration in PBS, then subsequently after 7-day incubation of the same samples in 80µM H_2_O_2_ in PBS at 37°C. Measurements were performed in submerged contact mode, with both the sample and AFM tip completely submerged in PBS. For each sample slide, 3 replicate force-displacement curves were acquired at different locations on the sample surface. Force-displacement curves were fitted with a Hertzian viscoelastic model to determine the elastic modulus, which was reported for each sample as the mean of the replicate measurements.

### UV-Vis Spectroscopy

Glass coverslips (10.5mm x 50mm x 150µm, Chemglass (CLS-1762-1050)) were coated with tape (Scotch, Matte Finish MagicTM Tape), black marker, or HEC_MTP_ solution and dried. Coverslips coated with HEC_MTP_ were additionally treated with 1280µM H_2_O_2_, dipped in 70% isopropyl alcohol (30% water), or autoclaved. Measurements were taken using a Hitachi U-3010 UV–vis Spectrophotometer. Spectra spanned 300–900 nm and were taken at a scan speed of 60 nm min^−1^.

### Brightfield Imaging and Image Sharpness

The ‘Glia Engineering’ logo was printed using a RICOH IM C3500 Color Laser Multifunction Printer onto standard printer paper. Brightfield images were taken using an Axiocam 208 color mounted microscope camera on a Stemi 305 stereo microscope (ZEISS). Image sharpness was calculated using a custom MATLAB script, which first thresholded images in the frequency domain, and then applied an adapted, no-reference scoring algorithm for quantifying image sharpness.^80^

### Widefield Fluorescence Imaging and Image Sharpness

Coronal sections of healthy murine brain were prepared and stained as previously described^101^ either using 4’,6’-diamidino-2-phenylindole dihydrochloride (DAPI; 2 ng mL^−1^; Molecular Probes) to stain cell nuclei or primary antibodies for rabbit anti-NeuN (1:1000). Secondary antibodies were conjugated to: Alexa Fluor 488, Alexa Fluor 555, Alexa Fluor 647, or Alexa Fluor 790. All secondary antibodies were purchased from Jackson ImmunoResearch Laboratories and were affinity-purified whole IgG(H + L) antibodies with donkey host and target species dictated by the specific primary antibody used. Secondary antibodies were diluted 1:500 prior to incubation. HEC_MTP_ films were fabricated as previously described and press fit to coverslips. Coverslips were then placed over coronal sections and imaged on an Olympus IX83 epifluorescent microscope using standardized exposure times. Rectangular regions of interest (ROIs) from fluorescent images of sections with and without a HEC_MTP_ coating were generated and cropped into separate images. Image sharpness was calculated as previously described.^80^ For each fluorescent channel, conferred changes in sharpness scores were reported as the differences between the sharpness scores of a HEC_MTP_-coated ROI and the average sharpness score of all uncoated ROIs. Additionally, changes in sharpness score were calculated for uncoated DAPI ROIs before and after Gaussian blurring filters of increasing size were applied.

### Two-Photon Fluorescence Microscopy

Yellow-green fluorescent beads of 0.2 µm diameter (Polysciences, Cat# 17151) were dried onto glass slides. HEC_MTP_ films were created as previously described, with either 1 or 3 layers applied during fabrication. HEC_MTP_ films were then hydrated in PBS for 7 days or incubated in 10 mL of 80 µM H_2_O_2_ in PBS for 7 days. Control samples were created by covering bead samples with mounting media and a glass coverslip. HEC_MTP_ and HEC_MTP_ + 80 µM H_2_O_2_ samples were created by covering bead samples with a biomaterial film press-fit to a glass coverslip.

Fluorescent bead samples were excited at 950 nm and imaged on a Bruker Ultima Investigator two-photon microscope with a tunable Ti:Sapphire laser (Insight X3, Spectra Physics) and a 16x water-immersion objective (Nikon). Z-stacks were created by raster scanning across a field of view (FOV) of 25 µm x 25 µm, with a pixel size of 24.4 nm. Acquired image stacks were processed using FIJI. From the ∼10 beads visible in each FOV, 3 were selected for analysis. For each bead, the axial centroid was determined from the axial intensity profile, and the corresponding image in the Z-stack was analyzed. A 3.5 µm line was manually drawn through the center of the bead, and the fluorescent intensity profile was fitted with a one-dimensional Gaussian. The full width at half maximum (FWHM) of the Gaussian fit was calculated and reported as the apparent bead diameter.

### Release Assay

To measure release from HEC_MTP_ films and coatings, either small fluorescent molecules (7- Diethylamino-4-methylcoumarin (C1), coumarin 6 (C6), and 4-methylumbelliferone (4MU)) or larger FITC-dextran polymers (10kDa, 70kDa, and 150kDa) were first solubilized in DMSO at 1mg/mL and 10mg/mL, respectively. HEC_MTP_ was solubilized at 3wt% in DMSO and molecule solutions were added to achieve loading ratios of 1µg/mg and 100µg/mg for small molecules and dextrans respectively. Loaded HEC_MTP_ was either drop-cast in a glass scintillation vial to form approximately 4mg HEC_MTP_ coatings, loaded with either 4µg model small molecule or 400µg dextran (**Figure 6A-C**) or drop-cast to make free-standing films (3mm diameter) (**Figure 6D-F**) as previously described. All coatings were incubated in 1mL PBS for the first 14 days, with daily sampling and full buffer replacement. After 14 days, the C6-loaded coatings were switched to a PBS + 0.1% Triton X-100 buffer to enhance solubility, while the dextran-loaded coatings were changed into a solution of PBS + 80µM H_2_O_2_ for 7d and then finally PBS + 320µM H_2_O_2_ for the final 7d (all with daily sampling and full buffer replacement). All films were incubated in 250µL of release buffer (PBS, PBS + 80µM H_2_O_2_, PBS + 800µM H_2_O_2_) and sampled daily with full buffer replacement. For pulsed H_2_O_2_ experiments, HEC_MTP_ films were exposed to 24-hour pulses of 800µM H_2_O_2_, 320µM H_2_O_2_, or 80µM H_2_O_2_ after 2, 5, and 8 days of incubation in PBS-only solutions. All collected samples were read on a Molecular Devices SpectraMax i3X Microplate Detection Platform using an Excitation/Emission appropriate for each molecule (C1 (375/445nm); C6 (443/505nm); 4MU (360/450nm); FITC-dextran polymers (491/516nm))

### Neural Progenitor Cell (NPC) Culture and Differentiation

NPCs were cultured as previously described.^102,103^ Briefly, the cells were grown in neural expansion (NE) media consisting of DMEM/F12 supplemented with B27 and growth factors EGF and FGF (100 ng/mL for each). NPC stocks were maintained so that all cell stocks used in this study were less than P30. For astrocyte differentiation, NE media was removed and replaced with astrocyte differentiation media consisting of DMEM/F12 supplemented with B27 and 1% fetal bovine serum (FBS) for 72hrs.

### Cytotoxicity Assay and Non-fouling Assay

A 96-well plate was coated with 75µL 1wt% HEC, HEC_MTP_, HEC_HEX_ in DMSO for 1 hour at 35°C. The solution was removed, and well plates were dried in a vacuum oven as previously described. After drying, wells were treated with H_2_O_2_ as appropriate, and all wells were washed once with PBS before UV sterilization.

Following differentiation, astrocytes were trypsinized, resuspended in differentiation media, plated onto the well plates at 50k cells/well, and allowed to adhere for 24hrs (original plate). After 24hrs, the supernatant was collected for all wells and added to a separate, tissue culture-treated well plate (transfer plate). The cells were allowed to adhere for 24hrs before testing the viability and fouling. For the dead control, an 80% methanol solution was added to the cells and allowed to incubate for 30min. A fresh stock of Calcein AM solution was prepared at a molarity of 2µM. 100µL of Calcein AM was added to each well and allowed to incubate for 30min. The 96-well plate was read on a Molecular Devices SpectraMax i3X Microplate Detection Platform using an Excitation/Emission of 488/520nm. Viability was determined as the sum of the original plate and the transfer plate, with the results normalized to the live/dead columns where the HEC column represents 100% viability and dead represents 0% viability. Fouling percent was determined as the ratio of Calcein AM signal in the original plate over the sum of the signal from the two plates.

### Protein Fouling Assay

HEC_MTP_ films were created as previously described. HEC_MTP_ films were then punched out using a 1mm diameter biopsy punch and placed into PBS +/- 800µM H_2_O_2_ for 24 hours. HEC_MTP_ films were then transferred into PBS for an additional 24 hours. Gelatin hydrogels were formed by solubilizing gelatin at 50 mg/mL in PBS while heating in a water bath at 85°C. Soluble gelatin was deposited into a silicone mold (3mm diameter) and allowed to cool for 24hrs. Subsequently, gelatin gels were removed from the silicone mold and transferred into PBS for 24 hours. To test protein fouling, hydrated HEC_MTP_ films (+/- 800µM H_2_O_2_) and gelatin were placed in 1mL of 0.5mg/mL albumin-fluorescein isothiocyanate conjugate (FITC-BSA) solubilized in PBS for 1hr. After 1hr, samples were transferred onto a coverslip and imaged with an Olympus IX83 epifluorescent microscope. Images were taken at a standardized exposure time and evaluated using a standard intensity setting. Mean fluorescence intensity was measured within a 500x500µm area at the center of the materials using NIH Image J (1.53) software.

### Biomaterial Implant Formation

To prepare sterile films for implantation, HEC_MTP_, HEC_HEX_, and HEC_OCT_ stocks were first sterile filtered as 1wt% solutions in DMSO and subsequently dialyzed against MilliQ water and lyophilized under minimal particulate conditions. HEC_MTP_, HEC_HEX_, and HEC_OCT_ films were then fabricated as previously described. Layering and drying steps were repeated to form films of thickness 100-200µm. Regenerated cellulose (RC) implants were formed from 1kDa RC dialysis bags (Spectrum, P/N: 132104). These materials were dialyzed and lyophilized under the same conditions to form a dry film. Individual subcutaneous implants were then punched from these films using a 3mm biopsy punch. Subcutaneous implants were autoclaved and stored sterile until implantation. Prior to implantation, subcutaneous implants were hydrated in sterile PBS for 24hrs (HEC_MTP_, RC) or at the time of surgery (HEC_HEX_, HEC_OCT_).

### Subcutaneous Implant Surgery

All surgical procedures were approved by the BU IACUC (Protocol number: PROTO202100013) and conducted within a designated surgical facility. All procedures were performed on mice C57/BL6 female mice (Cat#000664, JAX) that were aged 8–12 weeks at the time of surgery under general anesthesia achieved through inhalation of isoflurane in oxygen-enriched air. The dorsal regions of the mice were shaved, stabilized, and horizontally leveled using a stereotaxic apparatus with ear bars (David Kopf, Tujunga, CA, USA) and a custom device to hold the mouse tail taut. A small incision was made on the dorsum and the subcutaneous tissue was separated from muscle via blunt dissection. Placement of hydrated subcutaneous implants (HEC_MTP_, HEC_HEX_, HEC_OCT_, and RC) was visually aided by an operating microscope, with each mouse receiving 4 implants (2 per side). After placement of subcutaneous implants, the surgical incision site was sutured closed, and the animals were allowed to recover.

### Transcardial Perfusions

After terminal anesthesia by an overdose of isoflurane, mice were perfused transcardially with heparinized saline (10 units mL^−1^ of heparin) and 4% paraformaldehyde (PFA) that was freshly prepared from 32% PFA aqueous solution (Cat# 15 714, EMS), using a peristaltic pump at a rate of 7 mL min^−1^. Approximately 10 mL of heparinized saline and 50 mL of 4% PFA were used per animal. Following perfusion, the skin (epidermis, dermis, subcutaneous) was resected, and the 4 subcutaneous implants were located. This tissue was placed in 4% PFA for 48 hours, then transferred into 0.01% sodium azide in Tris-buffered saline (TBS) for at least 3d with the replacement after 2d and stored at 4 °C until further use.

### Histological Staining, Brightfield Imaging, and Image Analysis

Tissue samples were embedded, cut, and stained with hematoxylin and eosin (H&E), as well as Masson’s Trichrome (MT) by the Tufts Animal Histology Core. All histological images were taken using an Olympus Virtual Slide Microscope (VS120). Quantification of peri-implant, fibrotic capsule thickness (defined as all collagenous tissue below the PC muscle layer) and dense collagen capsule thickness (defined by intensity of MT stain and fiber morphology at implant surface) was performed using NIH Image J (1.53) software. Dense collagen capsule thickness was calculated by dividing the measured dense collagen area by the average of the outer and inner perimeters. Missing or defective portions of the capsule were excluded. Morphology-based classification of cells immediately adjacent to biomaterial implant was performed using the QuPath software.^104^ Fibroblasts were defined as cells with elongated nuclei whereas round or irregular cells were classified as macrophages. Large, multinucleated cells were classified as foreign body giant cells (FBGCs) and cells with lobed nuclei were defined as neutrophils. Due to relative sparsity, FBGCs and neutrophils were excluded from further analysis. Cellular densities were obtained by counting the number of cells within 20 µm of each implant (tissue sections were 5µm thick).

### Statistical Analysis

Graph generation and statistical evaluations of repeated measures were conducted by one-way or two-way ANOVA with post hoc independent pairwise analysis via Tukey’s multiple comparison test or by Student’s t-tests where appropriate using Prism 10 (GraphPad Software Inc, San Diego, CA). Statistical details of experiments can be found in the figure legends, including the statistical tests used and the number of replicate samples. Across all statistical tests, significance was defined as p-value <0.05. Unless otherwise specified, all graphs show mean values plus or minus standard error of the means (s.e.m.) as well as individual values overlaid as dot plots.

## Supporting Information

Supporting Information is available from the Wiley Online Library or from the author.

## Acknowledgments

This research was supported by the Biomedical Engineering Core Facilities at Boston University (BU). A special thanks to Xin Brown and the Biointerface Technologies (BIT) core, as well as the Micro and Nano Imaging (MNI) core at BU. This research was also supported by the BU Chemistry Department Chemical Instrumentation Center and staff. We thank the Boston University Neurophotonics center for providing the Bruker Ultima Investigator two-photon microscope used in this work. We are grateful to the National Science Foundation for the purchase of the NMR’s (CHE0619339) and PHI Genesis XPS system (CHE-2216008) used in this work. We also thank the Boston University Micro and Nano Imaging Facility and the Office of the Director, National Institutes of Health for their support under Award Number S10OD024993 (this content is solely the responsibility of the authors and does not reflect the official views of the NIH). We thank Lu Ping for help with XPS, Cara Ravasio for many helpful discussions, and both Patrick Kulaga and Jacob Labovitz for time spent on histological analysis. We also thank the Tufts Animal Histology Core for their processing and staining of the tissue.

## Competing interests

Authors declare that they have no competing interests. A patent application describing the polymer technology has been submitted.

Data and materials availability

All data needed to evaluate the conclusions in the paper are present in the paper and/or the Supplementary Materials. Materials can be made available upon request.

