## Supplementary Information for "Thioether-functionalized cellulose for the fabrication of oxidation-responsive biomaterial coatings and films"

**Table S1: Example reaction stoichiometry for $HEC_{MTP}$ synthesis.**

| Reagent | Molecular Weight | Equivalence (Eq) | mols (mmols) | mass (mg) | density (g/mL) | volume (mL) | Eq/free hydroxyl |
|---|---|---|---|---|---|---|---|
| **Hydroxyethyl cellulose (HEC)** | 380000 | 1 | 0.00263 | 1000 | – | 0 | – |
| **3-(methylthio)-propyl isothiocyanate (MTP-ITC)** | 147.26 | 59221 | 155.844 | 22949.61 | 1.1 | 20.86 | 30 |
| **Dimethylsulfoxide (DMSO)** | – | – | – | – | – | 100 | – |
| **n,n-diisopropylethylamine (DIEA)** | 129.25 | – | – | – | – | 0.5 | – |

**Table S2: $HEC_{MTP}$ synthesis reaction blends.**

| Reaction # | mol% Functionality |
|---|---|
| **1** | 18 |
| **2** | 16 |
| **3** | 19 |
| **4** | 23 |

**Table S3: Example reaction stoichiometry for $HEC_{HEX}$ synthesis.**

| Reagent | Molecular Weight | Equivalence (Eq) | mols (mmols) | mass (mg) | density (g/mL) | volume (mL) | Eq/free hydroxyl |
|---|---|---|---|---|---|---|---|
| Hydroxyethyl cellulose (HEC) | 380000 | 1 | 0.00263 | 1000 | – | 0 | – |
| Hexyl isothiocyanate (HEX-ITC) | 143.25 | 66130 | 174.026 | 24929.27 | 0.93 | 26.835 | 33.5 |
| Dimethylsulfoxide (DMSO) | – | – | – | – | – | 100 | – |
| n,n-diisopropylethylamine (DIEA) | 129.25 | – | – | – | – | 0.5 | – |

**Table S4: Example reaction stoichiometry for $HEC_{OCT}$ synthesis.**

| Reagent | Molecular Weight | Equivalence (Eq) | mols (mmols) | mass (mg) | density (g/mL) | volume (mL) | Eq/free hydroxyl |
|---|---|---|---|---|---|---|---|
| Hydroxyethyl cellulose (HEC) | 380000 | 1 | 0.00105 | 400 | – | 0 | – |
| Octyl isocyanate (OCT-IC) | 155.24 | 3158 | 3.325 | 516.12 | 0.88 | 586.5 | 1.6 |
| Dimethylsulfoxide (DMSO) | – | – | – | – | – | 100 | – |
| n,n-diisopropylethylamine (DIEA) | 129.25 | – | – | – | – | 0.1 | – |

**Table S5: Example stoichiometry for crosslinked ethoxylated polyol (CEP) formulation.**

| Reagent | Molecular Weight | Equivalence (Eq) | mols (mmols) | mass (mg) | density (g/mL) | volume (mL) |
|---|---|---|---|---|---|---|
| **poly(ethylene glycol) diacrylate (PEGDA 700)** | 700 | 3 | 1.286 | 900 | 1.12 | 0.804 |
| **ethoxylated trimethylolpropane tri (3-mercaptopropionate) (ETTMP 1300)** | 1300 | 2 | 0.857 | 1114.3 | 1.144 | 0.974 |
| **2,2-Dimethoxy-2-phenylacetophenone (DMPA)** | 256.3 | 0.01 wt% | 0.00079 | 0.201 | – | – |

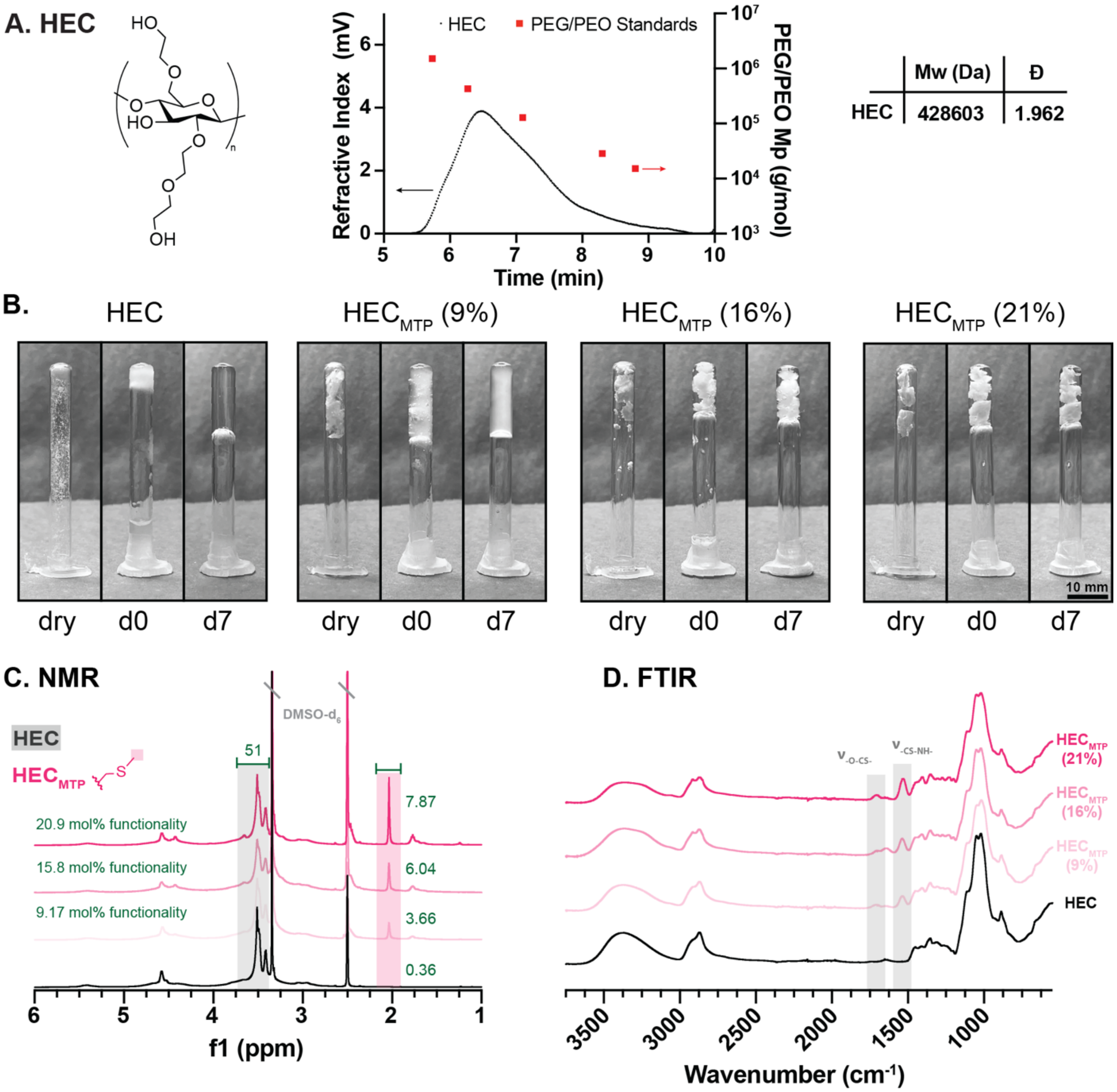


**Figure S1: Differences in HEC$_{MTP}$ functionality yields either a gel-like or an insoluble material in PBS. (A)** HEC structure and size exclusion chromatography (SEC) trace with resulting molecular weight (Mw) and dispersity (Đ). **(B)** HEC and HEC$_{MTP}$ at 9, 16, and 21mol% functionality were solubilized at 5wt% in PBS and allowed to equilibrate for a week. Images are taken of the dry polymer (dry), after the addition of PBS (d0), and after 1 week of equilibration at room temperature (d7). **(C)** $^{1}$H NMR of HEC and HEC$_{MTP}$ at 9, 16, and 21mol% functionality in dDMSO with integration of MTP-associated peaks. **(D)** FTIR of HEC and HEC$_{MTP}$ at 9, 16, and 21mol% functionality highlighting MTP-associated peaks.

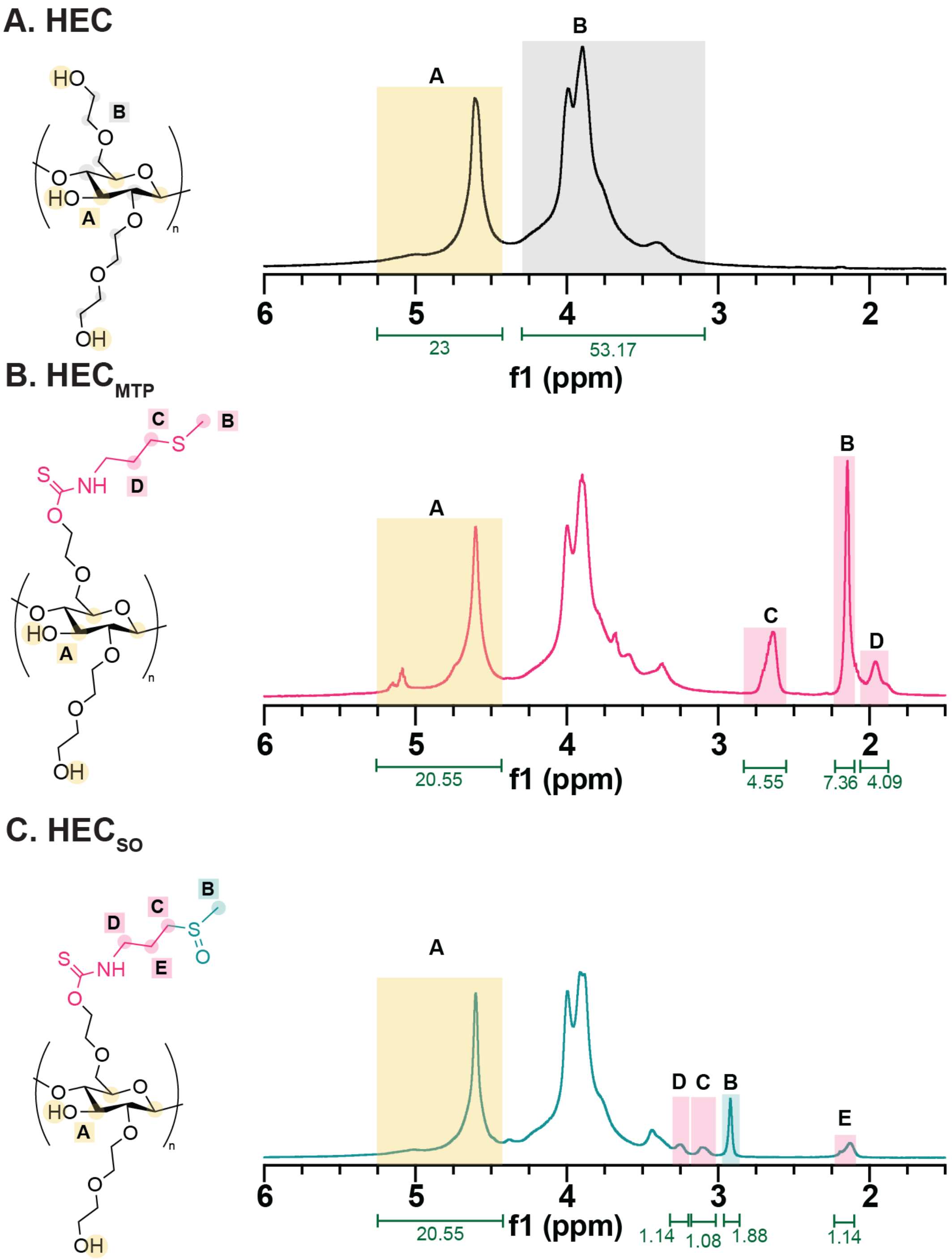


**Figure S2: NMR characterization of HEC, HEC$_{MTP}$, and HEC$_{SO}$. (A)** $^1$H NMR of HEC in dTFA with integration. **(B)** $^1$H NMR of HEC$_{MTP}$ in dTFA with integration normalized to conserved HEC peaks and highlighting MTP-associated peaks. **(C)** $^1$H NMR of HEC$_{SO}$ in dTFA with integration normalized to conserved HEC peaks and highlighting SO-associated peaks.

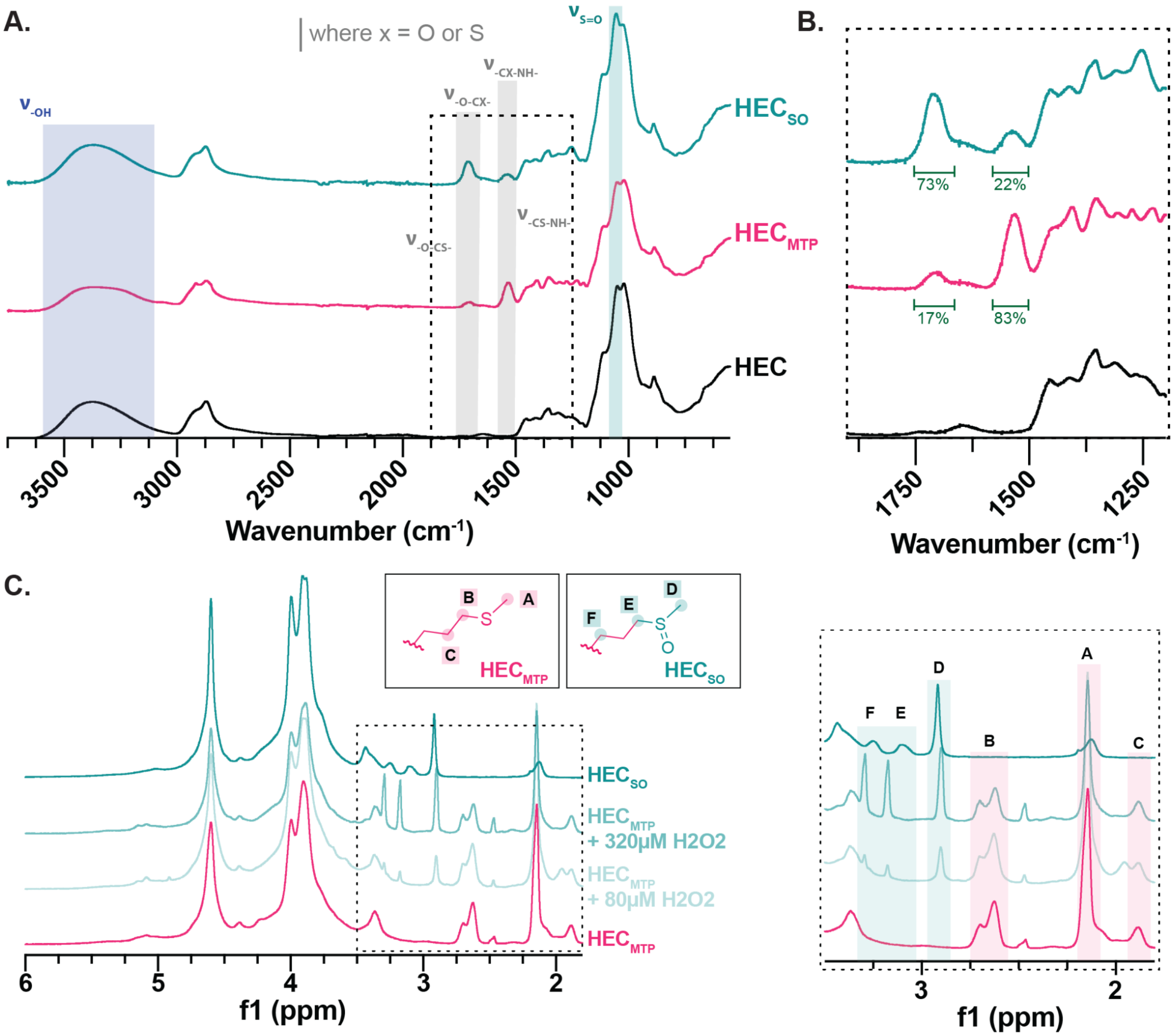


**Figure S3: FTIR and NMR characterization of HEC, $HEC_{MTP}$, and $HEC_{SO}$ reveals maintenance of MTP functionality after oxidation. (A)** FTIR with characteristic absorption bands of chemical bonds in HEC, $HEC_{MTP}$ and $HEC_{SO}$. Spectra is normalized so that the total area under the curve (AUC) for each condition is 1000. Highlights demonstrate MTP and SO -associated peaks. **(B)** Detailed image showing FTIR spectra region from 1200-1900$cm^{-1}$ with green integration showing the AUC for the thiocarbamoyl stretches, normalized such that the total AUC for both thiocarbamoyl stretches for $HEC_{MTP}$ add to 100%. **(C)** $^1$H NMR of $HEC_{MTP}$, $HEC_{SO}$, and $HEC_{MTP}$ treated with either 80µM or 320µM $H_2O_2$ in dTFA highlighting MTP-associated and SO-associated peaks.

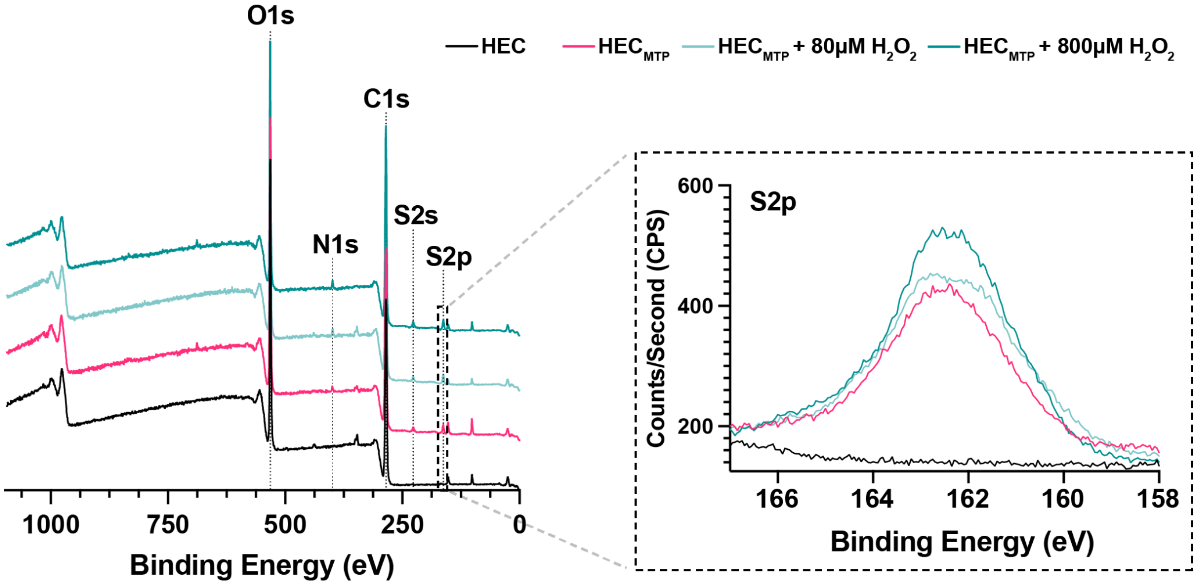


**Figure S4: XPS characterization of HEC, HEC$_{MTP}$ and oxidized HEC$_{MTP}$.** Survey scan of HEC, HEC$_{MTP}$ and oxidized HEC$_{MTP}$ (80µM $H_2O_2$ for 7d and 800µM $H_2O_2$ for 1d) shows the presence of MTP-related peaks (N1s, S2s, S2p) in HEC$_{MTP}$ and oxidized HEC$_{MTP}$ materials and absence of these peaks in HEC. Inset plot shows a high-resolution scan of the characteristic sulfur 2p (S2p) peak.

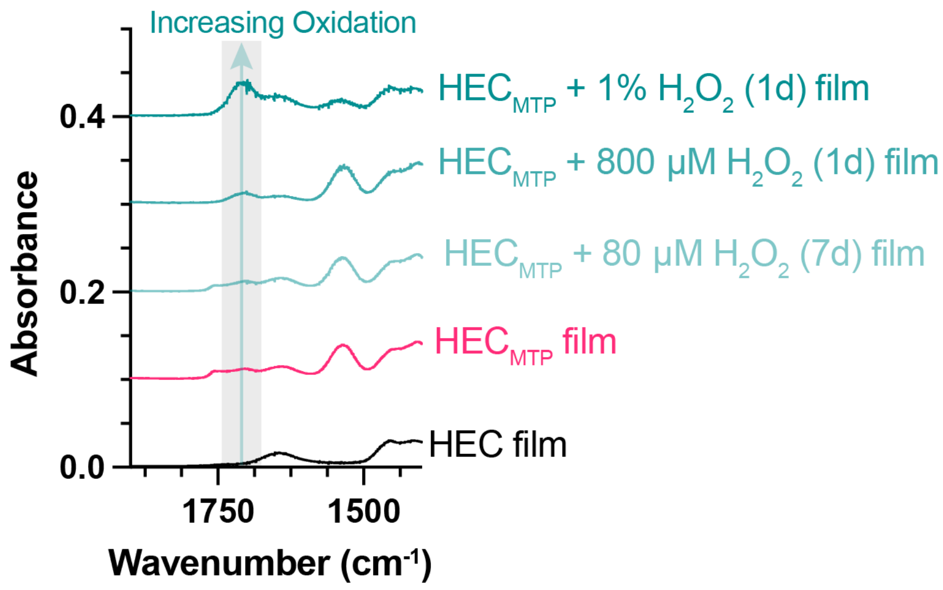


**Figure S5: FTIR of HEC, $HEC_{MTP}$ and oxidized $HEC_{MTP}$ films shows minimal changes to thiocarbamoyl stretches.** FTIR of HEC, $HEC_{MTP}$ and oxidized $HEC_{MTP}$ films (80µM $H_2O_2$ for 7d, 1% $H_2O_2$ and 800µM $H_2O_2$ for 1d). Gray highlight shows -O-CO- stretch, associated with increased oxidation, which only begins to emerge at supraphysiological $H_2O_2$ concentrations.

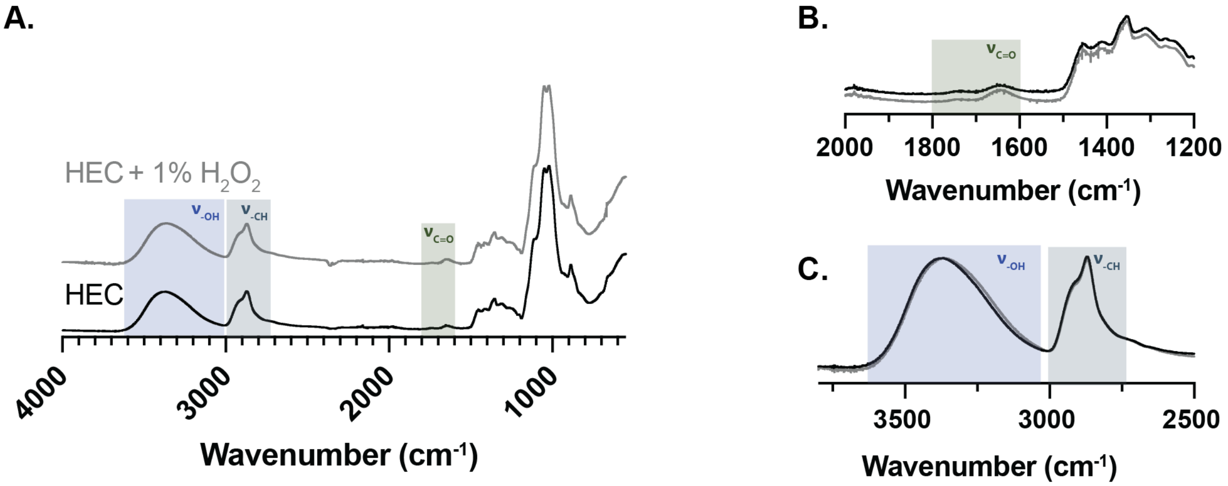


**Figure S6: FTIR characterization of HEC and HEC treated with 1% $H_2O_2$ for 24hrs. (A)** Full FTIR with highlights show key HEC-related peaks. **(B)** Magnified plot from 1200-2000 $cm^{-1}$ demonstrating no detectable emergence of carbonyl stretch (aldehyde) that would suggest oxidation of the cellulose backbone or ethylene oxide side chains. **(C)** Magnified plot from 2500-3800 $cm^{-1}$ demonstrating no detectable change in hydroxyl stretch.

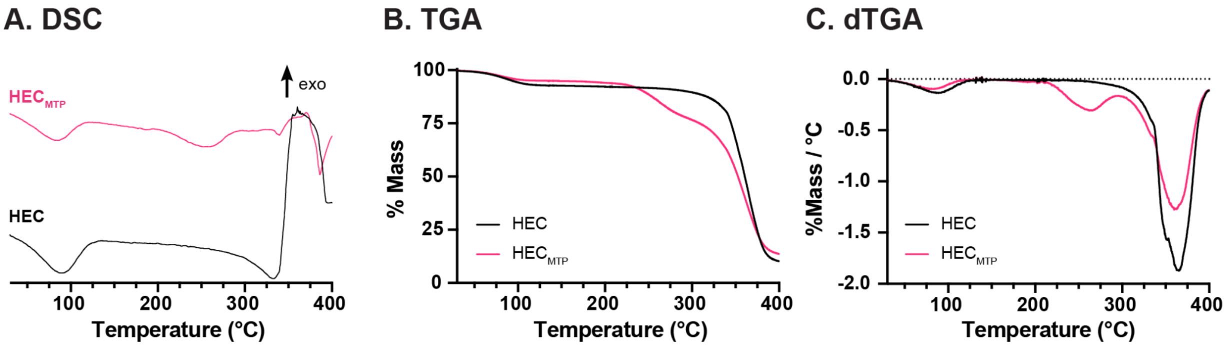


**Figure S7: DSC/TGA characterization of HEC and $HEC_{MTP}$. (A)** Differential scanning calorimetry (DSC) plots for HEC and $HEC_{MTP}$ taken from 30°C to 400°C. **(B)** Thermogravimetric analysis (TGA) of HEC and $HEC_{MTP}$ between 30°C and 400°C demonstrating that HEC and $HEC_{MTP}$ are thermally stable up to 215°C after which $HEC_{MTP}$ loses approximately 20% of its total mass. **(C)** dTGA plot derived from TGA for HEC and $HEC_{MTP}$.

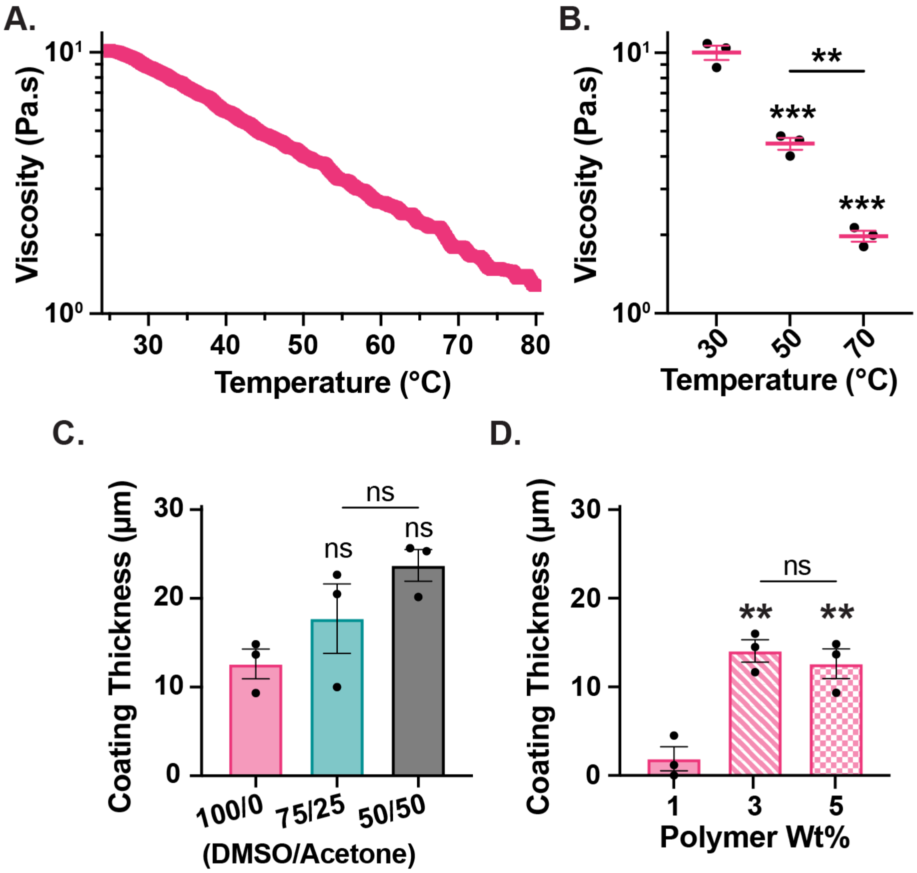


**Figure S8: $HEC_{MTP}$ solution viscosity is temperature dependent and solution viscosity dictates coating thickness. (A)** Dynamic rheological shear rate sweep measurements for $HEC_{MTP}$ solubilized at 5wt% in a 100/0 DMSO solvent system showing viscosity (Pa.s) against increasing temperature (°C). **(B)** Box-and-whisker plot showing the decrease in measured viscosity (Pa.s) at fixed temperature values. $^{**}P < 0.01$ and $^{***}P \leq 0.0001$ compared to 30°C. **(C)** Bar graph showing effect of solvent system on coating thickness at a fixed concentration of 5wt%. Not significant (ns) compared to 100/0 DMSO/Acetone or as designated. **(D)** Bar graph showing effect of concentration on coating thickness in a fixed solvent system (100/0 DMSO/Acetone). $^{**}P < 0.005$, not significant (ns) compared to 1wt% or as designated. All statistical tests were one-way ANOVAs with Tukey's multiple comparison tests. Graphs show mean ± s.e.m. with individual data points showing n = 3 per group.

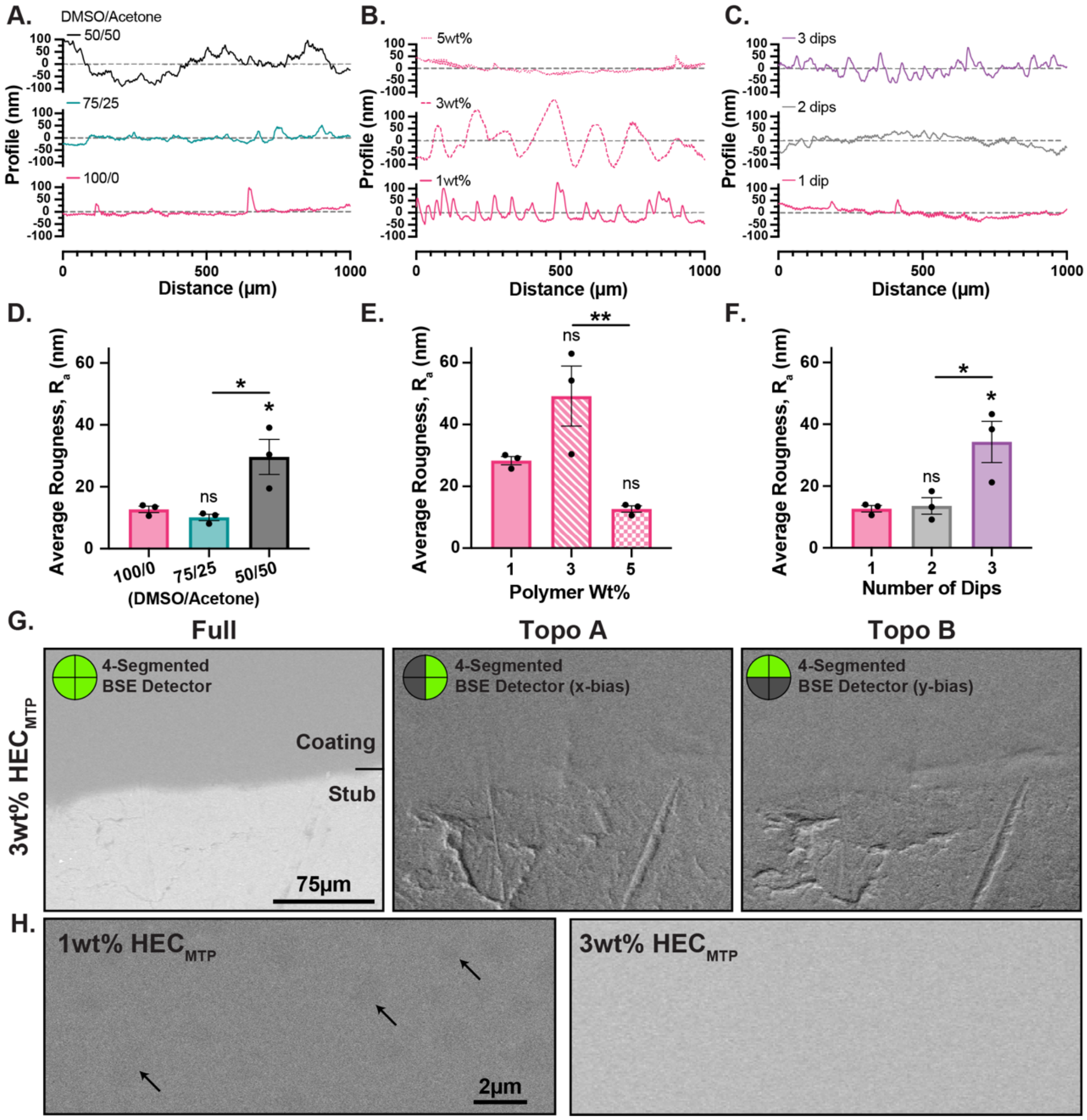


**Figure S9: Solvent system, concentration, and number of dips can affect coating surface roughness.** Profilometry data showing the measured surface roughness across 1 mm of coating for different **(A)** solvent systems (DMSO/Acetone) at 5wt% **(B)** concentrations in 100/0 DMSO/Acetone and **(C)** number of dips in 100/0 DMSO/Acetone at 5wt%. Roughness trace has been baseline corrected and the average displacement set as 0 nm. Gray dashed line shows the roughness of the glass without a coating. **(D)** Bar graph showing the effect of solvent system on the average roughness of the coating. $^*P < 0.03$ compared to 100/0 DMSO/Acetone or as designated. **(E)** Bar graph showing the effect of concentration on the average roughness of the coating. $^{**}P < 0.01$ compared to 1wt% or as designated. **(F)** Bar graph showing the effect of multiple dips on the average roughness of the coating. $^*P < 0.04$ compared to 1 dip or as designated. All statistical tests were one-way ANOVAs with Tukey's multiple comparison tests. Graphs show mean ± s.e.m. with individual data points showing n = 3 per group. **(G)** Scanning electron microscopy (SEM) micrograph of $HEC_{MTP}$ coated on an aluminum SEM stub at 3wt% imaged with 4-segmented backscattered electron (BSE) detector. Micrographs show region of interest (ROI) imaged with three detector modes: Full, Topo A (x-bias), and Topo B (y-bias). **(H)** SEM of $HEC_{MTP}$ coated on an aluminum SEM stub at 1wt% (3 layers) and 3wt% (1 layer). Micrographs were imaged on Full detector mode and arrows highlight small surface features on 1wt% $HEC_{MTP}$ coating.

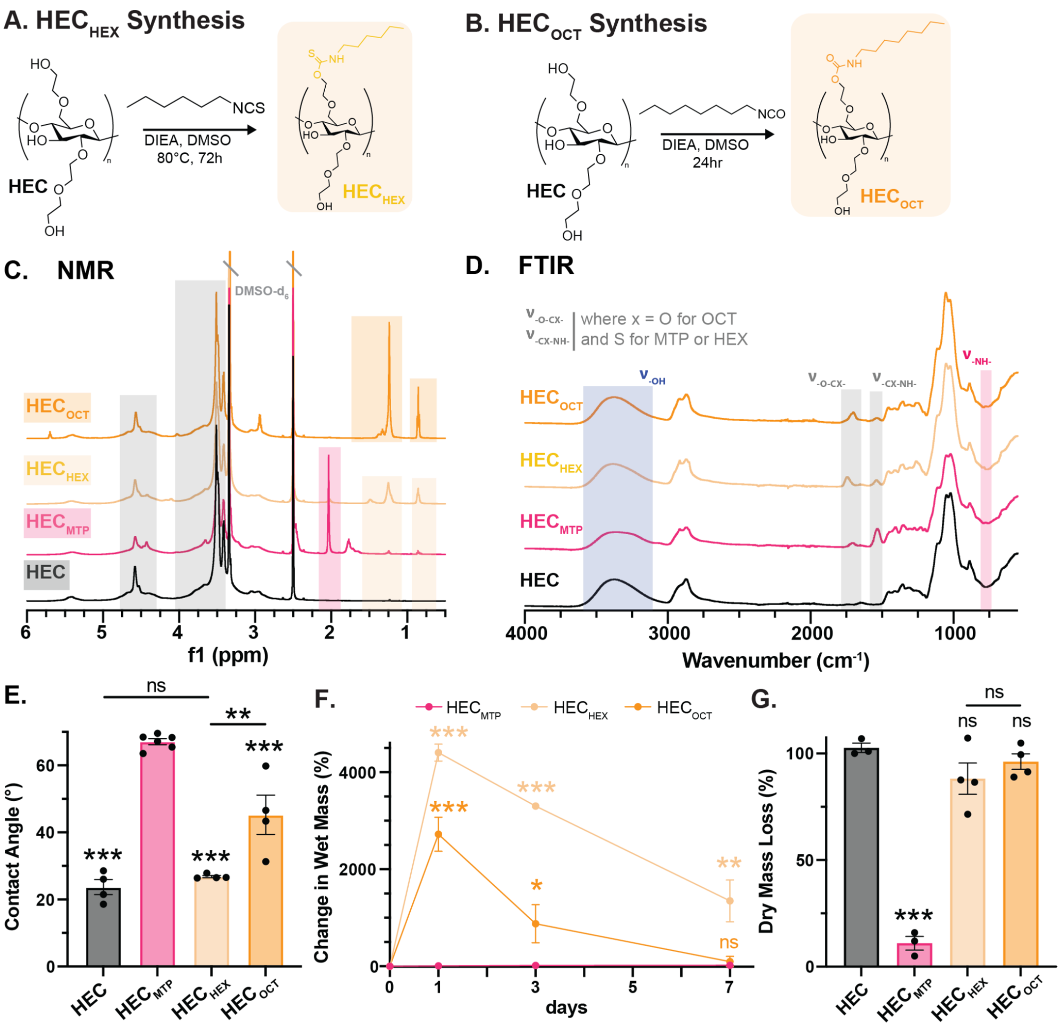


**Figure S10: HEC$_{HEX}$ and HEC$_{OCT}$ characterization. (A)** Synthesis scheme for base-catalyzed, nucleophilic addition of HEX-ITC to HEC to prepare HEC$_{HEX}$. **(B)** Synthesis scheme for base-catalyzed, nucleophilic addition of OCT-IC to HEC to prepare HEC$_{OCT}$. **(C)** $^1$H NMR of HEC, HEC$_{MTP}$, HEC$_{HEX}$, and HEC$_{OCT}$ in dDMSO. **(D)** FTIR characterization HEC, HEC$_{MTP}$, HEC$_{HEX}$, and HEC$_{OCT}$, highlights demonstrate MTP, HEX, and OCT -associated peaks. **(E)** Contact angle data for HEC, HEC$_{MTP}$, HEC$_{HEX}$, and HEC$_{OCT}$. Not significant (ns), **$P < 0.004$, and ***$P \leq 0.0008$ compared to HEC$_{MTP}$ or as designated, one-way ANOVA with Tukey's multiple comparison test. Graph shows mean ± s.e.m. with individual data points showing n = 6 for HEC$_{MTP}$ and n = 4 for all other groups. **(F)** Change in wet mass of HEC$_{MTP}$, HEC$_{HEX}$, and HEC$_{OCT}$ coatings over 7d. Not significant (ns), *$P < 0.04$, **$P < 0.0015$, and ***$P < 0.0001$ compared to HEC$_{MTP}$, two-way ANOVA with Tukey's multiple comparison test. Graph shows mean ± s.e.m. with individual data points showing n = 3 for HEC$_{MTP}$ and n = 4 for all other groups. **(G)** %Change in dry mass of HEC, HEC$_{MTP}$, HEC$_{HEX}$, and HEC$_{OCT}$ coatings after 7d. Not significant (ns), and ***$P < 0.0001$ compared to HEC or as designated, one-way ANOVA with Tukey's multiple comparison test. Graph shows mean ± s.e.m. with individual data points showing n = 3 for HEC and HEC$_{MTP}$ and n = 4 for all other groups.

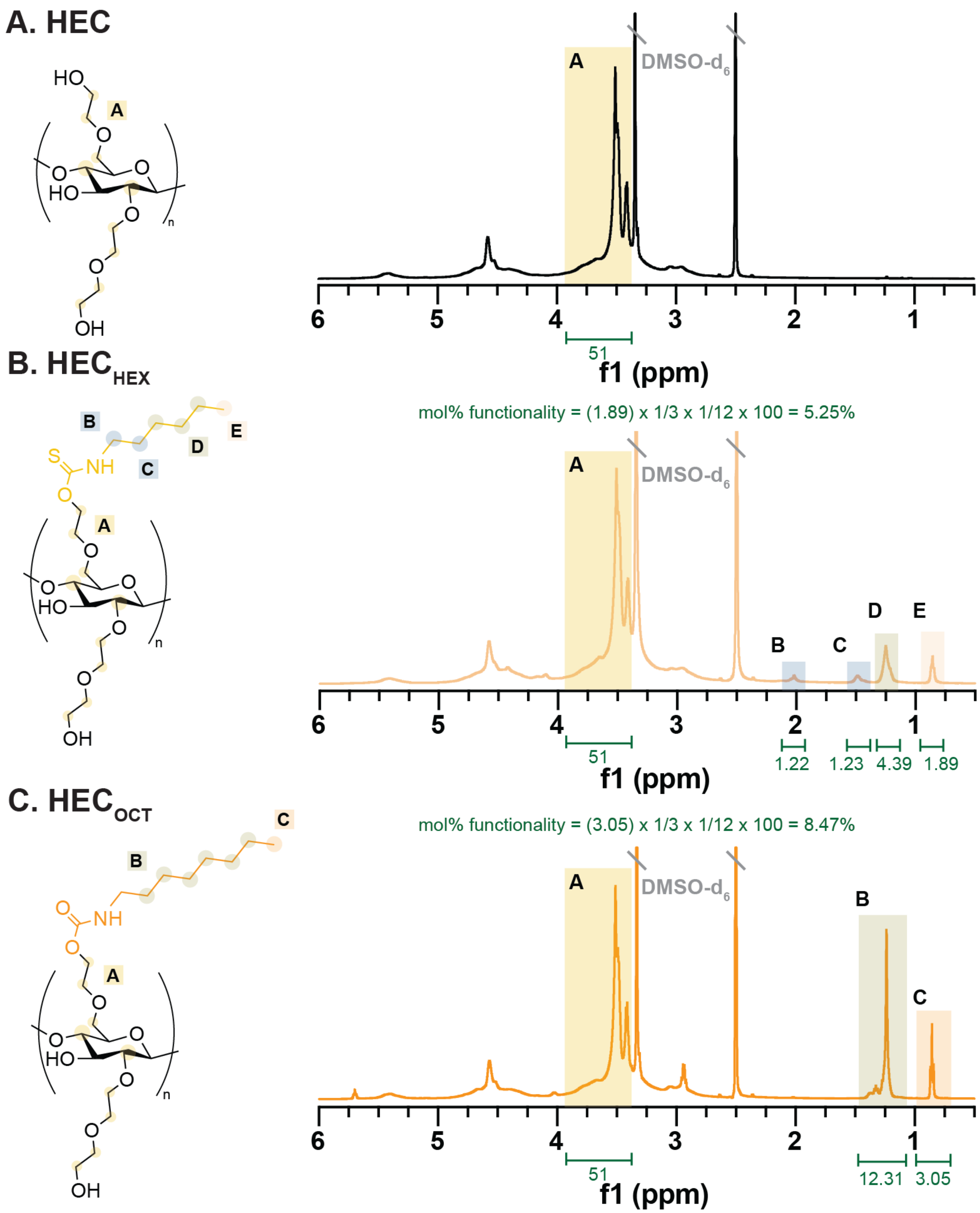


**Figure S11: Detailed NMR characterization of HEC$_{HEX}$ and HEC$_{OCT}$. (A)** $^1$H NMR of HEC in dDMSO. **(B)** $^1$H NMR of HEC$_{HEX}$ in dDMSO with integration normalized to conserved HEC peaks and highlighting HEX-associated peaks. **(C)** $^1$H NMR of HEC$_{OCT}$ in dDMSO with integration normalized to conserved HEC peaks and highlighting OCT-associated peaks.

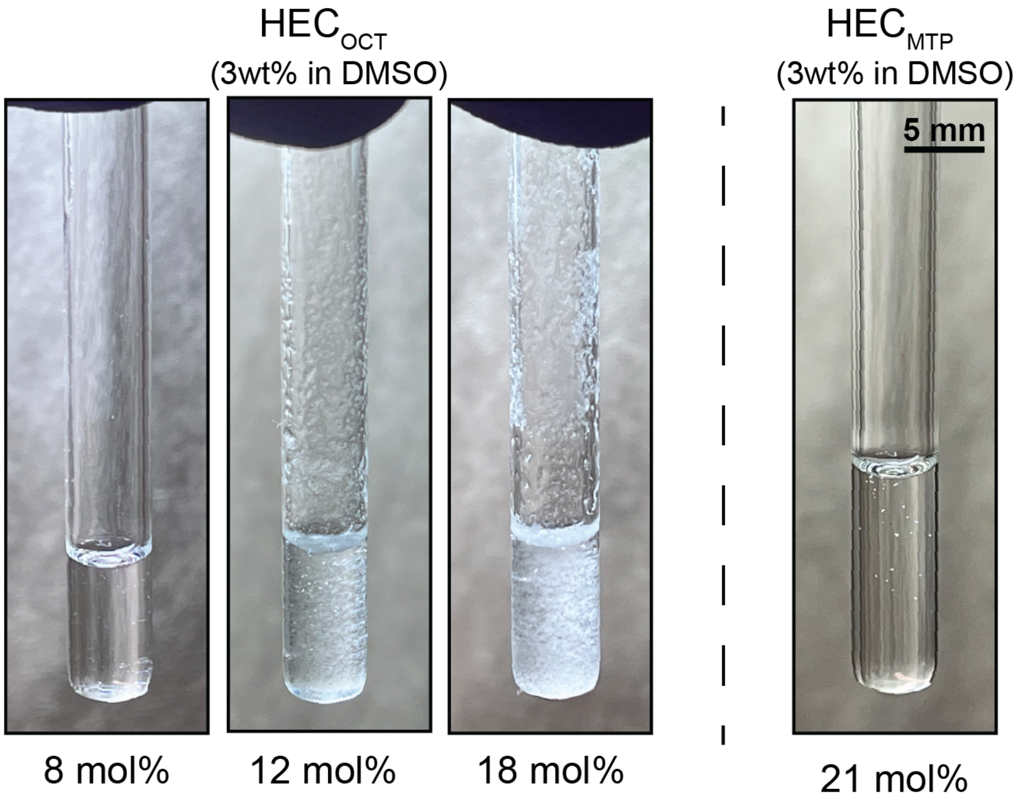


**Figure S12: HEC$_{OCT}$ DMSO solubility limits coating applications.** Brightfield images show HEC$_{OCT}$ with 8 mol%, 12 mol%, and 18 mol% functionality and HEC$_{MTP}$ at 21 mol% functionality solubilized at 3wt% in DMSO. 12 mol% and 18 mol% HEC$_{OCT}$ readily goes into solution at 50°C but quickly crystallizes out on returning to room temperature, while 8 mol% remains stable. Meanwhile, HEC$_{MTP}$ is stable in solution at functionalities as high as 21 mol% at a concentration of 3wt% in DMSO.

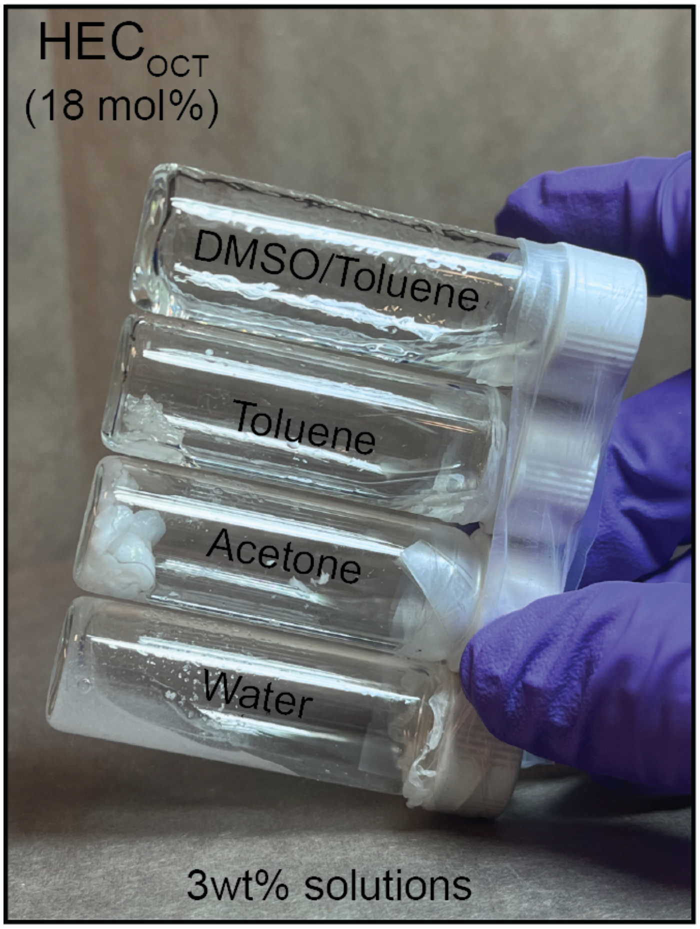


**Figure S13: HEC$_{OCT}$ is not readily soluble in common organic solvents.** 18mol% HEC$_{OCT}$ was solubilized at 3wt% in water, acetone, toluene, and 50/50 DMSO/toluene. This material forms a hydrogel in water, is insoluble in acetone and toluene, and is partially soluble in 50/50 DMSO/toluene as evidenced by the viscous/gel-like patterning seen above.

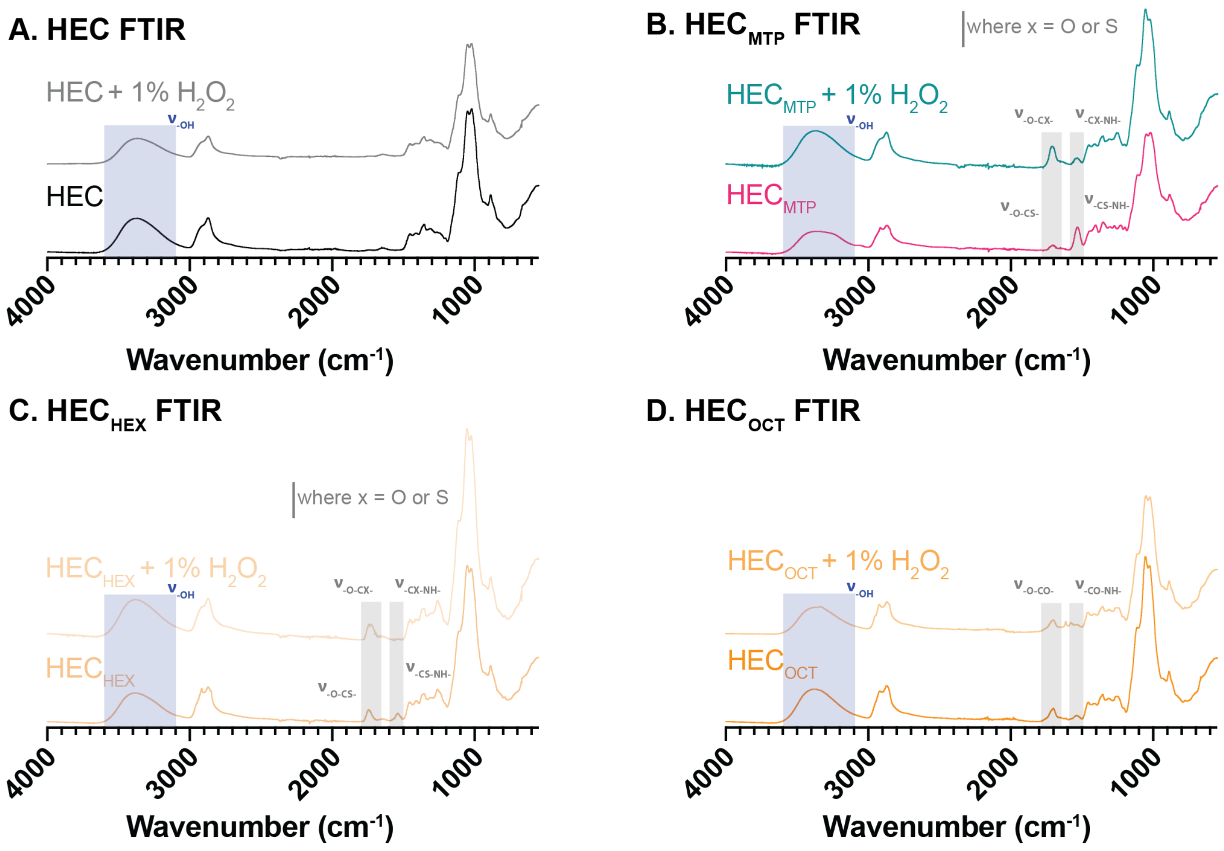


**Figure S14: Oxidation of HEC, $HEC_{MTP}$, $HEC_{HEX}$, and $HEC_{OCT}$ with 1% $H_2O_2$ demonstrates no significant loss of MTP, HEX, or OCT functional groups.** FTIR comparing **(A)** HEC **(B)** $HEC_{MTP}$ **(C)** $HEC_{HEX}$ and **(D)** $HEC_{OCT}$ both untreated and treated with 1% $H_2O_2$ for 24hrs. Highlights show either -CS-NH- and -O-CS- (thiocarbamoyl) or -CO-NH- and -O-CO- (carbamate) stretches.

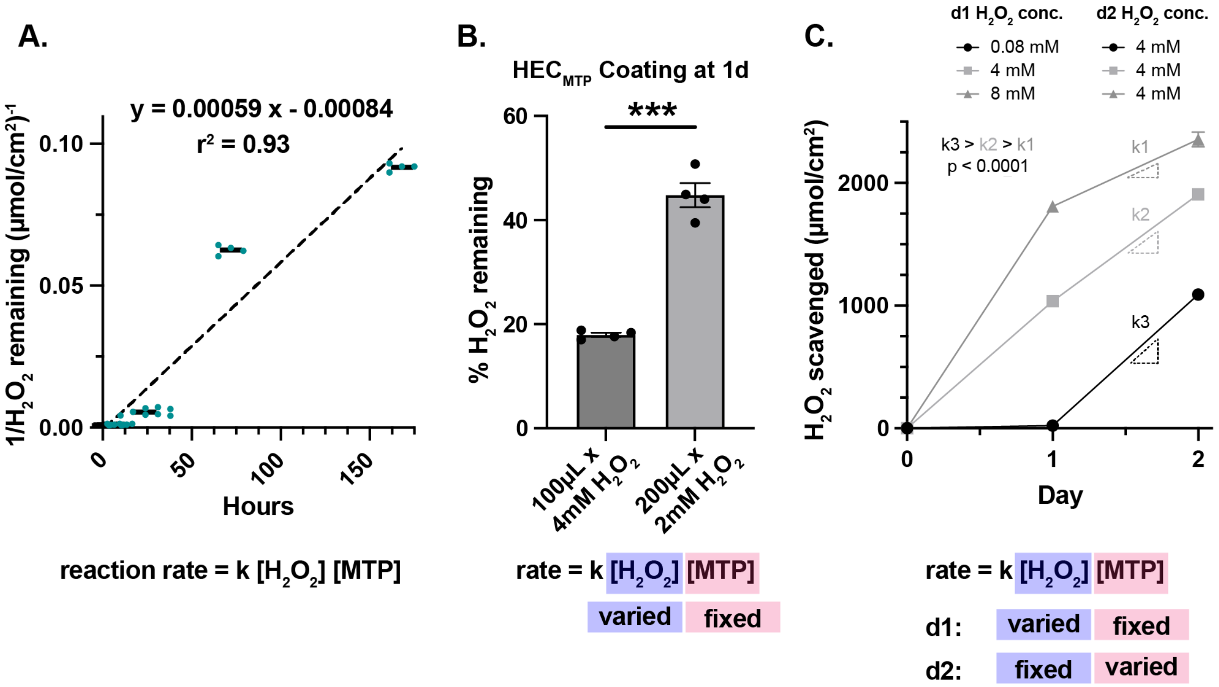


**Figure S15: $H_2O_2$ scavenging displays second order kinetics. (A)** Graph of $H_2O_2$ scavenging kinetics replotted such that the y-axis is 1/concentration. This shows that the reaction demonstrates 2$^{nd}$ order kinetics. k = 0.00059. **(B)** %$H_2O_2$ remaining after 1d of $H_2O_2$ reacting with $HEC_{MTP}$ coating. Red and blue highlights show the varied or fixed variable in the reaction rate equation. ***P< 0.0001, Student's t-test. Graph shows mean ± s.e.m. with individual data points showing n = 4 per group. **(C)** $H_2O_2$ scavenged (μmol/cm$^2$) by $HEC_{MTP}$ coating exposed to varied concentration of $H_2O_2$ on d1 and then a fixed concentration of $H_2O_2$ (4mM) on d2. Concentration of MTP moieties is fixed on d1, but following exposure to varied concentration of $H_2O_2$ on d1, there is a varied molar quantity of unreacted MTP moieties between conditions, creating a varied MTP concentration on d2. Dashed lines indicate the slope of the line between d1 and d2. Red and blue highlights show the varied or fixed variable in the reaction rate equation across d1 and d2. Slopes were compared using a two-way ANOVA with Tukey's multiple comparison test. Graph shows mean ± s.e.m. with individual data points showing n = 4 per group.

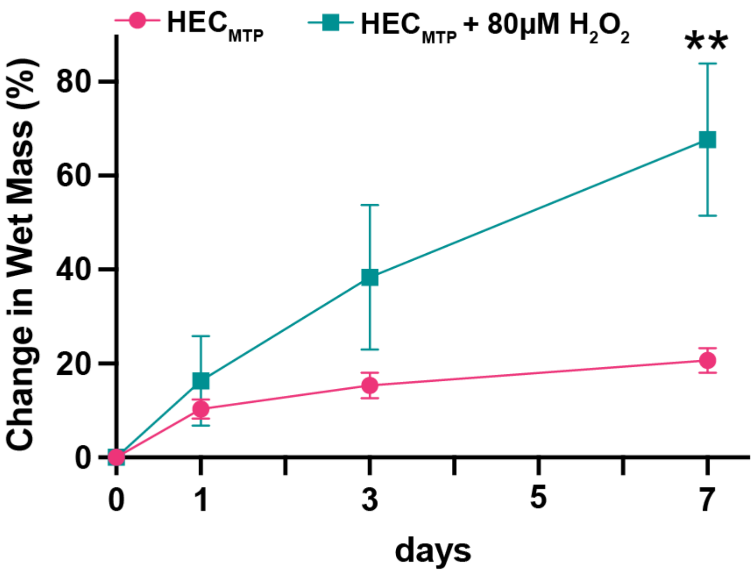


**Figure S16: Oxidized HEC$_{MTP}$ coatings exhibit increased swelling.** Change in wet mass of HEC$_{MTP}$ coatings over 7d exposed to either PBS or PBS + 80µM $H_2O_2$. **P= 0.0016 comparisons between HEC$_{MTP}$ and HEC$_{MTP}$ + 80µM $H_2O_2$ at equivalent timepoints, two-way ANOVA with Tukey's multiple comparison test. Graph shows mean ± s.e.m., n = 3 per group.

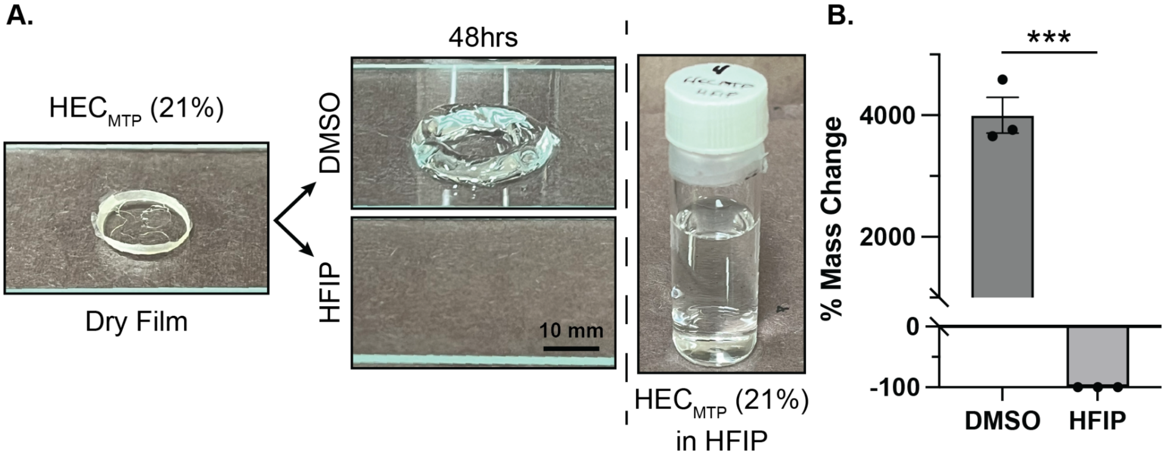


**Figure S17: HEC$_{MTP}$ films exhibit strong hydrogen bonding. (A)** Brightfield images showing a dry HEC$_{MTP}$ film that swells significantly in DMSO over 48 hrs, but completely dissolves in HFIP (a solvent that is excellent at disrupting hydrogen bonding). **(B)** %Mass change of HEC$_{MTP}$ films swollen in DMSO relative to HEC$_{MTP}$ films that were dissolved in HFIP. ***P=0.0002, Student's t-test. Graph shows mean ± s.e.m., n = 3 per group.

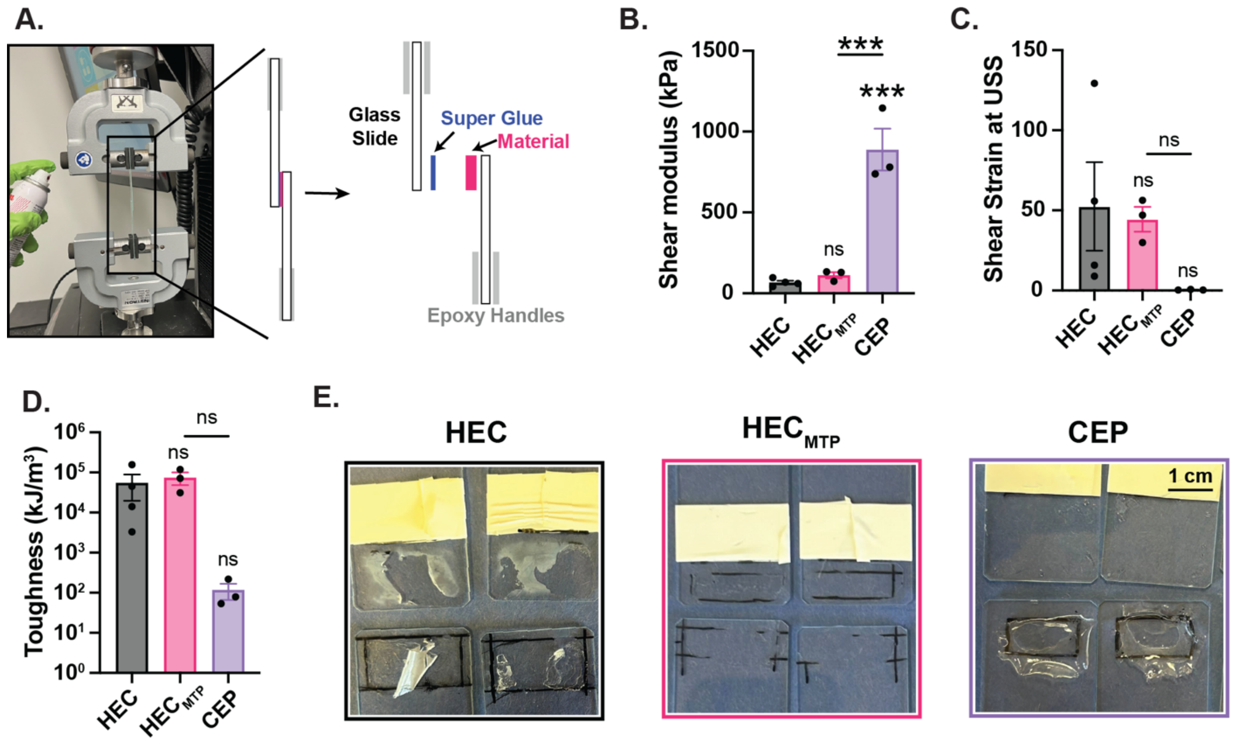


**Figure S18: Coating delamination testing set-up and results. (A)** Image and diagram demonstrating the assembly of lap shear slides. **(B)** Comparison of HEC, $HEC_{MTP}$, and crosslinked ethoxylated polyol (CEP) shear modulus. **(C)** Comparison of HEC, $HEC_{MTP}$, and CEP shear strain at the ultimate shear stress (USS). **(D)** Comparison of HEC, $HEC_{MTP}$, and CEP toughness as determined by taking the area under the curve (AUC) of the shear stress-shear strain curve. **(E)** Comparison of HEC, $HEC_{MTP}$, and CEP failure mechanisms demonstrating that HEC-based materials fail by both adhesive and cohesive failure, while CEP demonstrates pure adhesive failure.

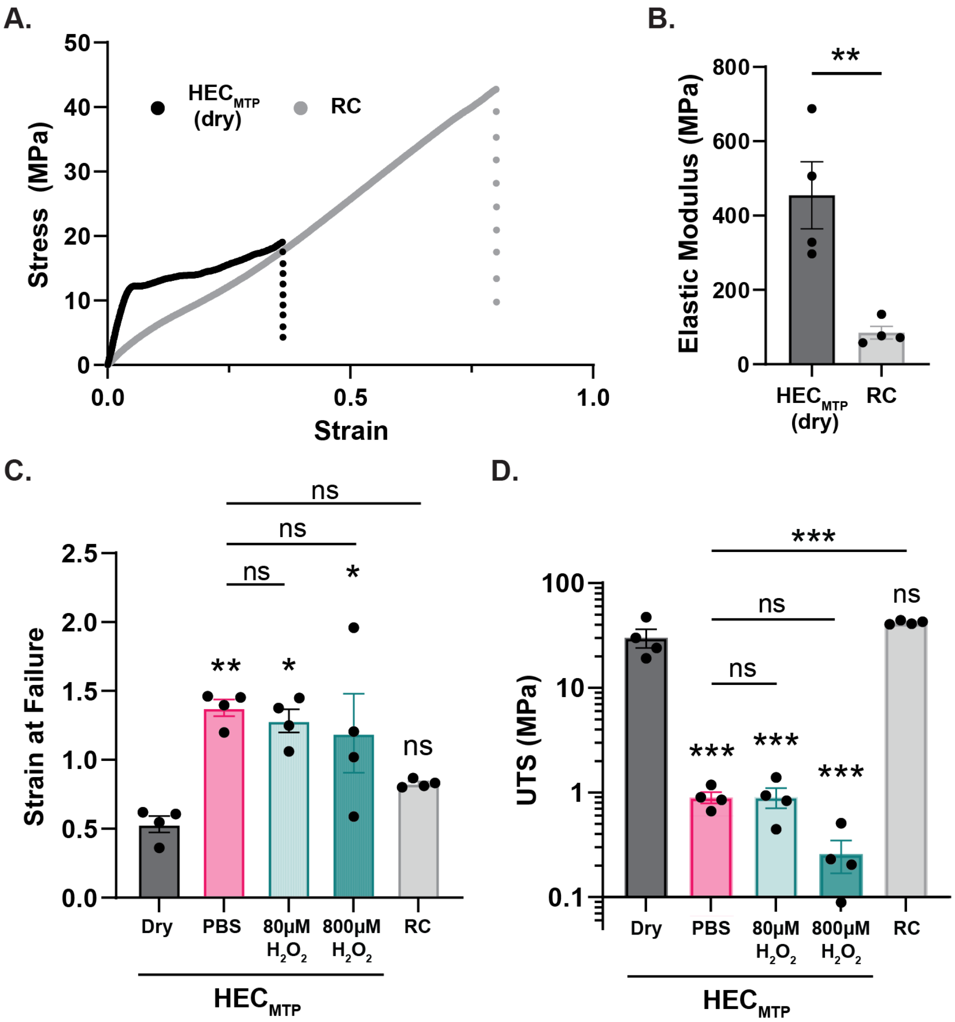


**Figure S19: Bulk material properties by tensile testing. (A)** Graph showing representative trace of tensile testing for dry HEC$_{MTP}$ films and hydrated (7d in PBS) regenerated cellulose (RC). **(B)** Bar graph showing the elastic moduli of dry HEC$_{MTP}$ films and hydrated (7d in PBS) RC. **$P<0.007$, Student's t-test. Graph shows mean ± s.e.m. with individual data points showing n = 4. **(C)** Bar graph showing the strain at failure for films of dry HEC$_{MTP}$, hydrated HEC$_{MTP}$ (7d in PBS), HEC$_{MTP}$ hydrated 7d in PBS + 80μM $H_2O_2$, HEC$_{MTP}$ hydrated 4d in PBS + 800μM $H_2O_2$, and hydrated RC (7d in PBS). *$P\leq0.03$, **$P\leq0.0049$, and not significant (ns) compared to dry HEC$_{MTP}$ or as designated, one-way ANOVA with Tukey's multiple comparison test. Graph shows mean ± s.e.m. with individual data points showing n = 4. **(D)** Bar graph showing the ultimate tensile strength (UTS) of films of dry HEC$_{MTP}$, hydrated HEC$_{MTP}$ (7d in PBS), HEC$_{MTP}$ hydrated 7d in PBS + 80μM $H_2O_2$, HEC$_{MTP}$ hydrated 4d in PBS + 800μM $H_2O_2$, and hydrated RC (7d in PBS). ***$P<0.0001$ and not significant (ns) compared to dry HEC$_{MTP}$ or as designated, one-way ANOVA with Tukey's multiple comparison test. Graph shows mean ± s.e.m. with individual data points showing n = 4.

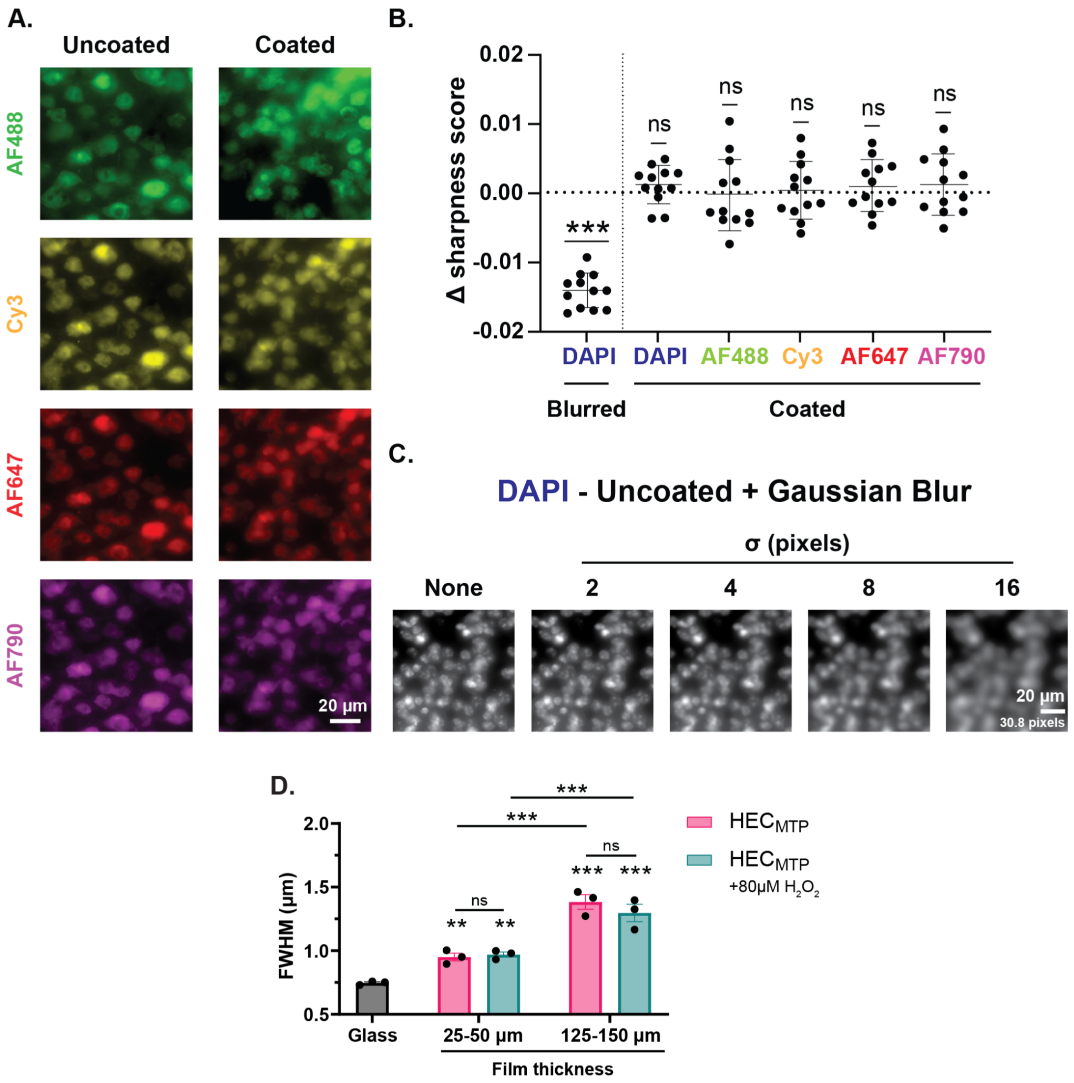


**Figure S20: $HEC_{MTP}$ coatings exhibit minimal fluorescent scattering. (A)** Representative widefield fluorescence images of stained mouse brain sections under $HEC_{MTP}$-coated and uncoated coverslips. **(B)** Graph showing the change in sharpness score in the $HEC_{MTP}$-coated vs uncoated images. Not significant (ns) and ***P <0.0001 compared to 0, one sample t-test. Graph shows mean ± s.d with individual data points showing the difference between n = 12 coated ROIs and the mean of n = 12 uncoated ROIs for all groups. **(C)** Images showing a characteristic ROI from a DAPI-stained mouse brain section under an uncoated coverslip after application of Gaussian blurring filters of increasing size. **(D)** Graph showing the average apparent size of sub-resolution fluorescent beads for an uncoated glass coverslip or a glass coverslip with a press fit film of either $HEC_{MTP}$ or $HEC_{MTP}$ + 80µM $H_2O_2$ for 7d. Data is binned by film thickness ranging from 25-50µm or 125-150µm. Apparent bead size was measured by the full width at half maximum (FWHM) of a 1D Gaussian fit. Not significant (ns), **P< 0.01, and ***P≤ 0.0002, compared to glass or as designated, two-way ANOVA with Tukey's multiple comparison test. Bar graph shows mean ± s.e.m. with individual data points showing n = 3 for all groups.

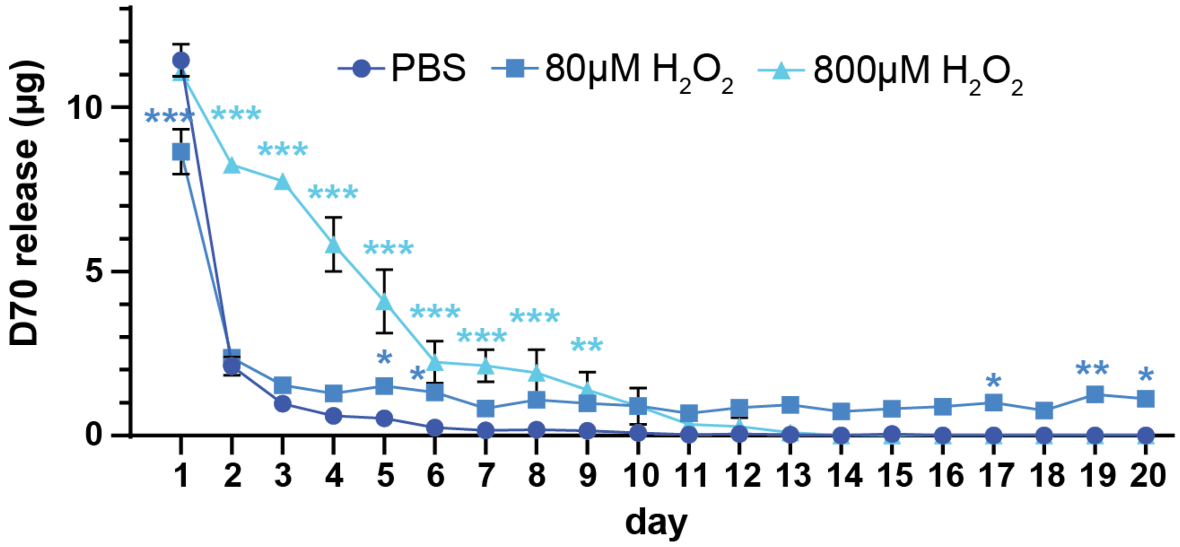


**Figure S21: HEC$_{MTP}$ film daily release of D70.** Graph showing 20d daily release for 70kDa FITC-Dextran (D70) from HEC$_{MTP}$ films into PBS, PBS + 80µM $H_2O_2$, or PBS + 800µM $H_2O_2$. All statistical comparisons are made relative to the PBS control at a given time point (ie. PBS vs PBS + 80µM $H_2O_2$ at day 5) and color-coded for clarity. *$P<0.04$, **$P<0.007$, ***$P\leq0.0001$, two-way ANOVA with Tukey's multiple comparison test. Graph shows mean ± s.e.m. with individual data points showing n = 3 per group.

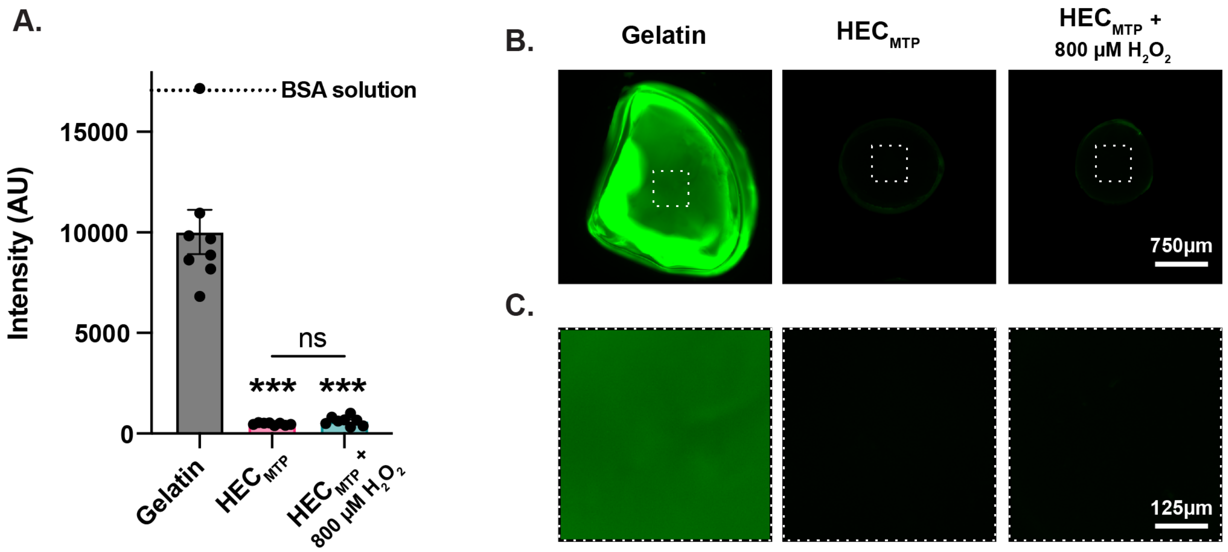


**Figure S22: $HEC_{MTP}$ coatings exhibit minimal protein adhesion. (A)** Plot showing the relative fluorescent intensity of albumin-fluorescein isothiocyanate conjugate (FITC-BSA) adsorbed to gelatin, $HEC_{MTP}$, and $HEC_{MTP}$ + 800µM $H_2O_2$ (1d) materials. Not significant (ns) and ***P< 0.0001 compared to gelatin or as designated, one-way ANOVA with Tukey's multiple comparison test. Bar graph shows mean ± s.e.m. with individual data points showing n = 8 for all groups. **(B)** Overview of gelatin, $HEC_{MTP}$, and $HEC_{MTP}$ + 800µM $H_2O_2$ (1d) materials with adsorbed FITC-BSA. Dashed box indicates representative ROI from which intensity measurements were taken. **(C)** Detail images showing the specified ROI in (B).

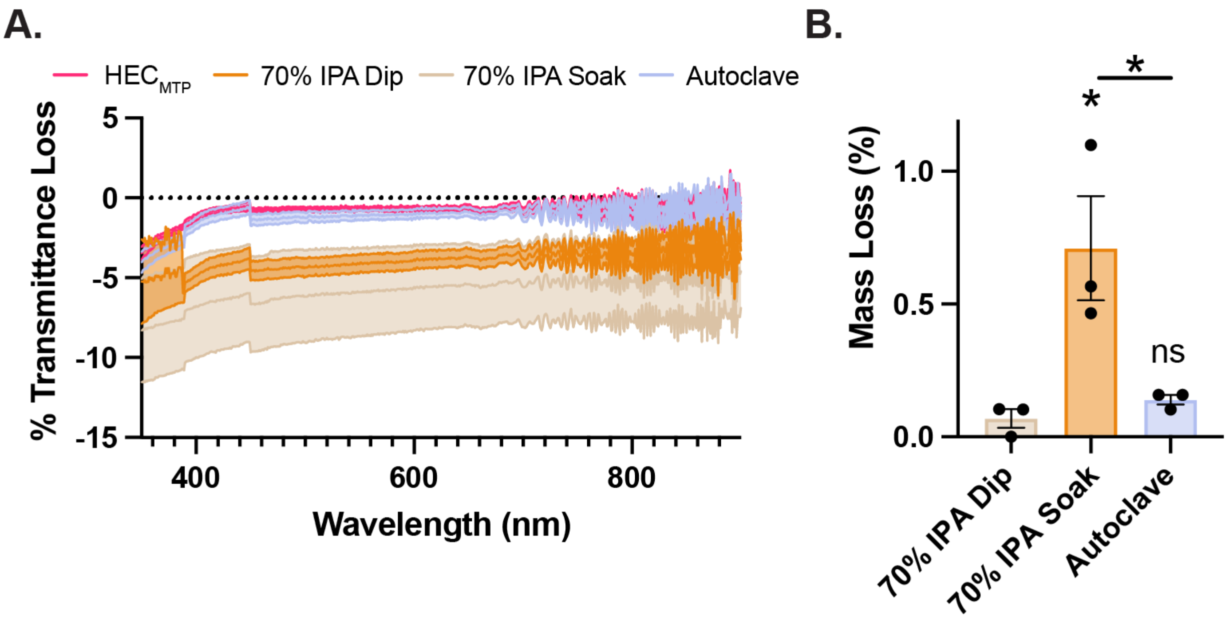


**Figure S23: HEC$_{MTP}$ coatings can be readily sterilized with minimal changes in mass loss and material properties. (A)** UV-Vis spectra of HEC$_{MTP}$ coatings on glass coverslips with no treatment, IPA sterilization with a single dip, as well as soaking overnight, and autoclaved. Spectra are background subtracted from a blank glass coverslip spectra. Graphs show mean ± s.e.m., with s.e.m. represented as light shaded areas. **(B)** %Mass loss for HEC$_{MTP}$ coatings with IPA sterilization with a single dip, as well as soaking overnight, and autoclaved. Not significant (ns) and *P ≤0.03 compared to 70% IPA dip or as designated, one-way ANOVA with Tukey's multiple comparison test. Graph shows mean ± s.e.m. with individual data points showing n = 3 for all groups.

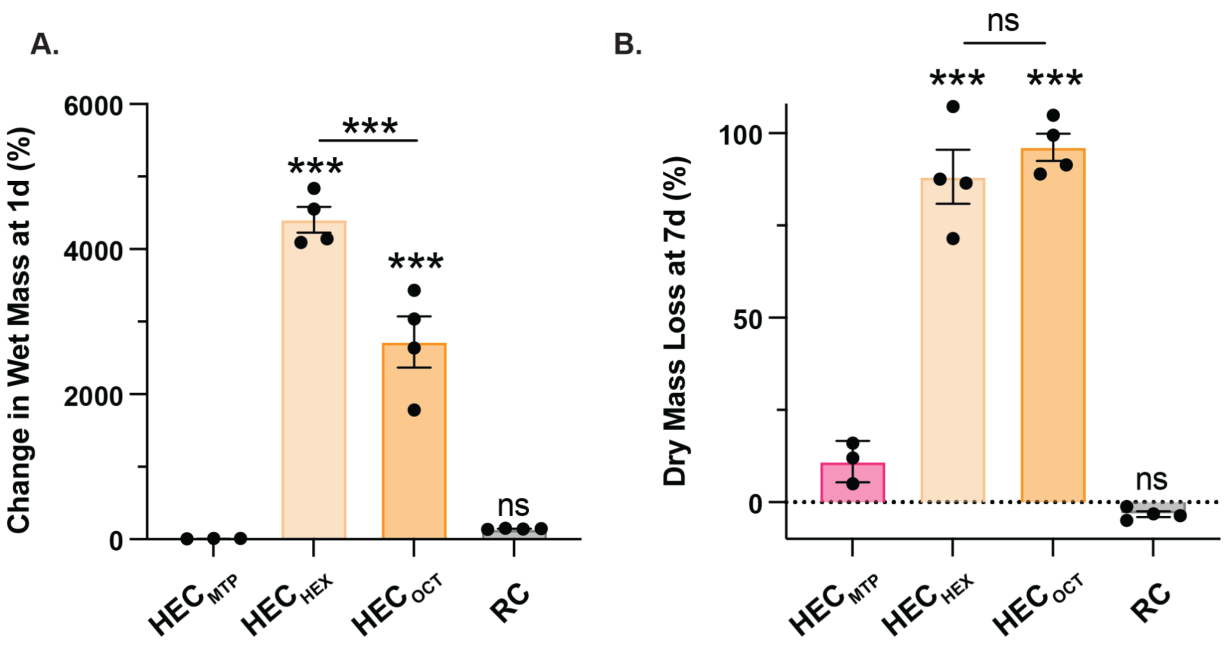


**Figure S24: Characterization of swelling and dissolution of implanted biomaterials. (A)** Change in wet mass of $HEC_{MTP}$, $HEC_{HEX}$, and $HEC_{OCT}$ and RC after 1d in PBS. Not significant (ns), and ***$P \leq 0.0006$ compared to $HEC_{MTP}$ or as designated, one-way ANOVA with Tukey's multiple comparison test. Graph shows mean ± s.e.m. with individual data points showing n = 3 for $HEC_{MTP}$ and n = 4 for all other groups. **(B)** %Dry mass loss of $HEC_{MTP}$, $HEC_{HEX}$, $HEC_{OCT}$, and RC after 7d in PBS. Not significant (ns), and ***$P < 0.0001$ compared to $HEC_{MTP}$ or as designated, one-way ANOVA with Tukey's multiple comparison test. Graph shows mean ± s.e.m. with individual data points showing n = 3 for $HEC_{MTP}$ and n = 4 for all other groups.

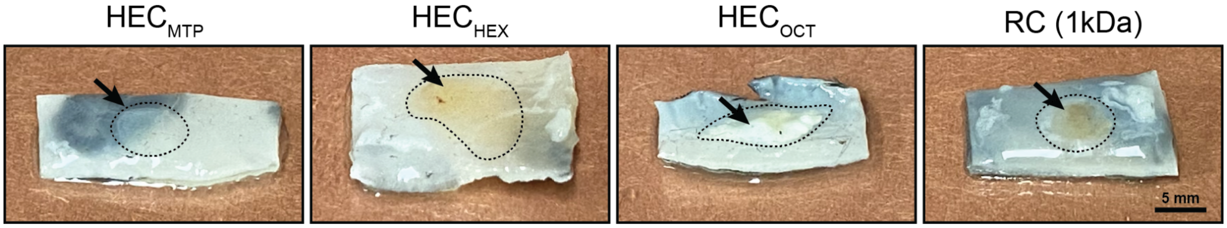


**Figure S25: Images of cellulose-based biomaterial explants after 4 weeks of subcutaneous implantation.** Arrows and dashed outlines indicate either the implanted material ($HEC_{MTP}$, RC) or the foreign body response to the implanted material ($HEC_{HEX}$, $HEC_{OCT}$).

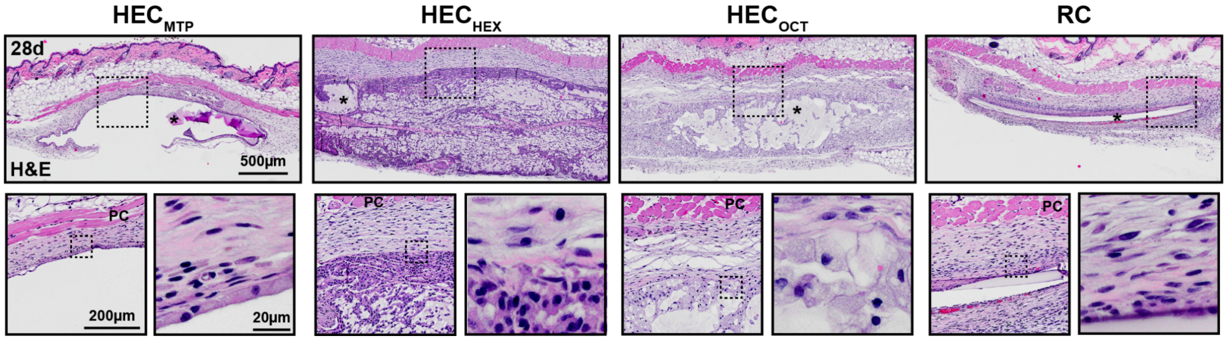


**Figure S26: H&E staining of subcutaneous implant tissue.** Survey and detail images of histology sections stained with hematoxylin and eosin (H&E) showing implant site and foreign body response (FBR) to subcutaneously implanted $HEC_{MTP}$, $HEC_{HEX}$, $HEC_{OCT}$, and regenerated cellulose (RC) films after 28d.

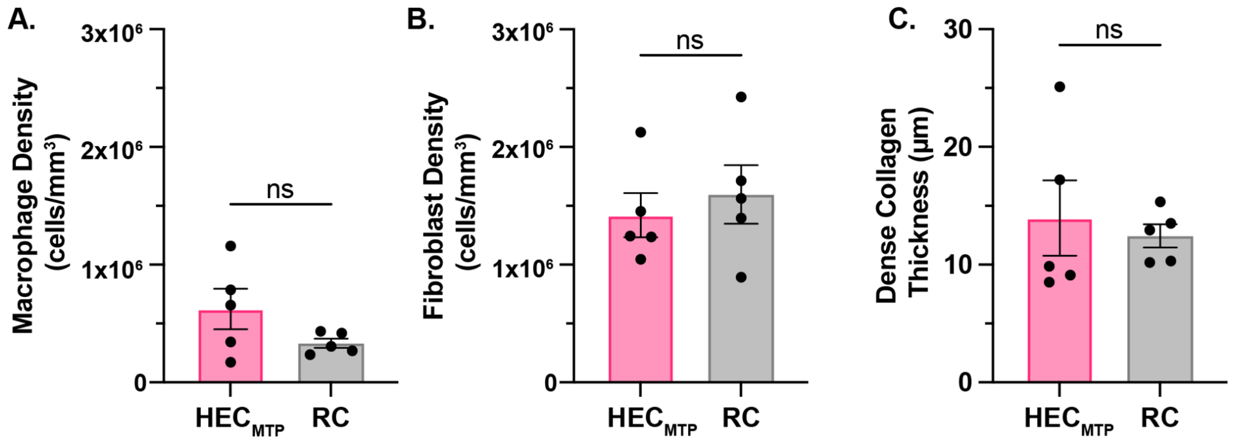


**Figure S27: Histological analysis of subcutaneous implant tissue reveals no significant difference in $HEC_{MTP}$ and RC FBR.** Plots comparing **(A)** macrophage density **(B)** fibroblast density and (C) dense collagen thickness between $HEC_{MTP}$ and RC materials. Not significant (ns), Student's t-test. Graph shows mean ± s.e.m. with n = 5 per group.
